# An anatomical and day–night atlas links cochlear immunity to adjacent bone marrow

**DOI:** 10.64898/2026.10.31.763358

**Authors:** Han Wang, Joe Yeong, Yao Cai, Yekang Mo, Ye Tian, Weibang Chen, Dexiao Zeng, Hongyu Ning, Yue Xu, Shuangjie Dai, Xiaoyu Liu, Ruoyun He, Huafeng Liu, Junmin Huang, Junfeng Hao, Yusu Ni, Chunjie Tian

**Affiliations:** Department of Otorhinolaryngology, Affiliated Hospital of Guangdong Medical University, Zhanjiang 524000, Guangdong, China; Guangdong Medical University, Zhanjiang 524000, Guangdong, China; Guangdong Provincial Key Laboratory of Autophagy and Major Chronic Non-communicable Diseases, Zhanjiang 524001, Guangdong, China; Key Laboratory of Prevention and Management of Chronic Kidney Disease of Zhanjiang City, Affiliated Hospital of Guangdong Medical University, Zhanjiang 524001, Guangdong, China; Institute of Nephrology, Affiliated Hospital of Guangdong Medical University, Zhanjiang 524001, Guangdong, China; Department of Family Medicine, Shengjing Hospital of China Medical University, Shenyang 110022, Liaoning, China; ENT Institute and Department of Otorhinolaryngology, Eye & ENT Hospital, Fudan University, Shanghai 200031, China; NHC Key Laboratory of Hearing Medicine (Fudan University), Shanghai 200031, China; Otology and Skull Base Surgery, Eye & ENT Hospital, Fudan University, Shanghai 200031, China; Otolaryngology Research Center, Guangdong Medical University Affiliated Hospital, Zhanjiang 524000, Guangdong, China; Institute of Molecular and Cell Biology (IMCB), Agency for Science Technology and Research (A*STAR), Singapore, Singapore; Department of Anatomical Pathology, Singapore General Hospital, Singapore, Singapore; Singapore Immunology Network (SIgN), Agency for Science Technology and Research (A*STAR), Singapore, Singapore; Cancer Science Institute, National University of Singapore, Singapore, Singapore

## Abstract

The cochlea lies within marrow-bearing bone. How this neighbouring marrow relates to cochlear immunity and daily physiology remains poorly resolved. We profiled 160,643 cells from mouse cochlea and adjacent temporal-bone marrow at six times of day, with rhythm inference calibrated for one pooled sample per time. Three-dimensional confocal reconstruction mapped vascular connections between marrow and cochlea and localized neutrophils along this interface. Neutrophils also occupied healthy cochlear fluids. Parabiosis findings were consistent with a local marrow contribution to cochlear immunity. Structural cochlear cells maintained synchronized, light-entrained clocks. Cochlear neutrophil programmes aligned with circulating neutrophils and opposed those in marrow, with independent flow cytometry supporting daily changes in immune composition and CXCR4. A shared night-associated heat-shock programme was attenuated after noise in an independent cohort. These anatomical and temporal data provide a reference for testing local marrow supply and the origins of cochlear immune rhythms.

---

The cochlea supports hearing within a fluid-filled bony enclosure close to haematopoietic marrow. Studies of its immune cells have focused on macrophage maintenance and responses to injury ^1–3^, while neutrophils have been described principally during inflammation^4,5^. Nearby marrow could contribute to this immune environment alongside recruitment from the systemic circulation. Establishing its contribution requires an anatomical map of the connection, evidence of cellular origin and measurements of cell states in both compartments. These observations would help explain how the sensory organ maintains and changes its immune population.

Skull and vertebral marrow supply meningeal myeloid cells, and skull marrow also supplies meningeal B cells through vascular channels ^6–8^. Cerebrospinal fluid can reach these niches and influence their activity^9,10^, and skull marrow differs molecularly from marrow in other bones ^11^. Vascular connections between temporal marrow and the labyrinth have also been mapped in the rat^12^. This anatomy suggests that marrow associated with the cochlear bony capsule could contribute to local immune maintenance. An anatomical connection, cellular origin and migration are distinct forms of evidence; assessing them together is needed to test whether the inner ear draws on an adjacent immune reservoir.

The cochlea has a local clock, and identical noise exposures can produce different hearing outcomes depending on time of day^13–15^. Leukocyte recruitment and marrow release also vary across the day^16–20^. Circulating neutrophils undergo daily changes in state, which tissues can select or modify ^21–23^. A tissue’s immune cell programme may therefore differ from both its structural-cell clocks and the programme of the marrow supplying it. Cell-type-resolved brain atlases have revealed temporal differences obscured in whole-tissue measurements ^24,25^, while cochlear atlases have described cellular responses to noise and ageing ^26–28^. Comparing cochlea and neighbouring marrow across the day allows anatomical proximity and possible cellular supply to be examined alongside temporal state.

We profiled mouse cochlea and adjacent temporal-bone marrow at six times of day, with rhythm inference calibrated for one pooled sample per tissue and time point. We combined this single-cell atlas with three-dimensional confocal reconstruction, cellular imaging, parabiosis and independent flow cytometry. The reconstructions map marrow, cochlear vessels and neutrophil locations, and descriptive parabiosis findings are consistent with a local marrow contribution. Cochlear structural cells retain synchronized, light-entrained clocks, whereas cochlear neutrophil programmes align with blood and oppose those in marrow. Independent datasets and pre-specified flow measurements test separate components of these findings. The atlas provides a resource for studying how a sensory organ and surrounding marrow organize local immunity, with observed relationships distinguished from hypotheses about cell movement and the origin of immune timing.

## Results

### A day–night atlas of the cochlea and adjacent temporal-bone marrow

We sampled the cochlea and adjacent temporal-bone marrow of perfused mice every 4 h across a 12-h light–dark cycle, at zeitgeber times (ZT) 3, 7, 11, 15, 19 and 23 (lights on at ZT0, 08:00), and profiled 160,643 cells by single-cell RNA sequencing: 116,278 from the cochlea and 44,365 from the marrow, in 35 subtypes of 12 classes (Fig. 1a–e, Extended Data Fig. 1 and Supplementary Table 1). Each tissue and time point is one pooled sample (*n* = 6 per tissue); rhythms were therefore tested against exact nulls over the 60 ways of assigning the samples to times (time relabelling), phases by leaving out one time point at a time, and claims graded on pre-specified evidence (Methods). Subtypes intermixed across samples, least so neutrophil states, whose programmes change over the day (Extended Data Fig. 1d).

**Fig. 1.**
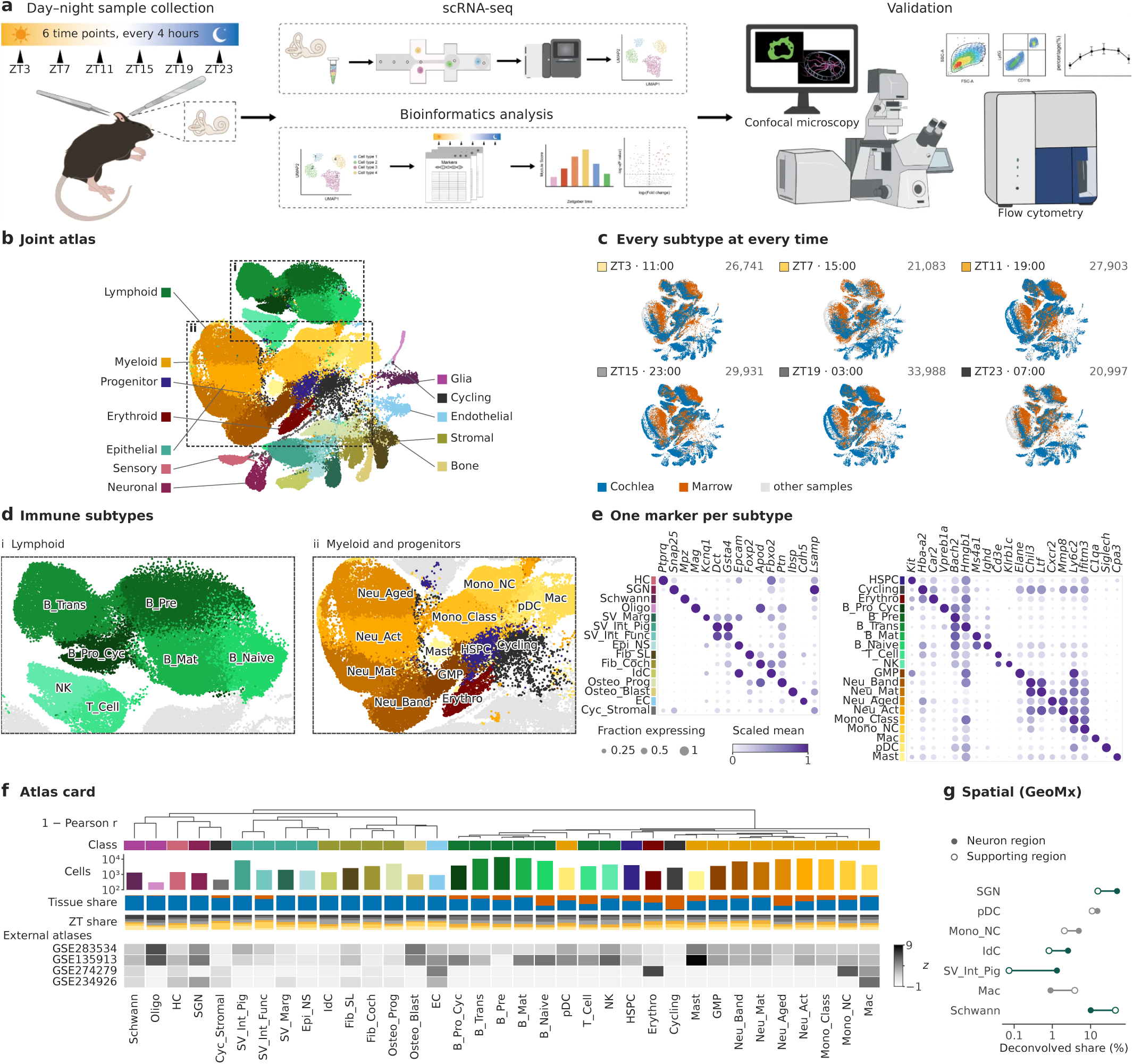
A day–night atlas of the cochlea and adjacent temporal-bone marrow. **a**, Study design: cochlea and temporal-bone marrow of perfused mice collected every 4 h across a 12-h light–dark cycle (ZT0, lights on at 08:00; clock time, ZT + 8 h), one pooled sample per tissue and time (*n* = 6 per tissue), profiled by single-cell RNA sequencing and validated by imaging and flow cytometry. **b**, Joint UMAP embedding of all 160,643 cells coloured by the 35 subtypes (class hue, subtype lightness); dashed boxes, areas enlarged in d. **c**, Cells of each time point on the joint embedding, coloured by tissue (grey, all other cells); numbers, cells per time point. **d**, Enlargements of b: i, lymphoid subtypes; ii, myeloid subtypes and progenitors. **e**, One marker gene per subtype (structural, left; haematopoietic, right): dot area, fraction of cells expressing; colour, mean expression scaled per gene within each compartment. **f**, Atlas card, subtypes ordered by average-linkage clustering of pseudobulk profiles (distance, 1 *− r*). Tracks: class; cells (log scale); tissue share; ZT share; *z*-score of each subtype’s markers in the cells assigned to it in four independent cochlear atlases (blank, *<* 10 cells). **g**, Deconvolved shares in the neuron and supporting-cell regions of public cochlear spatial transcriptomics data (GSE312646, controls) for subtypes that differ between regions (two-sided Welch’s *t*-test across 16 and 14 areas of illumination; Benjamini–Hochberg *q <* 0.05). Subtype abbreviations, Supplementary Table 1.

After label transfer to four independent adult cochlear atlases ^27,29–31^, cells assigned our labels showed higher expression of the expected markers for 77–97% of labels. Deconvolution of public cochlear spatial transcriptomes^32^ placed spiral ganglion neurons (0.49 versus 0.15 of the deconvolved signal in neuron versus supporting-cell regions; *q* = 2.3 *×* 10*^−^*^14^), Schwann cells and macrophages in their expected regions (Fig. 1f,g and Extended Data Figs. 2 and 3). To calibrate detection under the sampling design, we subsampled dense held-out time series^14,33^ to one sample at each of six times. Our strict (Tier 1) and lenient (Tier 2) rhythm calls (Methods) kept the realized false-discovery proportion at 3.4–5.5% and 7.6–13.6%, against 31.6–36.8% for nominal *P <* 0.05, at the cost of sensitivity (3–12% for Tier 1; Extended Data Fig. 4). The rules favoured specificity over sensitivity; failure to detect a rhythm therefore does not establish arrhythmicity. The annotated atlas, rhythm calls and cell-type summaries are released as a resource (Supplementary Tables 2–5).

### Cochlear structural cells keep synchronized, light-entrained clocks

All 14 structural cell types of the cochlea expressed core-clock genes with high amplitude and in their canonical order ^34^, as did the marrow, at the light-entrained phase of reference organs^33^ (Fig. 2a and Extended Data Fig. 5a–c). The cochlear clock lay +0.55 h (interval 0.18–0.92 h) and the marrow clock +0.71 h from the canonical phase, close to independent cochlear series after sham surgery (*−*0.42 h) or adrenalectomy (*−*0.05 h) ^14^ and to an independent marrow series (+0.12 h) ^35^ (Fig. 2b), and clocks fitted to these reference series estimated our sampling times to within about 1–2.5 h (Extended Data Fig. 5d). This phase alignment is consistent with the established dependence of the cochlear clock on central light entrainment^14^.

**Fig. 2.**
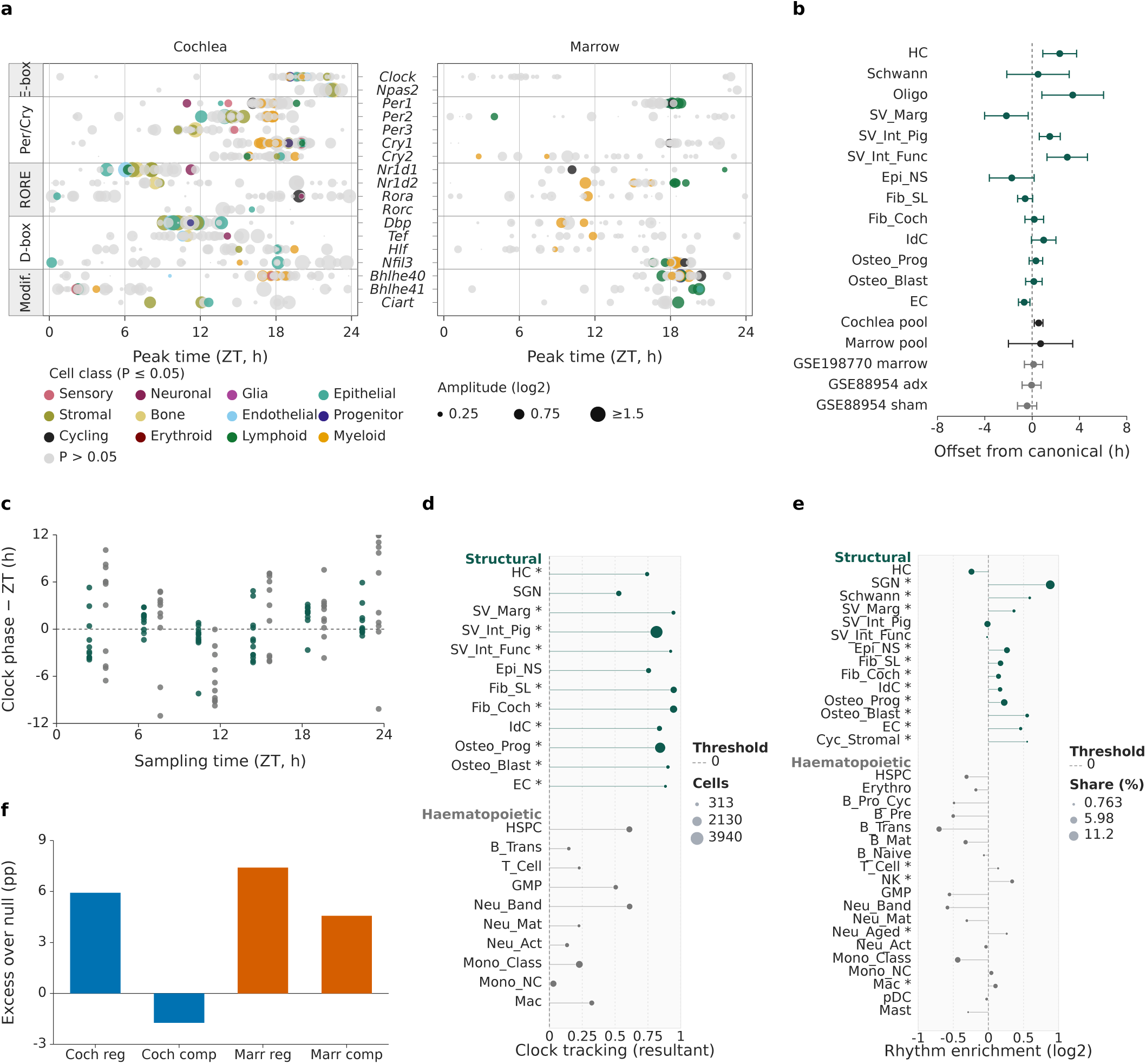
Cochlear structural cells keep synchronized, light-entrained clocks. **a**, Peak time of 18 core-clock genes (*Arntl* not detected) in every cell type of cochlea and marrow, grouped by regulatory arm. Colour, cell class when rhythmic (exact time-relabelling *P ≤* 0.05; grey otherwise); dot area, amplitude (log_2_). **b**, Offset of each cochlear structural cell type’s clock from the canonical clock phase (bars, 95% gene-bootstrap confidence interval), with all cochlear and all marrow cells (pools) and independent time series of cochlea (GSE88954; sham and adrenalectomy (adx)) and marrow (GSE198770, control). **c**, Mean single-cell clock phase minus sampling time for every cell type and sample (structural, green; haematopoietic, grey). **d**, Clock tracking per cell type: resultant length of the single-cell clock offsets across samples; asterisk, exact time-relabelling *P ≤* 0.05; dot area, cells. **e**, Contribution of each cell type to the rhythmic genes of the whole cochlea (GSE88954, sham): log_2_ ratio of its mean share of these genes’ counts to its share of expression-matched non-rhythmic genes; asterisk, bootstrap *P ≤* 0.05; dot area, mean share. **f**, Rhythms of whole cochlea (Coch; GSE88954, sham) and marrow (Marr; GSE198770, control) reproduced by within-cell regulation (reg) or by composition (comp): excess over a gene-label permutation null, in percentage points (pp). *n* = 6 pooled samples per tissue.

**Fig. 3.**
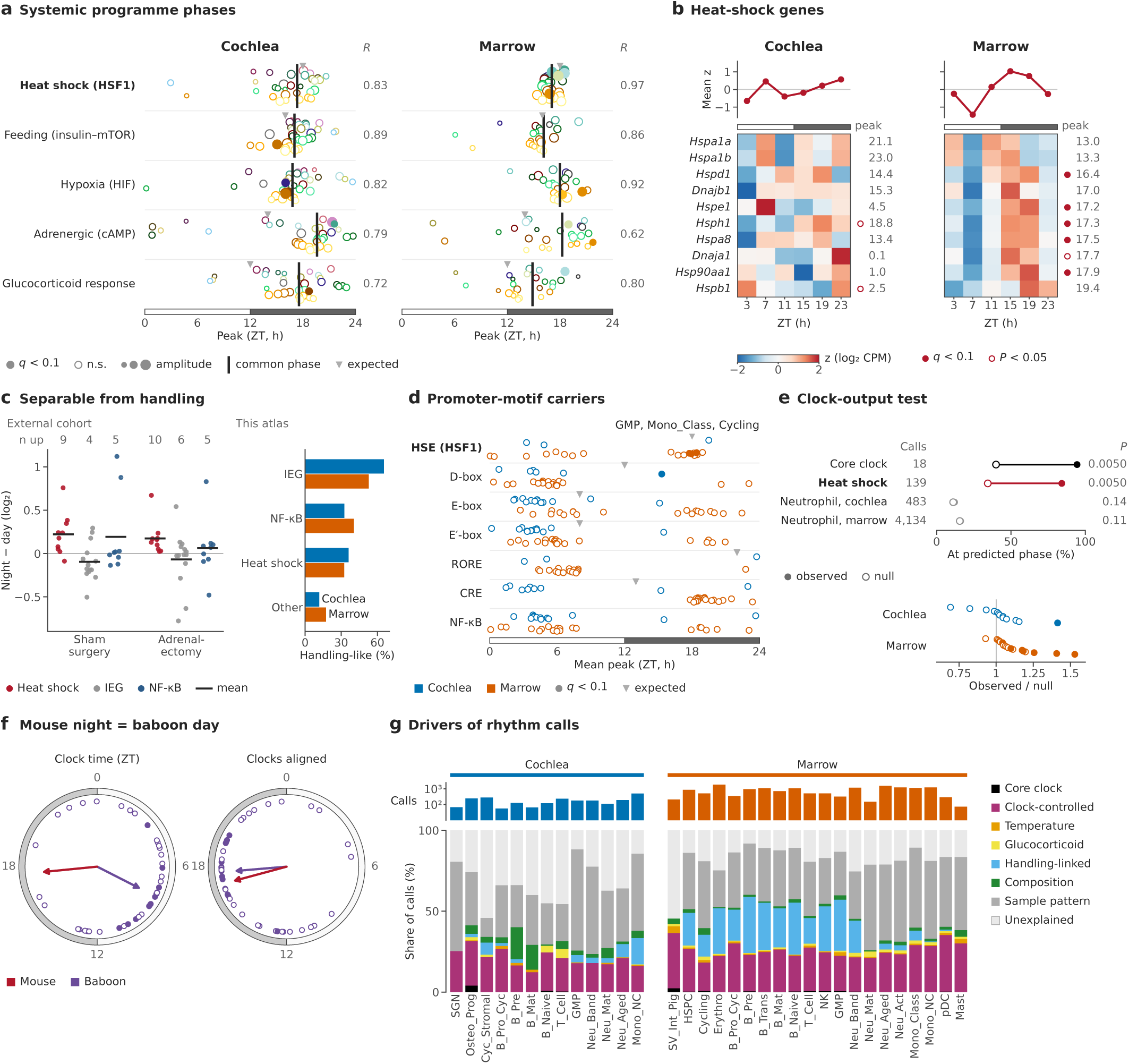
A systemic night heat-shock programme follows the activity phase. **a**, Peak phase of five systemic-cue gene programmes in every cell type of cochlea and marrow; *R*, resultant length of the amplitude-weighted common phase. Cell types share the six samples, so the convergence is descriptive. **b**, Ten heat-shock genes in the whole-tissue pseudobulk (*z*-scored log_2_ CPM; ordered by marrow peak); top, programme mean; right, cosinor peak. **c**, Left, night*−*day change (about ZT17.5 versus ZT5.5) of heat-shock, immediate-early (IEG) and NF-*κ*B genes in an independent cochlear series without noise (GSE107086), after sham surgery or adrenalectomy; bar, mean; *n* up, genes with a positive change. Right, share of our rhythm calls whose per-sample profile follows the dissociation-stress (handling) profile. **d**, Circular mean peak of the rhythmic carriers of each promoter motif per cell type (filled, directional permutation *q <* 0.1); labelled, marrow cell types with significant heat-shock element (HSE) carriers. **e**, Top, share of rhythm calls peaking at the phase predicted from constant darkness (GSE54650) versus a phase-permuted null, with calls and permutation *P*. Bottom, observed/null ratio per cell type (filled, *P <* 0.05). **f**, Heat-shock programme phase in mouse (78 rhythm calls) and baboon (GSE98965^40^; dots, 45 tissues; filled, *P <* 0.05) on each species’ clock time and after aligning the clocks; arrows, circular mean (length, resultant length). **g**, Primary driver of each Tier-1/2 rhythm call per cell type (types with *≥* 20 calls). *n* = 6 pooled samples per tissue; Tier-1/2, calibrated rhythm calls (Methods).

Clock phase also followed the time of day within single structural cells: it tracked sampling time in 10 of 12 cochlear structural types (exact time-relabelling *P ≤* 0.05; Fig. 2c,d) but in none of 10 cochlear haematopoietic types, which carry few clock transcripts; among cells with 3–5 clock transcripts, 9 of 12 structural and none of the haematopoietic types did (Extended Data Fig. 5g). Daily immune cell states were therefore assessed through transcriptional programmes rather than inferred single-cell clock phase.

Within-cell-type regulation accounted for part of the rhythms measured in whole cochlea. Genes rhythmic in an independent whole-cochlea time series^14^ drew their counts disproportionately from neural, glial, vascular, bone and strial cells (Fig. 2e). We separated the contributions of expression and cellular composition by rebuilding each gene’s tissue profile with either composition or expression held fixed: regulation within cell types reproduced 20.8% of the 2,845 rhythms, against 14.8% expected under a gene-label permutation (+5.9 percentage points; *P* = 0.0050), whereas changing composition reproduced 6.7%, against 8.4% (*P* = 1.0; Fig. 2f). In the marrow, by contrast, both mechanisms contributed (+7.4 and +4.6 points; *P* = 0.0050 each). Consistent with regulation inside cells, transcription-factor activity lagged the factors’ own mRNA by about 1 h in the cochlea (+1.1 h; Rayleigh *P* = 4.3 *×* 10*^−^*^9^; Extended Data Fig. 6d).

### A systemic heat-shock programme rises at night in cochlea and marrow

Five systemic cues can influence peripheral clocks in addition to light^36–39^: heat shock, feeding, hypoxia, adrenergic and glucocorticoid signalling. Across cell types in both tissues, programmes associated with these cues peaked in the active (dark) phase, between ZT14.9 and ZT19.6 (Fig. 3a). The heat-shock programme had the most concentrated peak phases, peaking at ZT17.1 in the marrow (resultant length 0.97) and at ZT17.3 in the cochlea (0.83). As cell types share their tissue’s six samples, this convergence is descriptive; we tested the heat-shock programme with four independent lines of evidence.

First, independent data and technical checks argued against a simple handling-related explanation. In whole-tissue pseudobulk, 5 of 10 heat-shock genes were rhythmic in the marrow (*q <* 0.1; peaks ZT16.4–17.9), although none reached this threshold in the cochlea (Fig. 3b). In an independent cochlear cohort without noise exposure ^14^, heat-shock genes were higher at night after sham surgery (+0.22 log_2_; 9 of 10 genes up; *P* = 0.0019) and after adrenalectomy (+0.17; 10 of 10; *P <* 0.001), whereas immediate-early genes were not (Fig. 3c); in our atlas, the per-sample dissociation-stress effect was not rhythmic (*P* = 0.11 and 0.12; Extended Data Fig. 7a). Second, heat-shock elements in promoters carried the night phase: in marrow cell types, their rhythmic carriers peaked at ZT17.8–18.4 (*q ≤* 0.033; Fig. 3d). Third, like core-clock genes and unlike the neutrophil-state programmes, heat-shock rhythms peaked where constant-darkness data predict (84% of calls versus 34% under a phase-permuted null; core clock, 94% versus 40%), suggesting a clock output (Fig. 3e). Fourth, in a diurnal primate^40^ the programme peaked by day (ZT7.8; 14 tissues with *P <* 0.05), at the same internal clock phase as in the nocturnal mouse once the clocks were aligned (17.7 versus 17.0 h; Fig. 3f): this cross-species alignment was consistent with activity phase rather than the external light phase.

Among all rhythm calls, the largest driver class was the tissue-wide sample pattern (36% in both tissues), followed by clock control (25% in marrow, 20% in cochlea; Fig. 3g; annotations, not causal tests); the heat-shock evidence above supports a night-associated programme shared by both tissues.

### Three-dimensional mapping of the cochlea and its marrow connection

To place the temporal atlas in its anatomical setting, we reconstructed the cochlea within the surrounding temporal bone. Haematopoietic cells made up 60–83% of the cochlear cells recovered for sequencing (Extended Data Fig. 1f), a proportion that describes the recovered preparation. In three-dimensional confocal reconstructions, vessels ran from the marrow onto the cochlea and branched over its surface (Fig. 4a); their hand tracing followed vessel ridges detected independently in the unannotated renders for 91% and 84% of its length, against 29% and 45% for random placements (*P* = 0.001 each; Supplementary Fig. 1). Neutrophils lay along vessels beside the cochlea (Fig. 4b), and spatial transcriptomics resolved neutrophil states within the tissue (Fig. 4c). Neutrophils also occupied the fluid spaces of the healthy cochlea: FUT4 marked 23 of 23 nucleated perilymph and 56 of 64 endolymph cells (marrow, 29 of 30), most of them CD45^+^ (Fig. 4d).

**Fig. 4.**
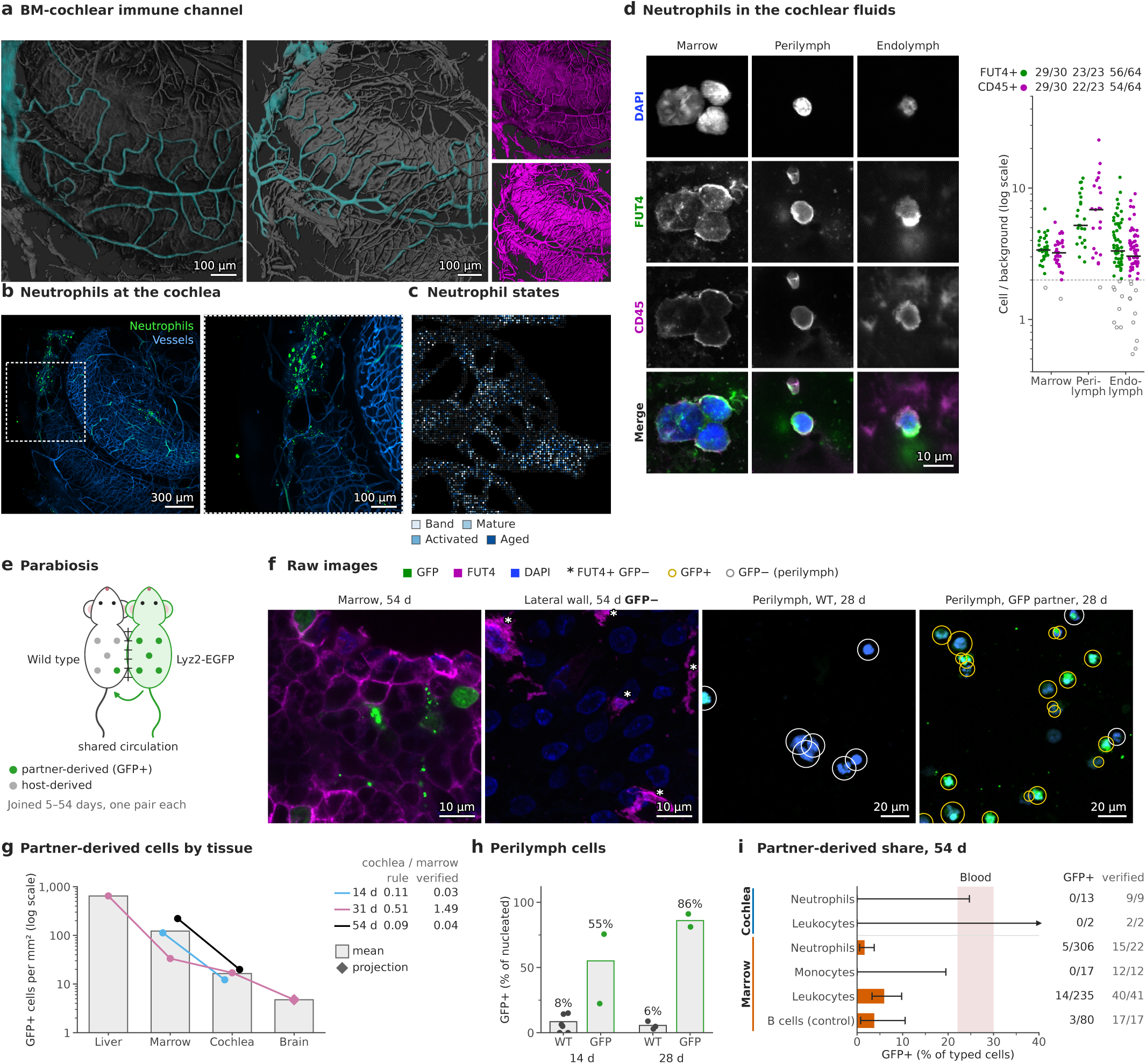
Three-dimensional anatomy and cellular-origin evidence at the cochlea–marrow interface. **a**, Vessels connecting temporal-bone marrow (BM) and the cochlea, traced (teal) on three-dimensional confocal renders; right, untraced. **b**, Neutrophils (green) along vessels (blue) beside the cochlea; dashed box, enlarged at right. **c**, Spatial transcriptomics: deconvolved neutrophil state per spot. **d**, Wild-type marrow, perilymph and endolymph cells; right, FUT4 and CD45 of each cell relative to its local background (filled circles, at least 2; bar, median; numbers, positive/all cells; 9, 10 and 9 fields). CD45 intensity is not comparable between preparations. **e**, Parabiosis design; in a wild-type partner, GFP^+^ cells are partner-derived. **f**, Wild-type partners: 54-day marrow and lateral wall; 28-day perilymph of both partners (one GFP display range). **g**, Partner-derived cells per mm^2^ of tissue, all tissues counted in the format of the brain image, on local background and by a stricter cell rule (Supplementary Fig. 5): bars, mean of pairs; lines, pairs (liver and brain, 31-day partner; diamond, brain projection); log scale; right, cochlea-to-marrow ratio by this count and corrected with blinded visual labels (verified). **h**, Perilymph GFP^+^ share of nucleated cells (bars, pooled; points, fields; 6, 2, 3 and 2 fields). **i**, Partner-derived share of immune cells typed on nuclei, 54-day wild-type partner (lines, Clopper–Pearson 95% CI; cochlear neutrophils from 60*×* sections; B cells, reporter-negative control; shaded, blood neutrophils in published parabiosis^7,41^); right, GFP^+^/typed cells and typed cells confirmed in a blinded review (verified). One pair per pairing duration; pairs are descriptive. Scale bars as labelled.

To assess labelling through a shared circulation, we joined wild-type mice to Lyz2-EGFP partners, whose myeloid cells carry GFP, so that GFP^+^ cells in a wild-type partner are partner-derived (one pair per pairing duration, 5–54 days; Fig. 4e,f and Supplementary Figs 2–4). Partner-derived cells were sparser in the cochlea than in the adjacent marrow: with the counting rule’s calls corrected by blinded visual review, the cochlea-to-marrow density ratio was 0.03 (95% confidence interval (CI) 0–0.08) at 14 days and 0.04 (0–0.14) at 54 days, whereas the 31-day pair was inconclusive (1.49, 0.67–4.39; Fig. 4g). Counted alike in the format of the only brain image, by a stricter rule set on the blinded labels, the 31-day partner held 646 partner-derived cells per mm^2^ in the liver, 33 in the marrow, 17 in the cochlea and 4.8 in the brain (Fig. 4g and Supplementary Fig. 5). Wild-type perilymph cells were 6–8% GFP^+^, against 55–86% in the GFP partners (Fig. 4h). In the 54-day partner, immune cells typed on their nuclei were rarely partner-derived even in the marrow (neutrophils, 5 of 306; reporter-negative B cells, a false-positive background, 3 of 80), and none of 13 cochlear neutrophils was GFP^+^ (Fig. 4i and Supplementary Fig. 6). Published parabiosis puts blood neutrophils at 22–30% partner-derived ^7,41^, above every share we measured in the cochlea (Fig. 4h,i). A single noise-exposed pair showed no excess of partner-derived cells (Supplementary Fig. 4), and the results held across the rules’ parameter settings (Supplementary Fig. 7). The sampled cochlear neutrophils were predominantly host-derived, as were those in adjacent marrow. Together with the vascular connection, these observations are consistent with a local marrow contribution, although they do not identify the source of individual host-derived cells.

### Cochlear immune composition varies across the day

Within the haematopoietic cells of the cochlea, myeloid cells dominated in the light (rest) phase (76% at ZT7) and lymphoid cells at night (51–56% at ZT15–19; Extended Data Fig. 1f). Cochlear band neutrophils peaked at ZT5.2 and classical monocytes at ZT7.0, and in the marrow naive and transitional B cells peaked at ZT20.5 and ZT12.8 (Fig. 5a,b; B-stage rhythms in Extended Data Fig. 8b). Each of these rhythms was supported by three to five core composition models (Methods) and kept its phase when single time points were left out (s.d. *≤* 0.4 h; Fig. 5a). A pre-specified flow-cytometry cohort of 20 mice tested these predictions at ZT5 and ZT17 (13:00 and 01:00). Cochlear immature (Ly6G^int^ CD11b^int^) neutrophils, the flow counterpart of band neutrophils, and classical monocytes were more abundant at ZT5 (ZT17/ZT5 ratio 0.67, 95% CI 0.52–0.85, Holm-adjusted *P* = 0.0025; and 0.40, 0.30–0.53, *P <* 0.001), and recirculating (naive) B cells of the marrow at ZT17 (1.68, 1.34–2.10, *P <* 0.001; Fig. 5c and Supplementary Fig. 8). All three primary predictions replicated and remained supported when gate thresholds were varied by 10%. The atlas cochleae differed 4.8-fold in their content of marrow-resident precursors, without a daily rhythm (*P* = 0.37); relative to these precursors, the timed shares kept their phases (shifts *≤* 0.9 h) with weaker *P* values, so we report unadjusted shares as primary (Extended Data Fig. 9g).

**Fig. 5.**
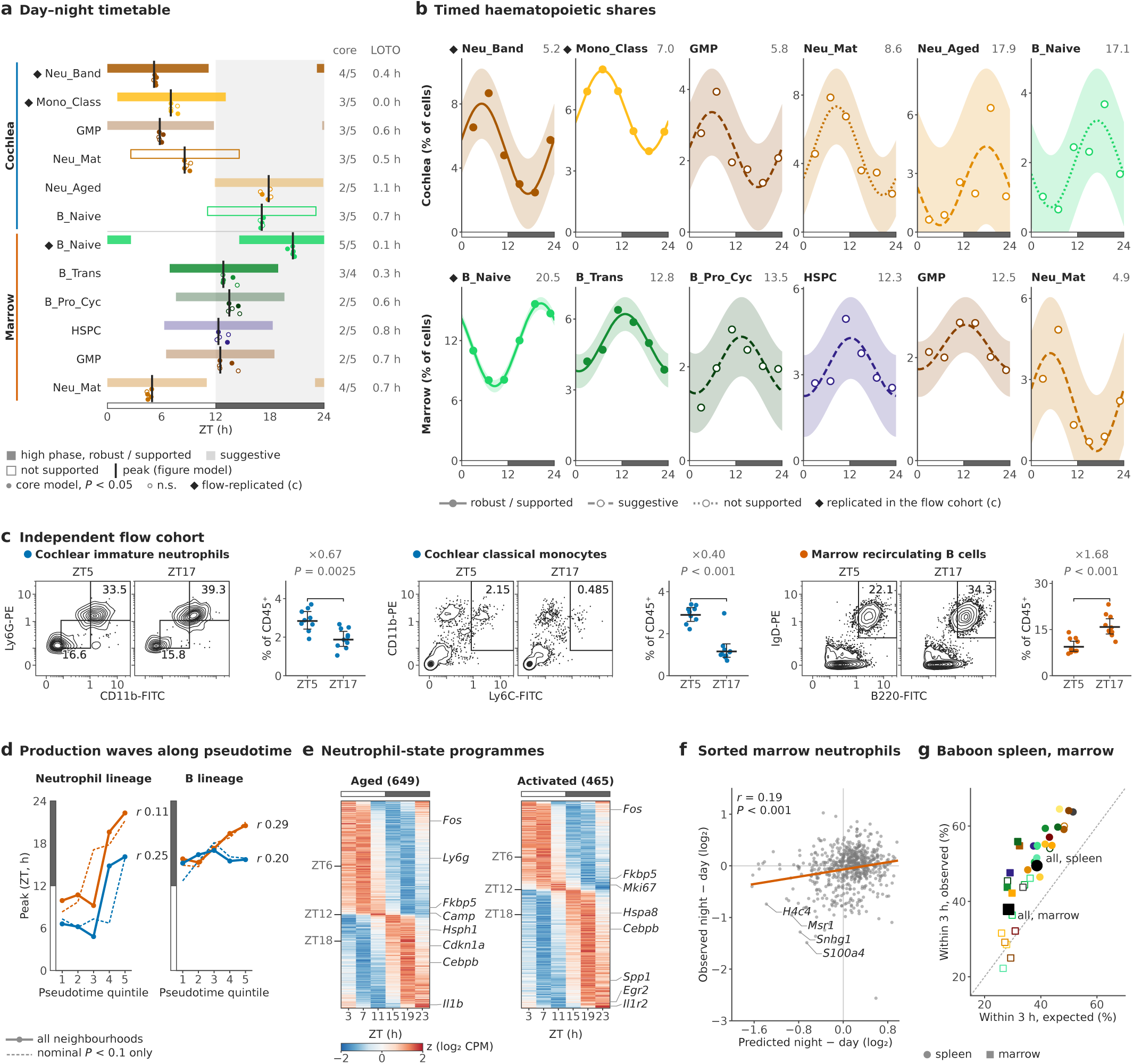
The cochlear immune compartment alternates over the day. **a**, Timetable of the timed haematopoietic shares of b: bars, phase in which the fit exceeds its mean (dark, robust or supported; light, suggestive; open, not supported); black ticks, peaks; dots, peaks in each of five composition models (filled, *P <* 0.05); core, supporting/tested models; LOTO, s.d. of leave-one-time-point-out peaks; diamonds, replicated in c; grey, dark phase. **b**, Shares of tissue cells: points, samples; lines, cosinor fit with 95% confidence band (solid, robust or supported; dashed, suggestive; dotted, not supported); header, peak (ZT). **c**, Pre-specified flow cytometry at ZT5 and ZT17: contour plots of the median mouse for each primary readout (numbers, % of the plotted population; gating, Supplementary Fig. 8) and % of CD45^+^ cells per mouse (bar, geometric mean; whiskers, 95% CI); ratio, ZT17/ZT5; *P*, Holm-adjusted, linear model of log % with time and cohort; *n* = 9 and 10 cochleae and 10 and 10 marrows (ZT5, ZT17). **d**, Circular mean peak of Milo neighbourhoods per pseudotime quintile (solid, all; dashed, nominal *P <* 0.1); *r*, circular–linear correlation. **e**, Tier-1 genes of aged and activated marrow neutrophils ordered by peak (*z*-scored log_2_ CPM). **f**, Sorted marrow neutrophils (GSE312298; ZT22 versus ZT14): observed versus predicted change; *r*, Pearson; *P*, gene-label permutation. **g**, Marrow rhythm genes within 3 h of the aligned baboon peak, observed versus expected (circles, spleen; squares, marrow; filled, *P <* 0.05; black, all genes). Blue, cochlea; vermilion, marrow; *n* = 6 pooled samples per tissue (a, b, d–g).

Several observations argue against marrow carryover as a complete explanation of the cochlear immune profile: neutrophils were present in cochlear fluids, cochlear classical monocytes peaked about 6 h before those of the marrow (ZT7.0 versus ZT12.7; Supplementary Table 5) and, as shown below, cochlear neutrophil programmes opposed those in marrow.

Later pseudotime neighbourhoods (label-free groups of similar cells) of the neutrophil and B lineages peaked later in the day (circular–linear *r* = 0.11 and 0.25 for neutrophils in marrow and cochlea, 0.29 for marrow B cells; permutation *P ≤* 0.0012; Fig. 5d), and 481 of 3,386 marrow neighbourhoods varied over the day (Extended Data Fig. 8a), although pre-specified tests of developmental order remained partial (Extended Data Figs. 8b–d and 9a). Aged and activated neutrophils (atlas labels; by a published maturation score^42^, ‘aged’ neutrophils are the most mature; Extended Data Fig. 9d) carried 1,114 of the 1,712 Tier-1 rhythm calls of the marrow (65%), forming day programmes (*Fos*, *Ly6g*, *Fkbp5*, *Camp*) and night programmes (heat-shock genes, *Cdkn1a*, *Cebpb*, *Il1b*, *Spp1*; Fig. 5e). These programmes predicted the day–night change of sorted marrow neutrophils in an independent study ^43^ (*r* = 0.19 over 644 genes; *P <* 0.001; Fig. 5f), and baboon immune tissues suggested partial conservation of their phases (within 3 h of ours for 50% versus 39% expected in spleen, *P* = 0.0010; Fig. 5g).

### Cochlear neutrophil programmes align with blood and oppose those in marrow

Circulating neutrophils undergo daily changes in transcription and surface phenotype associated with an intrinsic timer ^21,22^. We asked whether cochlear neutrophils shared this programme or instead resembled neutrophils in the adjacent marrow. Using the circulating-neutrophil programme as an independent reference, we compared, gene by gene, the change that each neutrophil state’s daily rhythm predicts between ZT5 and ZT13 with the change of circulating neutrophils between the same times^22^ (Extended Data Fig. 9e).

Across thousands of genes, predicted daily changes in cochlear neutrophils aligned with the blood programme, whereas those in marrow were inversely related. For aged neutrophils, the correlation was *ρ* = 0.35 in the cochlea, rising to 0.62 for the 63 genes with *P <* 0.05 in both data sets, and *ρ* = *−*0.33 in the marrow (*−*0.63 for 133 genes; Fig. 6a,b); removing stress and immediate-early genes left the split unchanged. It was specific to neutrophils: neutrophil states ranked first to third of 33 cochlear cell types (exact rank-sum *P* = 0.00018) and first, fourth and tenth of 21 marrow types (*P* = 0.040; Fig. 6c). A score of the blood programme peaked at ZT17.5 in cochlear neutrophils (*P* = 0.067) and at ZT4.0 in marrow neutrophils (*P* = 0.017; paired difference *P* = 0.050; Fig. 6d), and the split did not rest on neutrophil-identity genes (*ρ* = 0.25 in cochlea and *−*0.25 in marrow on genes not enriched in neutrophils; Fig. 6e).

**Fig. 6.**
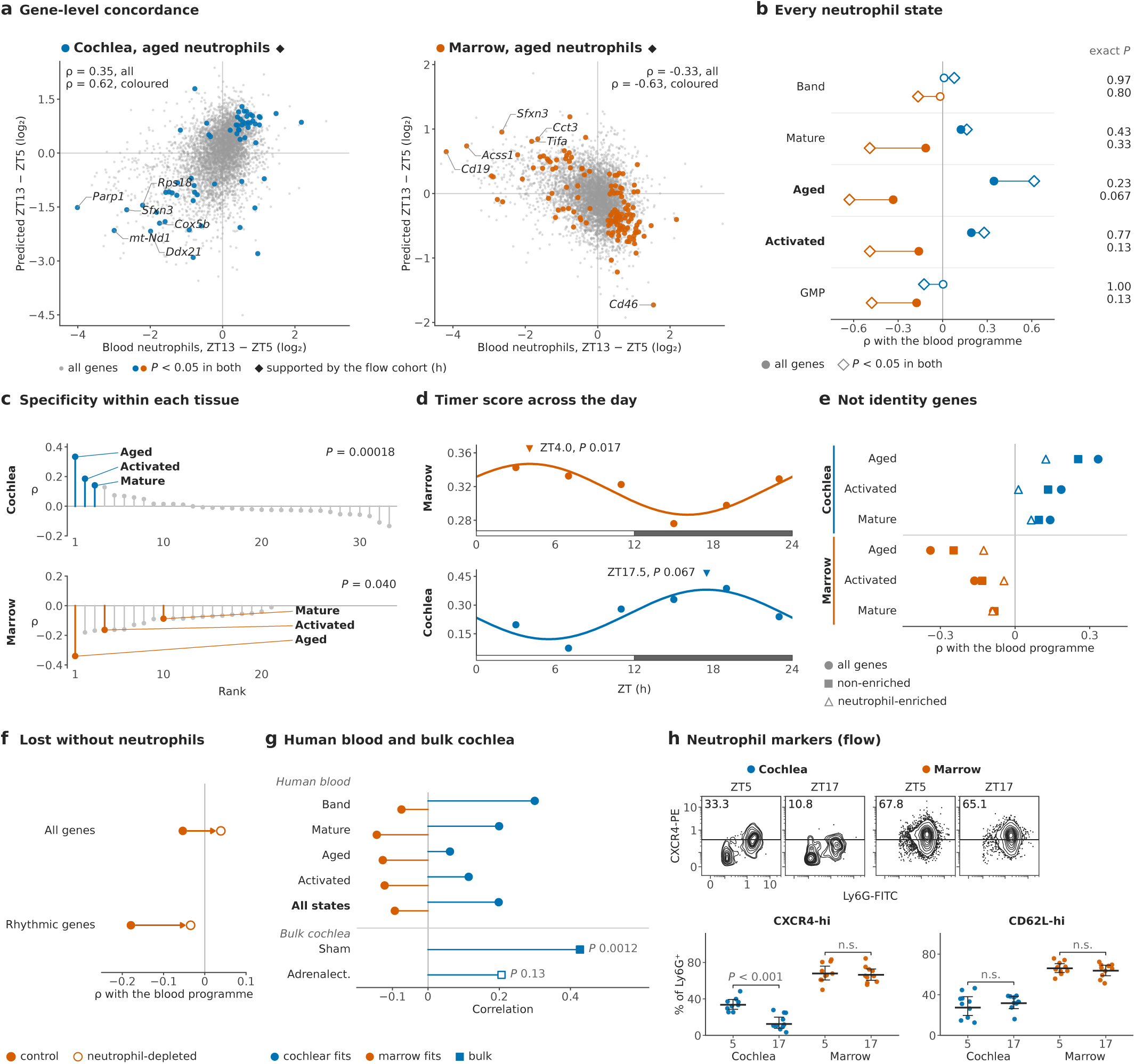
Cochlear neutrophil programmes align with blood and oppose those in marrow. **a**, Per gene, the ZT13*−*ZT5 change of blood neutrophils (GSE86619) versus that predicted for aged neutrophils of cochlea and marrow (grey, all shared genes; coloured, *P <* 0.05 in both); *ρ*, Spearman; diamonds, supported by the flow cohort (h). **b**, *ρ* per neutrophil state and tissue (circles, all genes, filled if gene-label permutation *P <* 0.05; diamonds, genes with *P <* 0.05 in both); right, exact time-relabelling *P*. **c**, *ρ* of every cell type, ranked per tissue; *P*, exact rank-sum test of neutrophil ranks. **d**, Blood-timer score of pooled neutrophils per sample with cosinor fit, peak and permutation *P*. **e**, *ρ* on all (circles), non-enriched (squares) and neutrophil-enriched (triangles) genes. **f**, *ρ* of control and neutrophil-depleted long-bone marrow rhythms (GSE198770; *n* = 3 per time) with the blood programme. **g**, Human blood neutrophils (GSE136632; 14:00 versus 08:00 as ZT19 versus ZT13): Pearson *r* with our fits (filled, one-sided permutation *P <* 0.05); squares, bulk cochlea (GSE88954; marker genes; gene-level *P*). **h**, Pre-specified flow cytometry: contour plots of Ly6G^+^ SSC^hi^ cells of the median mouse and % CXCR4-hi and CD62L-hi of Ly6G^+^ cells per mouse (bar, geometric mean; whiskers, 95% CI); *P*, ZT17 versus ZT5 within tissue, linear mixed model (n.s., *P ≥* 0.05); *n* = 10 marrows and 10 cochleae per time, except 9 cochleae at ZT5 for CXCR4. *n* = 6 samples per tissue (a–e).

In long-bone marrow ^35^, the inverse signature fell from *ρ* = *−*0.18 to *−*0.03 when neutrophils were depleted (Fig. 6f). The positive, cochlear relation recurred in bulk cochlea (neutrophil-marker genes, *ρ* = 0.43)^14^ and in human blood neutrophils sampled at 14:00 and 08:00 (*r* = 0.20 with our cochlear fits, *P <* 0.001; *−*0.09 with marrow fits) ^44^, and the blood programme itself recurred in independent single-cell data (higher at ZT13 than at ZT5; cell-level *P* = 0.010) ^23^ (Fig. 6g).

At the protein level, the flow cohort showed the predicted split for CXCR4, which marks aged circulating neutrophils^21,22^: CXCR4-high neutrophils were less frequent at night in the cochlea (odds ratio ZT17/ZT5 0.29, 95% CI 0.18–0.47; *P <* 0.001) but not in the marrow (0.91; *P* = 0.70; tissue *×* time interaction *P <* 0.001; Fig. 6h). CD62L, predicted to rise at night in the cochlea, did not change significantly (odds ratio 1.20, *P* = 0.35; interaction *P* = 0.21). With six samples per tissue, no single correlation reached an exact time-relabelling *P* below 0.067 (Fig. 6b). Tested as one statistic against independent relabelling of both tissues’ samples, the split was significant for aged neutrophils (*P* = 0.022) but not for neutrophils pooled (*P* = 0.099; Extended Data Fig. 9h). The strongest evidence for this tissue contrast therefore came from the aged-neutrophil transcript programme and the CXCR4 tissue-by-time interaction.

### An independent cohort supports daily rhythms and shows noise-associated heat-shock changes

An independent cochlear data set with noise exposure by day or by night^28^ lacked collection times, so we estimated each library’s collection phase from its clock-gene profile: single-cell controls fell at ZT1.7, day-noise libraries at ZT1.0 and night-noise libraries at ZT13.9, 11.2 and 11.8 h from the day-noise and control libraries and beyond the upper 95% bound of a clock shift attributable to noise (5.5 h; Fig. 7a,b and Extended Data Fig. 10a–c). We mapped 110,912 of its cells onto our labels, 72% of them confidently (Fig. 7c).

**Fig. 7.**
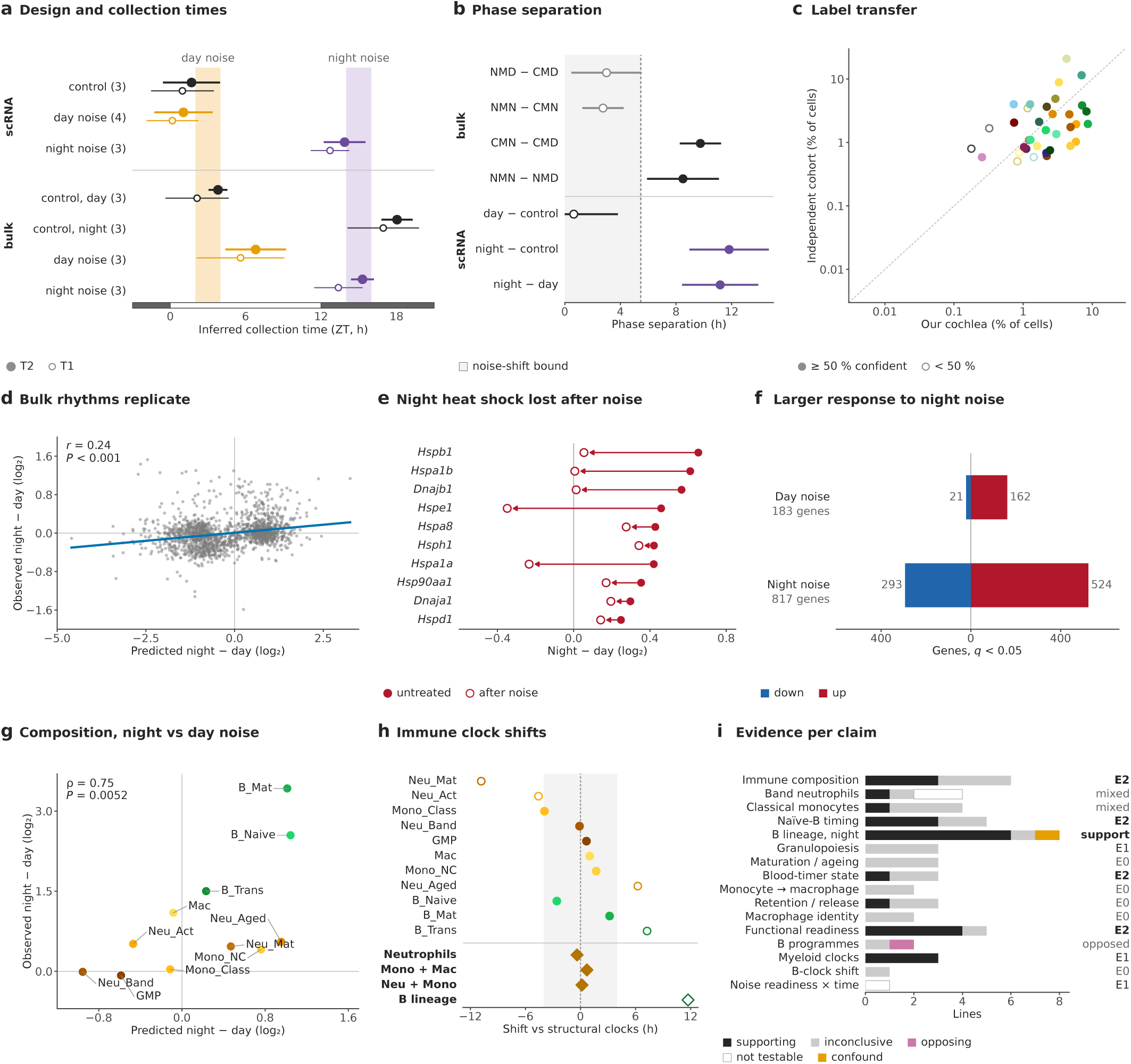
Independent support for daily rhythms and exploratory noise-associated responses. Independent cochlear data ^28^ (collection times not reported): single-cell libraries of untreated mice and 3 days after day or night noise; bulk RNA-seq of untreated cochleae (day, CMD; night, CMN) and 1 day after day (NMD) or night (NMN) noise (*n* = 3 per group). **a**, Collection phase of each group inferred from core-clock genes (filled, primary estimator with 95% interval; open, alternative); shading, noise exposures; numbers, libraries; negative ZT, the preceding night. **b**, Phase separation between groups (95% interval); grey, up to the upper 95% bound of a noise-induced clock shift; filled, beyond it. **c**, Share of each transferred label among 110,912 cohort cells versus our cochlea. **d**, Untreated bulk: night*−*day difference per gene (1,595 non-clock genes rhythmic in our cochlea) predicted from our fits at the inferred phases versus observed; *r*, Pearson; *P*, gene-label permutation. **e**, Night*−*day difference of ten heat-shock genes, untreated and after noise. **f**, Genes changed 1 day after day or night noise (*t*-test of contrasts; Benjamini–Hochberg *q <* 0.05). **g**–**i**, Exploratory analyses: **g**, change in each label’s share between night-and day-noise libraries, observed versus predicted from our abundance rhythms (*ρ*, Spearman; *P*, one-sided permutation); **h**, night*−*day clock shift of each immune cell type relative to structural cells (grey, *±*4 h; filled, within 4 h; diamonds, pooled populations); **i**, evidence lines per claim and the exploratory grade (E2, support; E1, qualified support; E0, none).

In time-matched untreated bulk cochlea, the observed night–day difference followed the difference predicted from our cochlear rhythms (*r* = 0.24 over 1,595 genes; *P <* 0.001), and all ten heat-shock genes were higher at night (mean +0.45 log_2_; Fig. 7d,e). After noise, this night rise was absent (+0.06), and night noise elicited the larger response: 817 genes changed one day after night noise, against 183 after day noise (*q <* 0.05; Fig. 7f), although exposure and collection times coincide in this design. In an exploratory analysis, the immune composition of cochleae exposed to noise at night or during the day, collected about 12 h apart, differed as our abundance rhythms predict (*ρ* = 0.75), if noise *×* time interactions are small (Fig. 7g–i and Extended Data Fig. 10e).

## Discussion

Three-dimensional reconstruction maps marrow-associated vessels connecting to the cochlea, and complementary imaging locates neutrophils beside these vessels and within cochlear fluids. Parabiosis findings are consistent with a local contribution to this immune compartment. Across the day, structural cochlear cells retain light-entrained clocks, while cochlear neutrophils follow a programme aligned with circulating neutrophils and opposed by those in marrow. The anatomical and temporal observations thus allow cell location, origin evidence and daily programme to be considered separately within the same sensory system.

The reconstruction preserves the spatial relationship between marrow cavities, connecting vessels and the cochlea. Tissue dissociation removes this context, leaving cell profiles without their original relationship to bone and vasculature. The anatomical map also identifies a route for future source-specific labelling and live-cell tracking. It builds on vascular connections mapped between temporal marrow and the rat labyrinth ^12^ and on marrow cells observed in bony channels to the endolymphatic sac during experimental labyrinthitis^45^. In healthy mice, we combine this anatomy with neutrophil localization, parabiosis and parallel temporal profiles to interpret source and timing observations across physically connected compartments. The images establish a vascular connection, but cell passage through it, including the direction and rate of movement, was not measured.

The anatomical and cellular observations support a local marrow contribution to cochlear immune cell supply as a working hypothesis. The evidence concerns the myeloid populations examined here, including neutrophils, and does not assign a local origin to all cochlear immune lineages. Neutrophils were present in healthy cochlear fluids, and partner-derived cells were less dense in the cochlea than in adjacent marrow at 14 and 54 days of parabiosis. Host-derived cells could nevertheless arrive through the host circulation or persist locally; host origin alone does not identify a marrow site. Region-specific fate mapping is needed to distinguish these routes. The hypothesis that cochlea-associated marrow forms part of a local immune niche extends work at the borders of the brain and spinal cord ^6–8^ to a peripheral sensory organ.

Signals from the inner ear might in turn influence adjacent marrow. Cerebrospinal fluid can access skull marrow and influence its niches^9,10^, but comparable communication from the inner ear has not been established. Such local signals could couple immune supply to the condition of the sensory organ, while systemic cues could coordinate both tissues. Neither process was demonstrated here. The anatomical map and cell-state reference identify where these possibilities could be tested, including before and after cochlear injury.

Cochlear neutrophils followed the daily programme of blood neutrophils rather than that of adjacent marrow, despite their anatomical proximity and compatible origin profiles. One interpretation is that the neutrophil timer ^21,22^ operates after marrow release, whether cells enter the systemic circulation or a neighbouring tissue. Alternatively, the cochlea may recruit or retain cells at particular stages of that programme, consistent with tissue-dependent neutrophil states^23^. Local signals could also help produce similar states. The strongest transcriptomic support came from the most mature neutrophil state, with independent protein-level support from CXCR4. The data support testing a post-release timer but do not establish when or where the programme is set. An *imported* immune timetable remains a model for this contrast with local structural-cell clocks.

Daily variation in immune composition and heat-shock responses may be relevant to cochlear physiology. Recruitment rhythms can arise through adrenergic signals and endothelial or leukocyte clocks^16,18^, so composition alone cannot identify the source of timing. The heat-shock programme rose at night in cochlea and marrow and persisted after adrenalectomy in independent cochlear data. Heat-shock responses can protect the cochlea: prior heat stress reduces permanent noise injury ^46^, HSF1 contributes to hair-cell survival after acoustic overexposure^47^, and pharmacological HSF1 activation protects spiral ganglion neurons ^32^. The normal night rise of heat-shock genes was absent after exposure in the independent noise cohort ^28^. This association suggests a relationship between temporal state and stress responses, alongside known glucocorticoid effects ^14^. Direct tests are needed to determine whether immune timing or heat-shock capacity mediates time-dependent hearing vulnerability.

Each tissue and time point is represented by one pooled sample. Exact within-tissue time-relabelling tests therefore have a minimum *P* of 1/60, and the tissue-wide sample pattern remains a potential confounder. Calibration on held-out data, specificity checks and independent replication address aspects of this problem but do not replace biological replication across full time series. The joint neutrophil contrast was significant for aged but not pooled neutrophils, and immune cells contained too few clock transcripts for their single-cell clocks to be compared reliably with structural-cell clocks. Sampling under a light–dark cycle establishes day–night variation; endogenous circadian control was not demonstrated for every programme.

Cochlear preparations differed in marrow content, for which unadjusted and adjusted analyses are reported. Flow cytometry tested two times of day, and digestion reduced CD62L and CXCR4 signals, requiring within-tissue comparisons. Parabiosis included one pair per pairing duration, blood chimerism was taken from published pairs, and the brain comparison used one rendered projection. Partner-derived cells were counted per tissue area rather than per immune cell, and GFP settings differed between compartments. These constraints limit source attribution despite the detection checks and blinded review by research personnel. Noise-cohort collection times were inferred, leaving single-cell exposure contrasts confounded with collection time. *Arntl* and sex-chromosome genes were absent from the atlas, and all mice were male.

Labelling marrow specifically within the temporal bone, together with contemporaneous blood measurements and directional tracking through the reconstructed connection, would test its contribution to cochlear immune cells. Following newly released neutrophils through local and systemic routes would distinguish acquisition of a post-release programme from selective recruitment of an existing state. Time-matched injury experiments could test whether local marrow responses and cochlear immune composition are coordinated, and whether this coordination affects hearing outcomes. Constant-darkness sampling would establish which observed programmes are endogenous circadian rhythms.

The annotated atlas, three-dimensional anatomical views, rhythm catalogue, hearing-related gene rhythms (Supplementary Table 6), calibrated analysis workflow and Source Data are provided for these investigations. They also allow cochlear experiments to be interpreted with attention to cell type, neighbouring marrow and collection time, and can be used to examine how structural clocks and immune cell supply contribute to inner-ear homeostasis.

## Methods

### Animals

All procedures were approved by the Institutional Animal Care and Use Committee of the Affiliated Hospital of Guangdong Medical University (approval no. AHGDMU-LAC-B-202508-0078) and followed the Guide for the Care and Use of Laboratory Animals and the ARRIVE guidelines. Atlas mice were specific-pathogen-free C57BL/6J mice aged 6–8 weeks from the Animal Center of the First Affiliated Hospital of Guangdong Medical University, housed four per cage in stable groups with standard chow and water ad libitum at 22 *±* 2 *^◦^*C and 55 *±* 10% relative humidity under a 12-h light–dark cycle (lights on at 08:00, ZT0) for at least 14 days before sampling. The flow-cytometry cohort comprised 20 C57BL/6J mice aged 10 weeks from the same supplier, housed and entrained in the same way. Imaging and spatial transcriptomics used C57BL/6J mice aged 6–8 weeks. Parabiosis joined age-and weight-matched 10-week-old wild-type C57BL/6J and Lyz2-P2A-iCre-P2A-EGFP (Lyz2-EGFP) mice, in which EGFP is co-expressed with the myeloid gene *Lyz2*; seven pairs were joined, one per pairing duration (5, 7, 14, 28 and 54 days, and two at 31 days, one of them exposed to noise). All mice were male (the flow cohort matched the atlas by design); sex was not a study variable, and sex-based analyses were not performed.

### Tissue collection and single-cell RNA sequencing

Mice were sampled at ZT3, 7, 11, 15, 19 and 23 in a single series that began at ZT3 on 20 August 2025, with dim red light for the dark-phase collections. They were anaesthetized with isoflurane (4% for induction and 2% for maintenance) and perfused transcardially with 20 ml of ice-cold PBS at 5 ml min*^−^*^1^. The cochlear soft tissue, separated from the vestibule and stripped of bone, formed the cochlear sample, and the apical bone fragments containing marrow formed the marrow sample (Supplementary Note 1). Cells of three mice were pooled per tissue and time point into one independently processed sample and library. Single-cell suspensions were prepared with the 10x Genomics tissue dissociation procedure (viability above 80%; Supplementary Note 1). Cochlear libraries were prepared with 10x Genomics Chromium Single Cell 3*^′^* v4 chemistry on a Chromium X and marrow libraries with the MobiSeq-V2 workflow (MobiCube Single-Cell 3*^′^* RNA-seq Kit v2.0) on a MobiNova-100 Single-Cell System (MobiDrop), and all were sequenced on an Illumina NovaSeq 6000 (10x libraries, 28 + 90 bp; MobiSeq libraries, 2 *×* 150 bp; about 300 million read pairs per 5,000 recovered cells). FASTQ files were processed into filtered count matrices with Cell Ranger 10.0.0 and MobiVision 2.0 against the GRCm39 genome, with gene identifiers mapped to MGI symbols.

### Quality control and annotation

The 12 libraries of this study were processed together with 30 libraries from other cochlear experiments (42 in total), and cell types were annotated on the combined data (Supplementary Note 1). The matrices were restricted to protein-coding and mitochondrially encoded genes of the GRCm39 autosomes and mitochondrial chromosome, which excludes sex-chromosome genes. Cells were filtered on detected genes, UMIs and mitochondrial, haemoglobin and ribosomal transcript fractions (OmicVerse^48^), doublets were removed per library with Scrublet ^49^, and raw counts were kept for all downstream analyses. Cells were clustered by landmark-based clustering (an in-house package) on the nearest-neighbour graph of a library-conditioned scVI ^50^ latent space of highly variable genes (Seurat v3 method^51^) and annotated by matching cluster marker genes to curated immune and cochlear structural gene sets, with markers confirmed by pseudobulk differential expression (PyDESeq2 ^52^). The resulting atlas comprised 160,643 cells in 35 subtypes of 12 classes (Supplementary Table 1). These labels were used throughout without re-annotation. Labels and the mixing of samples were audited per sample (Extended Data Fig. 1). Technical metrics (depth, detected genes, mitochondrial fraction and dissociation stress) were tested for rhythmicity and reported but were not included as covariates. No ambient-RNA or blood covariates were modelled; tissue perfusion and quality control were performed as described above.

### Rhythm-aware alignment of single-cell data

Because each library represented one tissue at one time point, correcting for library or sample, as in the integration used for annotation, also removes differences between times of day; the labels of that integration were kept after the per-sample checks above, but its latent space was not used for any time-dependent analysis. Embeddings were instead chosen per tissue and jointly by a benchmark with pre-specified selection rules (Supplementary Note 2), which scored conservation of cell identity (scib-metrics ^53^, including cell-type LISI^54^), time-of-day structure relative to a principal component analysis (PCA) of the same genes and, in the joint data, mixing of the two platforms (including batch-removal-adapted silhouette^55^). On feature sets screened by PCA, the candidates were PCA, Harmony^54^, Scanorama ^56^, BBKNN ^57^, sysVI ^58^, scVI ^50^ and scANVI ^59^, with no batch, sample or, in the joint data, platform as batch. Candidates were ranked by a weighted composite of the scores, and embeddings retaining less than 80% of the baseline time-of-day structure were rejected: all models with sample as batch or covariate (36–74% kept) and BBKNN (63%). The selected embeddings were scANVI without a batch covariate for the cochlea (5,000 genes, 30 dimensions; 93% kept) and the marrow (3,000 genes, 50 dimensions) and scANVI conditioned on platform only for the joint data (3,000 genes, 50 dimensions). They defined the neighbourhood graphs for the abundance tests and display; diffusion pseudotime and trajectories used unsupervised scVI without a batch covariate, on which embedding-dependent results were also repeated. Expression rhythms were tested on raw counts summed per sample, without any embedding (embeddings were analysed with Scanpy ^60^). External cells were mapped onto a scANVI reference conditioned on sample (scArches surgery^61^) only to transfer identity labels, never to estimate time.

### Rhythm inference with one pooled sample per time point

Counts were summed per tissue, cell type and sample. A cell type entered rhythm analyses with at least 20 cells in each of the six samples (types with 10–19 cells in a sample were analysed but flagged). Per cell type, a cosinor model ^62^ was fitted with limma–voom ^63,64^ (TMM normalization ^65^, robust empirical Bayes ^66^, moderated *F*-test of the cosine and sine terms; Supplementary Note 3), giving amplitude, peak phase, peak-to-trough fold change and *P* values. Because six equally spaced samples admit 60 distinct assignments to times up to rotation and reflection, each gene was refitted under the 59 non-identity relabelling classes, giving exact per-gene *P* values and a pooled-null empirical false-discovery rate. Six leave-one-time-point-out (LOTO) refits gave the circular s.d. of the phase; a rhythm was single-sample-driven if it collapsed when one time point was left out. Independent tests were the edgeR quasi-likelihood *F*-test ^67,68^, RAIN ^69^ and JTK_CYCLE^70^ in MetaCycle^71^. Tier 1 required a Benjamini–Hochberg ^72^ *q <* 0.05, a permutation *q <*0.10, a peak-to-trough fold change *≥* 1.5, an independent test with *P <* 0.05, a LOTO phase s.d. *<* 3 h and no single-sample dependence; Tier 2 required *q <* 0.10 and a fold change *≥* 1.5. The share of rhythmic genes was estimated as *π*_1_ (ref. 73). Cell-type summaries and all rhythm calls are in Supplementary Tables 2 and 3. The pipeline was calibrated on dense public time series subsampled to our design (GSE54650^33^, 12 organs in constant darkness; GSE88954^14^, sham-operated cochlea) and with rhythms injected into our own relabelled data. Analytic power used the sampling variance of the cosinor coefficients including the voom precision weights (Extended Data Fig. 4).

### Clocks

Core-clock genes were fitted per cell type. A clock phase offset was the amplitude-weighted circular mean of observed minus canonical peaks ^34^ (*Npas2*, the E-box partner with the phase of *Arntl*, stood in for the undetected *Arntl*), checked by phase coherence, the circular correlation of observed and canonical peaks and the clock correlation distance to the mouse pan-tissue reference^74^. A joint alignment estimated gene reference phases and offsets across our tissues and external data by alternating weighted circular means (Supplementary Note 4). Single-cell clock phase was the projection of *z*-scored log-normalized clock-gene expression onto the canonical peak phases, and the consistency of each cell type’s per-sample offset (mean phase minus ZT) was tested against the 60 relabelling classes; because haematopoietic cells carry fewer clock transcripts, the comparison was repeated at matched clock-transcript depth, with a pre-specified rule for judging the structural–haematopoietic contrast power-limited. External clock profiles estimated sampling phase by template matching, validated by leaving one sample out of each reference. Clock-gene fits are in Supplementary Table 4.

### Systemic programmes, motifs and rhythm drivers

Pathway and transcription-factor activities (CollecTRI regulons^75^, univariate linear model in decoupler ^76^) were scored per cell type and sample, tested as above in primary, depth-matched and sample-centred versions and compared with the factor’s own mRNA phase (Rayleigh test). Five systemic-cue programmes (glucocorticoid response, adrenergic–cAMP, heat shock (HSF1), feeding (insulin–mTOR) and hypoxia (HIF)) were tested per cell type and summarized by their amplitude-weighted common phase and its resultant length. Clock and cue-responsive promoter motifs were tested for enrichment among rhythmic genes and for the phase of rhythmic carriers. Each rhythm call was assigned one primary driver class from evidence flags. Further tests compared calls with the phases of the same genes in constant darkness (GSE54650) and with dissociation-stress profiles (handling), and used independent cochlear series with and without adrenalectomy (GSE88954; glucocorticoid dependence) and without noise (GSE107086^14^; night–day heat-shock check) (Supplementary Note 5).

### Composition and neighbourhoods

Cell-type shares were tested with a 24-h cosinor in several frames and models, with least squares on percentages of tissue cells as primary (as plotted); for cochlear immune types, a Bayesian model (NumPyro ^77^) separated each share into a component scaling with the sample’s marrow content and a residual 24-h rhythm (sensitivity analysis; Supplementary Note 6). Five core models (centred log-ratio with permutation, propeller ^78^, sccomp ^79^, quasi-binomial and beta-binomial generalized linear models) defined the multi-model verdict: robust, at least four supporting with LOTO and across-model phase s.d. below 3 h; supported, two or three (Supplementary Table 5). Neighbourhood abundance was tested with a sparse re-implementation of Milo ^80^ on the scANVI neighbourhood graph (edgeR quasi-likelihood cosinor with a spatial false-discovery rate) and repeated on unsupervised scVI embeddings. Production waves related each neighbourhood’s peak time to the diffusion pseudotime^81^ of its index cell by circular–linear correlation, with pseudotime-permutation *P* values. B-cell stages were matched to ImmGen populations (microarray, GSE15907^82^; RNA-seq, GSE109125^83^) by Spearman correlation of pseudobulk profiles.

### Comparison with circulating neutrophil programmes

The reference day–night programme was derived from wild-type blood neutrophils at ZT5 and ZT13 (GSE86619^22^; *n* = 3 each). For every neutrophil state and tissue, Spearman’s *ρ* between the change from ZT5 to ZT13 predicted by our cosinor fit and the observed change was computed across shared genes, with gene-label permutation *P* values (Supplementary Note 7). Specificity was tested by ranking the neutrophil states among all primary cell types of a tissue (exact rank-sum test) and against the exact relabelling null of each correlation. Because cochlea and marrow come from different mice, the split was also tested as one statistic, the cochlear minus the marrow correlation, against all combinations of the two tissues’ relabelling classes (one-sided; aged and pooled neutrophils pre-specified) and against all permutations of their time labels. The blood-timer score of a cell contrasted the genes most increased and most decreased between ZT5 and ZT13 in GSE86619, and per-sample medians of pooled neutrophils were fitted by cosinor with permutation *P*. Neutrophil dependence used long-bone marrow with and without neutrophil depletion (GSE198770^35^; *n* = 3 per time). Human relevance used blood neutrophils of three donors sampled at 08:00 and 14:00 (GSE136632^44^), mapped to activity-matched mouse times (ZT13 and ZT19) through one-to-one orthologues^84^, and blood single-cell data at ZT5 and ZT13 (GSE142754^23^; cell-level Mann–Whitney test). The maturation score contrasted the genes most positively and most negatively correlated with diffusion pseudotime in reference neutrophils (GSE137540^42^).

### External data, spatial deconvolution and attribution

External cochlear atlases (GSE283534^29^, GSE135913^30^, GSE274279^27^ and GSE234926^31^) were processed from raw matrices with emptyDrops ^85^, per-sample quality control and Scrublet ^49^, without ambient-RNA correction, and our labels were transferred with scANVI and a *k*-nearest-neighbour vote (Supplementary Note 8). Spatial transcriptomes (GeoMx; GSE312646^32^; control areas) were deconvolved by weighted non-negative least squares on our cochlear profiles, and regions were compared by two-sided Welch’s *t*-tests with Benjamini–Hochberg correction. Replicated bulk rhythms (GSE88954 sham cochlea; GSE198770 control marrow) were attributed by correlating each gene’s external time course, after clock alignment, with composition-only and regulation-only profiles built from our pseudobulk, against permutation and bootstrap nulls. Sorted marrow neutrophils (GSE312298^43^; PBS arms, ZT14 and ZT22) were compared with predictions from 644 selection-free Tier-1 genes of our neutrophil states. The baboon atlas (GSE98965^40^) and constant-darkness organs were compared after clock alignment, through one-to-one orthologues for baboon.

### Independent noise cohort

In the cohort of ref. 28 (CRA019820 and CRA019827; design, Supplementary Note 8), male CBA/CaJ mice (lights on at 06:00) were exposed to noise by day (ZT2–4) or night (ZT14–16); single-cell libraries were made before and 3 days after noise and bulk RNA-seq before and 1 day after. Single-cell reads were quantified with STARsolo 2.7.11b ^86^ and processed as above. Because collection times were not reported, library phases were estimated by clock-gene template matching (primary template, the independent sham-cochlea series), and clock genes were therefore excluded from the gene sets that validated rhythms in this cohort. The bound for a noise-induced clock shift (5.5 h) was the upper 95% limit of the 1-day bulk noise shifts; bulk differences were tested by *t*-tests of contrasts in a four-group linear model on log_2_ CPM with Benjamini–Hochberg correction. All analyses of this cohort were exploratory, with rules and timing assumptions fixed before the first run.

### Three-dimensional imaging of the cochlea and its marrow

Whole-cochlea imaging (Fig. 4a,b) was adapted from whole-mount cochlear clearing ^87^. Vessels were labelled in situ by tail-vein injection of DyLight 649-labelled tomato (*Lycopersicon esculentum*) lectin (Vector Laboratories, DL-1178-1), adapted from cranial marrow imaging ^11^, before perfusion-fixation; cochleae were decalcified and immunostained for neutrophils (Supplementary Note 9). Cochleae cleared with a modified ScaleS protocol were imaged on a ZEISS LSM 900 confocal microscope as tiled z-stacks for three-dimensional reconstruction. The tiled z-stacks provided the spatial basis for the anatomical reconstruction. Volumes were processed in Fiji^88^ and rendered in Imaris 11.0 without non-linear intensity remapping, and vessels linking marrow and cochlea were traced by hand on the renders (Fig. 4a). The tracing was checked against vessel ridges detected in the unannotated renders of the same views without reference to it, and against 999 random rotations and translations of the same traces (Supplementary Fig. 1).

### Spatial transcriptomics

For Stereo-seq (Fig. 4c), cochleae of an untreated male mouse killed by cervical dislocation were dissected under RNase-free conditions, formalin-fixed, decalcified and paraffin-embedded (Supplementary Note 9). Sections were profiled on the Stereo-seq platform ^89^, and FASTQ files were processed with the Stereo-seq Analysis Workflow (SAW 1.0) ^90^ into expression matrices at bin-50 resolution (53,877 spots). Cell types of our atlas were mapped onto the spots with Tangram ^91^ and, as a complementary estimate, with Cell2location ^92^; Fig. 4c shows the Cell2location estimate.

### Cochlear fluids and confocal image analysis

For Fig. 4d, perilymph, endolymph and marrow were sampled microsurgically from intact cochleae (Supplementary Note 9); endolymph came from dedicated mice, perilymph and marrow sequentially from the same mice. Cells were cytocentrifuged onto poly-L-lysine-coated slides, fixed, immunostained for FUT4 (rabbit polyclonal; Bioss, bs-1702R) and CD45 (mouse monoclonal, clone 4E9B2; Proteintech, 60287-1-Ig) and mounted with DAPI, alongside unstained, secondary-only, single-stained and biological positive and negative controls imaged with identical settings (Supplementary Note 9). Confocal images (Olympus FV3000, 60*×* objective) were analysed from the 16-bit acquisitions. Panels show crops with one linear display range per channel across compared images, without gamma, filtering or background subtraction. In marrow, perilymph and endolymph preparations of wild-type mice, nuclei were segmented from DAPI, and a cell was FUT4^+^ (or CD45^+^) when its signal (mean over the cell) was at least twice its local background. Where CD45 was excited only by other laser lines (561-nm line at 0%), its signal was not spill-over from FUT4 (2.6-fold larger relative to FUT4 than in marrow, rim-enriched), so CD45 there marks presence only and its intensity was not compared between preparations. Image analysis used scikit-image^93^.

### Parabiosis

Parabiosis followed a standard protocol^94^: the wild-type and Lyz2-EGFP partners were anaesthetized with isoflurane (4% for induction and 2% for maintenance, in oxygen), joined along their adjacent flanks, given analgesia (buprenorphine and carprofen) and antibiotics (enrofloxacin; doses in Supplementary Note 9) and co-housed for 5, 7, 14, 28, 31 or 54 days (blood chimerism stabilizes within 2–3 weeks). A second 31-day pair was exposed to 2–20 kHz noise at 107 dB SPL for 2 h; tissues were collected 3 days after exposure. Noise exposure recruits monocytes into the cochlea ^1^. Under deep terminal anaesthesia, mice were perfused transcardially with 20 ml of ice-cold PBS at 5 ml min*^−^*^1^, and dissected tissues were fixed in 4% paraformaldehyde overnight at 4 *^◦^*C, labelled with lectin or immunostained (FUT4, CD45, CD19 or CD14) and imaged by confocal microscopy (ZEISS LSM 900). In a wild-type partner, GFP marks partner-derived *Lyz2*-expressing myeloid cells that entered through the shared circulation, and GFP*^−^* myeloid cells, such as the FUT4^+^ neutrophils analysed, are host-derived. Every native acquisition of the wild-type partners with a GFP channel and pixels of at most 0.70 *µ*m was analysed by fixed rules (Supplementary Note 9), set before analysis and extended once to include every usable acquisition. GFP^+^ bodies (GFP above the tissue median plus 10 robust s.d.) counted as partner-derived cells when cell-sized (at least 25 *µ*m^2^), at least twice as bright as the tissue 2–6 *µ*m around them, not equally bright in other channels and, where DAPI was imaged, nucleated; this upgraded cell rule took its conventions from the other rules of this study and was fixed before any count with it was inspected. Cells were counted per mm^2^ of tissue in optical sections at least 10 *µ*m apart. Perilymph cells were GFP^+^ when the GFP of their cytoplasmic ring was at least twice the local background. Immune cells of the 54-day partner were typed on segmented nuclei: a cell was its nucleus with a 1.5-*µ*m ring; it was marker-positive when at least half of the ring exceeded the series threshold, the ring was at least twice its local surroundings and the positive ring surrounded the nucleus, and GFP^+^ when at least half of it lay in a GFP^+^ body at least twice as bright as its surroundings (Supplementary Fig. 6). Acquisitions of one position (stage coordinates) were counted once, and unlabelled fields were assigned to tissues by morphology (shown in Supplementary Fig. 2a). GFP settings were set per sample, so cochlear counts were checked against the detectability of the same animal’s marrow GFP^+^ bodies (laser transmission and detector gain, calibrated on one position imaged at two voltages) and against depth (each acquisition’s brightest section). Pairs were the unit of comparison (exact sign test); the noise-exposed pair was placed among the unexposed pairs (z-score and 95% prediction interval on the log scale), with sections resampled within animals (bootstrap) for its density ratios. The 54-day animal was also imaged in 60*×* sections (Olympus FV3000; Fig. 4f,i); there, every marrow GFP^+^ cell imaged with the cochlear settings cleared the cochlear threshold. The brain was available only as one rendered maximum projection of the 31-day partner (GFP and lectin, without DAPI or pixel size); it was counted at 2.773 *µ*m per pixel, the collection’s usual overview setting (its other overview pixel sizes as a sensitivity range). To compare tissues, every field was converted to that format (one section at a time, box-averaged to 2.773 *µ*m, GFP and lectin only), GFP^+^ bodies were detected on the local background (white top-hat of 15 *µ*m, at 10 robust s.d.; fixed before its run and used because it agreed better with the blinded labels than the global threshold), and all tissues were counted by a stricter rule chosen from a fixed grid of signal, contrast, area and nucleus thresholds by agreement (F1) with the blinded labels; in that format only contrast and area were tightened, because the saturated brain rendering caps body signal (Supplementary Note 9). Blood chimerism was taken from published parabiosis (22.1 *±* 2.0% of blood neutrophils after one month with Lyz2-GFP or CD45.1 partners ^41^; about 30% of blood neutrophils and monocytes after 60 days with UBC-GFP partners^7^).

### Verification and robustness of the image analysis

Research personnel checked the image calls by blinded visual review. Seeded random samples of the calls, drawn within each pair and compartment (survey bodies kept or rejected as cells) or each marker and compartment (typed and untyped cells, and every partner-derived call), were shown as crops identified only by a code, without the call, compartment or file, and were labelled by criteria fixed before labelling; agreement is reported with Wilson 95% CIs (Supplementary Fig. 7). The two upgraded rules were developed on a first labelled sample and scored on a fresh one. For the survey, partner-derived cells per pair and compartment were corrected with these labels (kept bodies times the share confirmed, plus rejected bodies times the share that were cells; 10,000 bootstrap draws within strata), and fewer partner-derived cells in the cochlea than in the marrow was claimed only for pairs whose upper 95% limit of the corrected ratio lay below 1. Every image rule and the flow gates were re-run over grids of their parameters (thresholds, sizes, contrasts and ring definitions; flow gate thresholds moved by *±*10%), and each claim was re-evaluated at every setting (Supplementary Fig. 7 and Supplementary Note 9).

### Flow cytometry

The validation cohort, its readouts, models and exclusions followed a plan written before any flow data were seen (Supplementary Note 10). The 20 mice were sampled at ZT5 (13:00) or ZT17 (01:00) in two cohorts of five mice per time point, with cage-balanced allocation, coded sample identifiers, the order of the sampling windows reversed in the second cohort and dim red light for night handling. After transcardial perfusion with ice-cold PBS under terminal anaesthesia, both cochleae of each mouse formed one preparation, which retained the otic capsule and modiolus with a fixed boundary to the adjacent marrow (a published preparation removed the bony capsule without exposing the marrow^95^), and the bilateral temporal-bone marrow a second (40 paired mouse–tissue samples; the two time points comprise different mice). Cochleae were digested enzymatically and marrow was released mechanically; cells were Fc-blocked (BD Fc Block, clone 2.4G2) and stained with BioLegend antibodies (1:100) in eight two-colour (FITC/PE) panels, T1–T8 (Supplementary Note 11). Ly6G was detected with clone 1A8 rather than the Gr-1 antibody RB6-8C5, which also binds Ly6C ^95^. Tubes for which insufficient cells were recovered (43 of 320, mostly cochlear T6 and T7) were treated as missing. Data were acquired on a BD FACSCelesta with BD FACSDiva 9.0, expressed in bead-relative fluorescence units and gated in FlowJo 11.2 and custom Python code with fixed gates set on the controls and locked before group comparisons (Supplementary Fig. 8 and Supplementary Note 12).

Primary readouts, each a percentage of CD45^+^ cells, were cochlear immature neutrophils (P1; Ly6G^int^ CD11b^int^), cochlear classical monocytes (P2; CD11b^+^ Ly6C^hi^ cells in the monocyte scatter gate) and marrow recirculating B cells (P3; B220^hi^ IgD^+^); these marker-defined proxies do not test same-event Ly6G exclusion, F4/80, CX3CR1 or viability, which published cochlear monocyte gating used ^95^. With the mouse as the unit of analysis, primary readouts were tested by linear models on log percentage with time and cohort and Holm correction ^96^ across the three (replicated, Holm *P <* 0.05 in the predicted direction), and secondary readouts (CD62L^hi^, S1, and CXCR4^hi^, S2, percentages of Ly6G^+^ cells; marker medians) by linear mixed models with tissue *×* time, cohort and a random intercept per mouse. The exclusions (cochlear tubes with fewer than 3,000 single-cell events, parent gates with fewer than 100 events, viability below 50%) removed one ZT5 cochlea from P1, P2 and S2; no parent gate fell below 100 events, and no sample fell below 50% viability (trypan blue exclusion, 78–96%, used because every cochlear T8 tube fell below the event floor). Sensitivity analyses repeated P1 and P2 without samples failing the consistency check between tubes and divided them by a marrow-content index. Perfusion reduces but does not remove intravascular leukocytes, so the readouts describe the recovered preparation. Per-mouse readouts and all tests are in Supplementary Tables 7 and 8.

### Evidence grading

Positive findings were graded on pre-specified evidence dimensions into verdicts (robust, supported, suggestive, not supported) by rules fixed before the outputs were inspected (Supplementary Note 13); the wording of each claim follows its verdict, and Supplementary Table 9 lists every graded finding. Pre-specified B-lineage and myeloid hypotheses were decided by their own rules (Extended Data Figs. 8c and 9a; Supplementary Tables 10–19).

## Statistics and reproducibility

For the atlas, the unit of replication is one pooled sample per tissue and time point (*n* = 6 per tissue); cells were never treated as replicates in rhythm tests, and the single cell-level test (circulating-neutrophil programme score) is labelled as such. Exact time-relabelling *P* values have a minimum of 1/60; unless stated otherwise, tests are two-sided, and Benjamini–Hochberg correction was applied within stated families. No statistical method was used to predetermine the size of the atlas; the flow cohort size was set by the pre-specified power analysis (Supplementary Note 10). Linear models were fitted on log or logit scales without formal tests of normality. Atlas mice were randomly assigned to sampling times at the individual level; flow samples carried coded identifiers, and the flow analyst was blind to ZT; image quantification used fixed automated rules applied identically to all fields and was checked by blinded visual review (the tissue of unlabelled parabiosis fields was assigned without blinding). The atlas was not repeated as a whole; its claims were tested in independent data (external atlases, spatial data, dense time series, primate, human, sorted neutrophils, noise cohort), by imaging and parabiosis, and in the pre-specified flow cohort. The structures shown in Fig. 4a,b,d were observed consistently in at least 10 mice; Stereo-seq (Fig. 4c) is one mouse, and parabiosis has one pair per pairing duration (Fig. 4e–i). Exclusions were limited to the pre-specified flow rules above and the parabiosis acquisition rules (Supplementary Note 9).

## Software

Trajectory checks of the pre-specified myeloid hypotheses used CellRank 2.3.3 (refs. 97,98) and Palantir 1.4.5 (ref. 99). All software and package versions are listed in Supplementary Note 14 and the code repository.

## Data availability

The study data and analysis release is deposited at Zenodo (version 1.1; DOI: 10.5281/zenodo.23555378) with restricted file access. Access requests should be directed to the corresponding authors; a confidential read-only link is available for editorial assessment and peer review. The deposited study data will be made publicly available upon publication of the peer-reviewed article. Version 1.1 contains the baseline archives and a dated incremental package with updated tissue-comparison analyses, source data and code. The baseline archives include the processed count matrix with cell annotations (h5ad), flow-cytometry event exports and imaging data. Raw sequencing data will be deposited in GEO/SRA before publication of the peer-reviewed article. Accession numbers and access details will be provided upon deposition. Public data analysed in this study are listed in Supplementary Table 20. Source data are provided with this preprint.

## Code availability

Analysis code is publicly available at https://github.com/Hanziwww/coch-rythum under the Apache License 2.0. An archived copy is included in the restricted-access Zenodo version 1.1 (DOI: 10.5281/zenodo.23555378).

## Use of artificial-intelligence tools

Claude (Anthropic) was used only for language polishing under the authors’ direction. Codex (OpenAI) assisted with manuscript editing and submission-material preparation.

## Acknowledgements

This work was supported by the National Natural Science Foundation of China (grant nos. 82160215 and 81660174 to Chunjie Tian; 82271169 to Yusu Ni); the Guangdong Provincial Key Laboratory of Autophagy and Major Chronic Non-communicable Diseases (grant no. 2022B1212030003 to Chunjie Tian); the Research Startup Funds for High-level Talents in the Affiliated Hospital of Guangdong Medical University (grant no. GCC20220013 to Chunjie Tian); the College Students’ Innovative Entrepreneurial Training Plan Program (grant no. S202510571025 to Chunjie Tian); and the 2025 Artificial Intelligence-Enabled Medical Device Innovation Task (Open Competition Mechanism, project no. 2025-AISJK-02007 to Chunjie Tian).

## Author contributions

Han Wang, Joe Yeong and Chunjie Tian designed the study. Han Wang performed all computational work, including data curation, formal analysis, software development and visualization, and contributed to manuscript writing. Yao Cai, Yekang Mo, Ye Tian, Weibang Chen, Dexiao Zeng, Hongyu Ning, Yue Xu, Shuangjie Dai, Xiaoyu Liu and Ruoyun He performed the experiments. Huafeng Liu, Junmin Huang, Junfeng Hao, Yusu Ni and Chunjie Tian reviewed the manuscript and provided feedback.

## Extended Data figure legends

**Extended Data Fig. 1.**
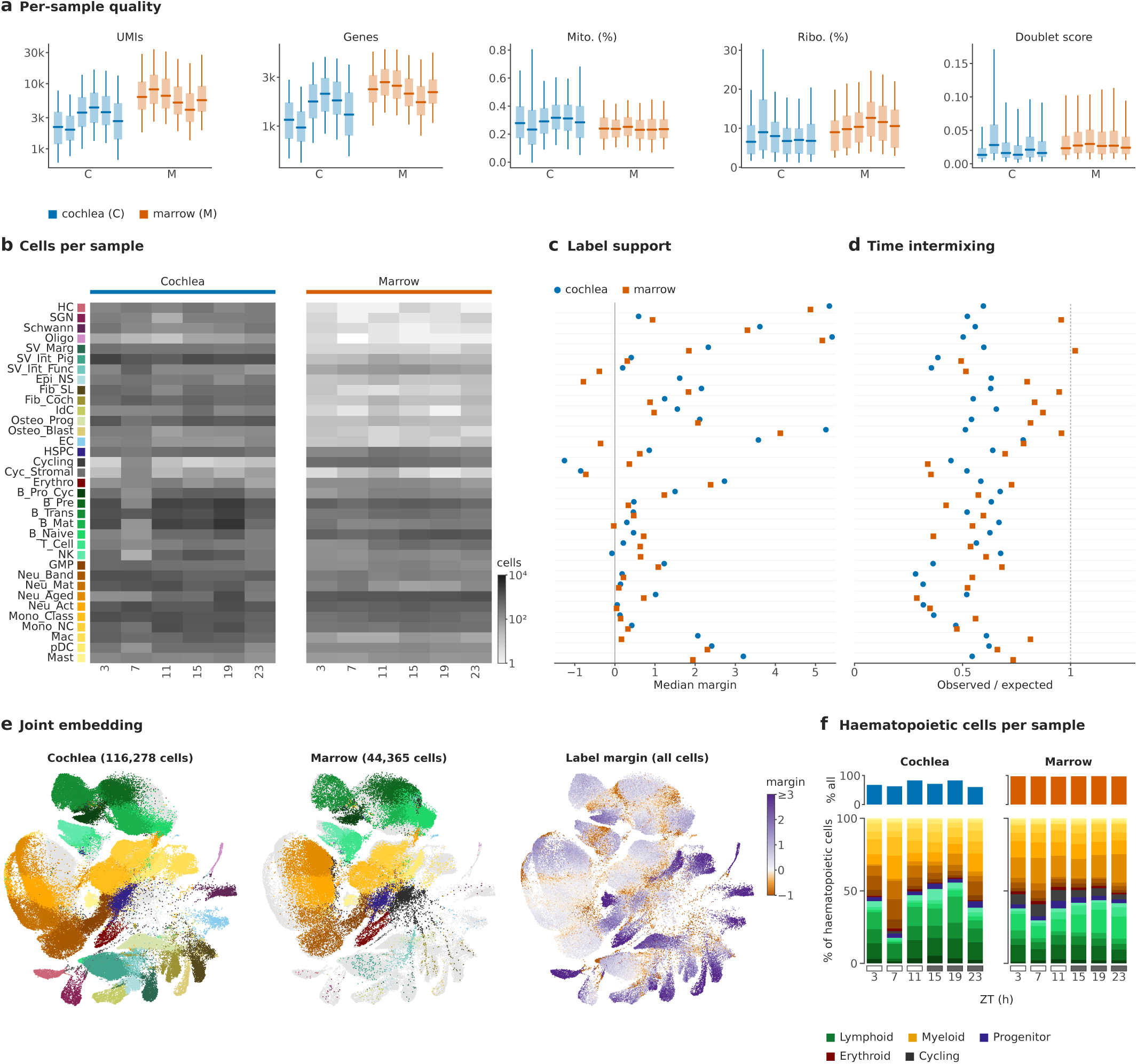
Sampling and quality control. **a**, Per-sample distributions of UMIs, detected genes, mitochondrial and ribosomal read shares and doublet score (box, interquartile range; line, median; whiskers, 5th–95th percentiles); C, cochlea; M, temporal-bone marrow (*n* = 6 samples each; 13,885–26,140 and 6,342–8,722 cells per sample). **b**, Cells of each subtype per sample (log_10_ scale), subtypes in atlas order. **c**, Label support: median margin per subtype and tissue between each cell’s score for its own label and for its best alternative label (score, mean *z*-scored expression of the label’s marker genes). Labels are the reference annotation and were not changed. **d**, Time intermixing: among each cell’s 15 nearest neighbours in the per-tissue latent space, the share of same-type neighbours from another sample, observed/expected under random mixing (1, random). **e**, Joint embedding with each tissue’s cells coloured by subtype (grey, all cells) and every cell coloured by its label margin (as in c). **f**, Haematopoietic cells per sample: share of all cells (top) and subtype composition (bottom; lymphoid below, myeloid above).

**Extended Data Fig. 2.**
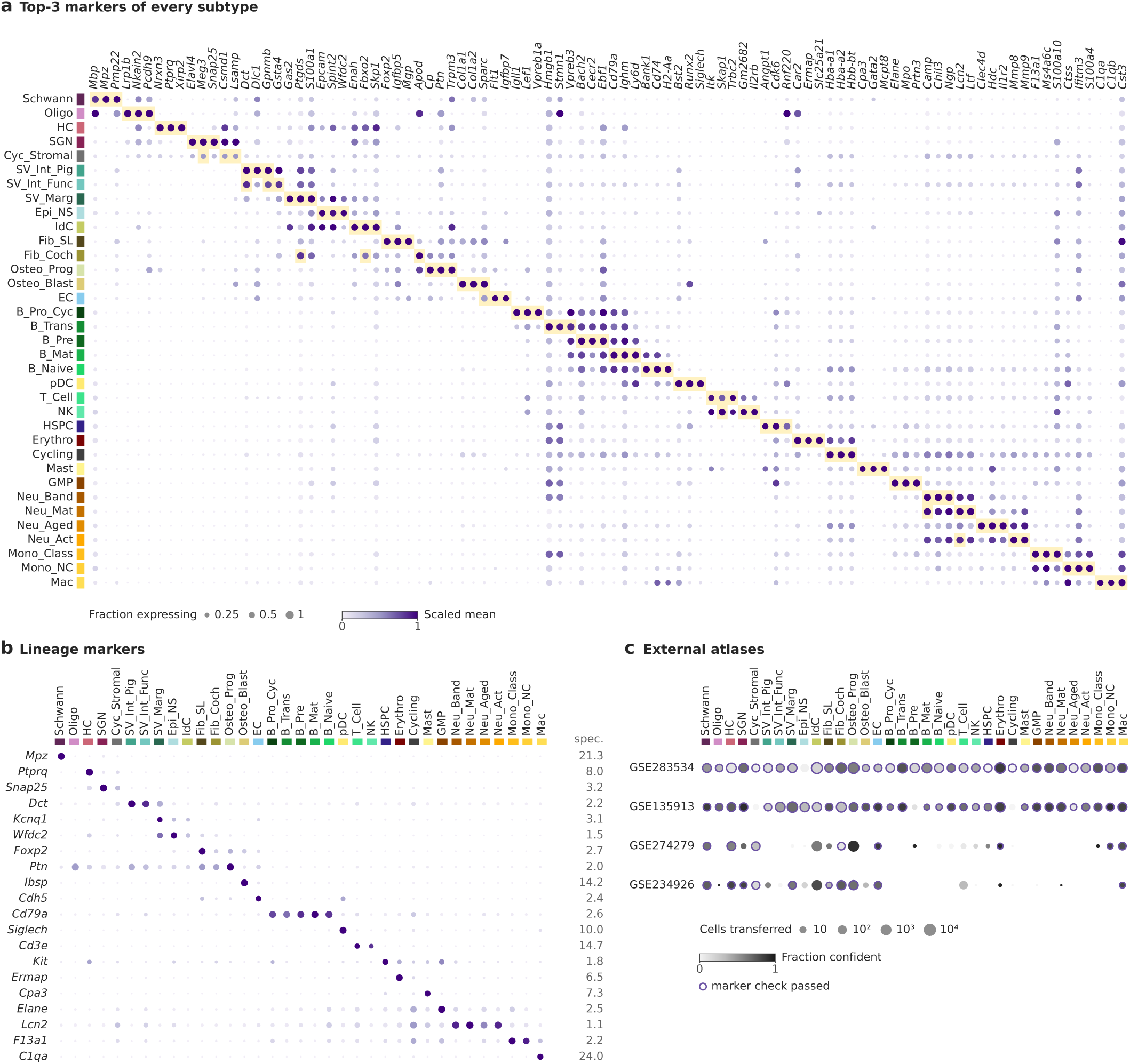
Marker atlas. **a**, The three top-ranked marker genes of every subtype (96 genes; *t*-test, log_2_ fold change *>* 1, detected in *≥* 25% of the subtype’s cells) across all 35 subtypes: dot area, fraction of cells expressing; colour, mean log-normalized expression scaled 0–1 per gene; shaded, the subtype’s own top-3 genes (genes shared by two subtypes are shaded in both rows). **b**, Lineage markers across subtypes (encoding as in a); right, lineage specificity (lowest mean among the target subtypes/highest mean among all other subtypes). **c**, Transfer of our labels to four independent adult mouse cochlear atlases (scANVI with a *k*-nearest-neighbour vote; confident, both agree with scANVI probability *≥* 0.7 and vote *≥* 0.5): dot area, cells transferred to each label; grey level, fraction transferred confidently; outline, our markers are higher in that label (*z ≥* 0.5; labels with *≥* 10 cells).

**Extended Data Fig. 3.**
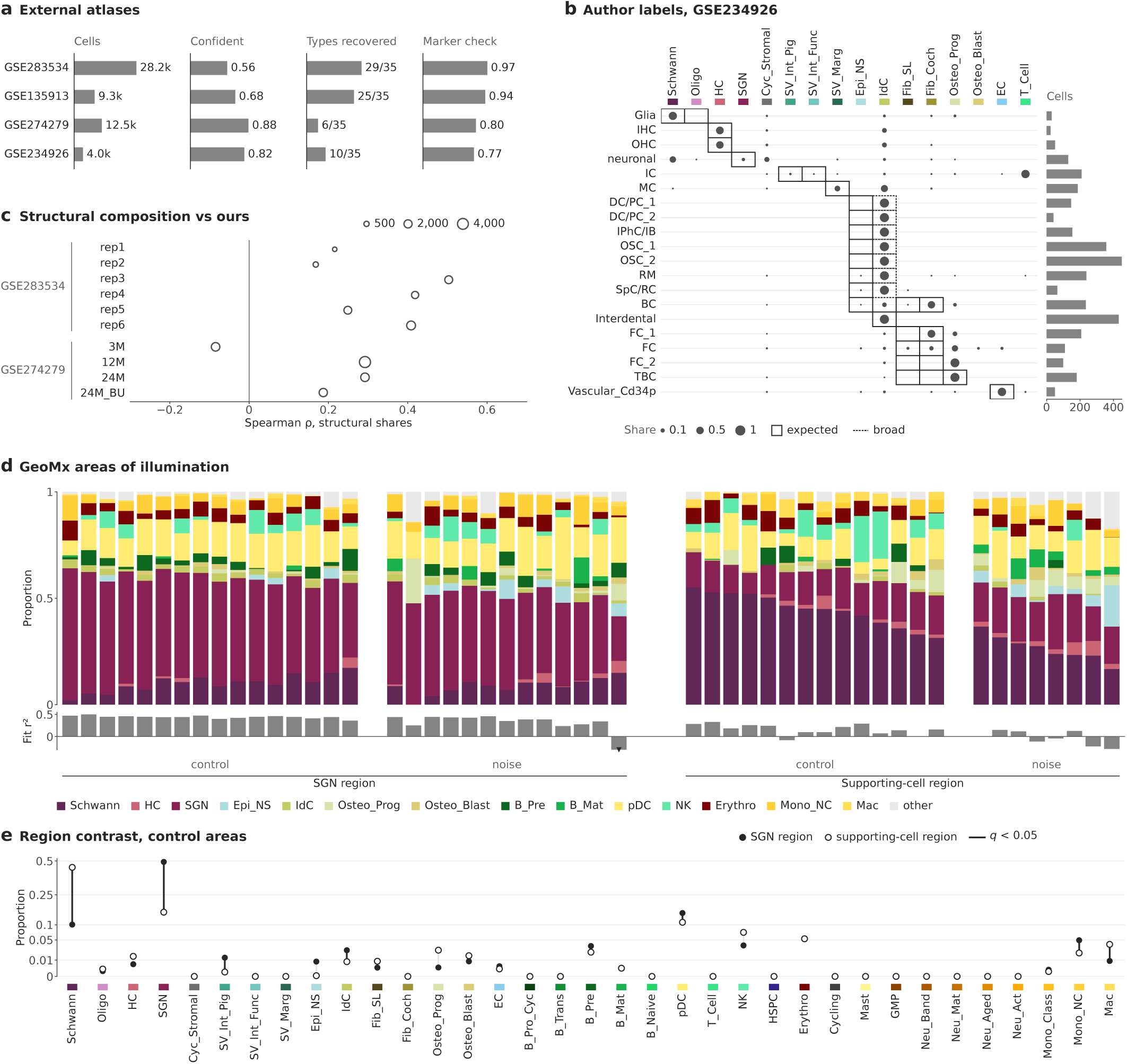
External positioning of the atlas. **a**, Four external cochlear atlases: cells after quality control, share transferred confidently, our subtypes recovered with *≥* 20 confident cells and marker-check pass rate (GSE283534, 10-week whole cochlea, 6 samples; GSE135913, adult cochlea, sorted fractions, 4; GSE274279, single-nucleus cochlea at 3, 12 and 24 months, 4; GSE234926, P90 cochlear-duct Fbxo2^+^ nuclei, 2). **b**, GSE234926 author cell types (rows; 3,442 nuclei) against our transferred labels (dot area, share of the author type); boxes, the expected label (solid) or the broader expected set (dashed); bar, nuclei per author type. **c**, Spearman *ρ* between the structural-subtype shares of each external whole-cochlea sample and our mean cochlear shares (dot area, structural cells). **d**, Public cochlear GeoMx data (GSE312646): deconvolved composition of each area of illumination by region (spiral ganglion, 16 control and 13 noise-exposed areas; supporting-cell region, 14 and 8), ordered by the region’s main subtype; subtypes with a mean share *<* 1% pooled (other). Below, weighted fit *r*^2^ of the deconvolution (triangle, one degenerate fit below the axis). **e**, Control areas: mean share of every subtype in the spiral-ganglion (filled) and supporting-cell region (open); dark links, two-sided Welch’s *t*-test with Benjamini–Hochberg *q <* 0.05.

**Extended Data Fig. 4.**
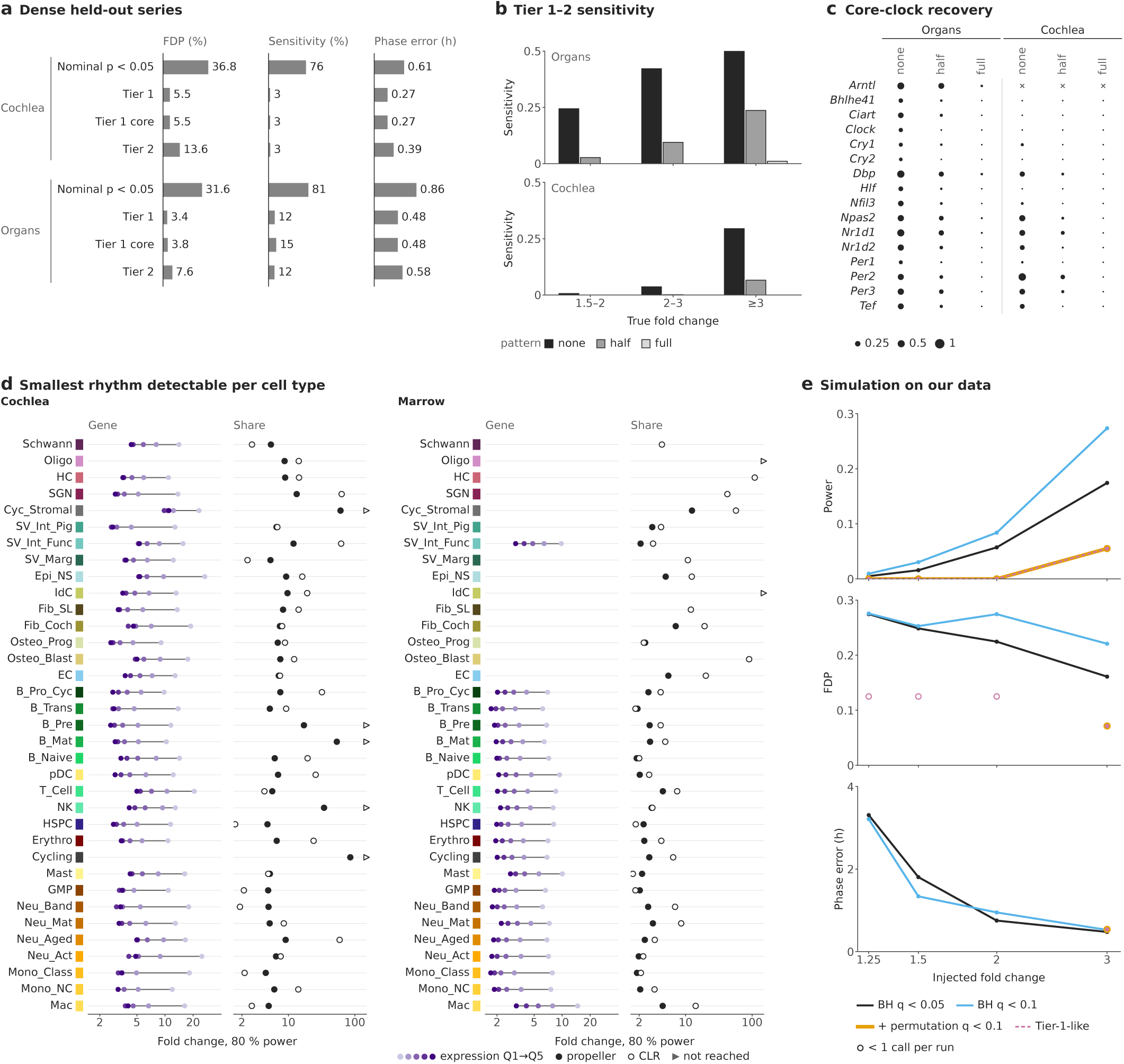
Calibration and power of the rhythm calls. **a**, Dense heldout series subsampled to our design (one sample at each of six times): GSE54650 (12 organs, constant darkness, every 2 h; 480 draws) and GSE88954 (cochlea, sham, every 4 h, *n* = 3; 40 draws), with the truth taken from the held-out samples. Realized false-discovery proportion (FDP; called genes with truth *P >* 0.2), sensitivity (truth *q <* 0.05 and peak-to-trough fold change *≥* 1.5) and median phase error per calling rule (tier definitions, Methods). **b**, Sensitivity of Tier-1/2 calls by true fold change, without and with a sample-level nuisance shaped like our cochlear sample pattern added to 40% of genes (full and half strength). **c**, Share of draws in which each core-clock gene is called Tier 1/2 (*×*, not measured). **d**, Smallest peak-to-trough fold change detectable with 80% power per cell type, for gene rhythms (moderated cosinor *F*-test, *α* = 0.001; one dot per expression quintile, Q1 lowest) and share rhythms (*α* = 0.05; filled, propeller on logit shares; open, centred log-ratio cosinor); triangles, not reached within 150-fold. **e**, Simulation on our data: sample times relabelled (removing true rhythms but keeping all sample-level structure) and rhythms of known fold change injected into 5% of genes by binomial thinning, for four cell types per tissue and three replicates; power, FDP and median phase error per rule (mean over 24 runs; open, *<* 1 call per run).

**Extended Data Fig. 5.**
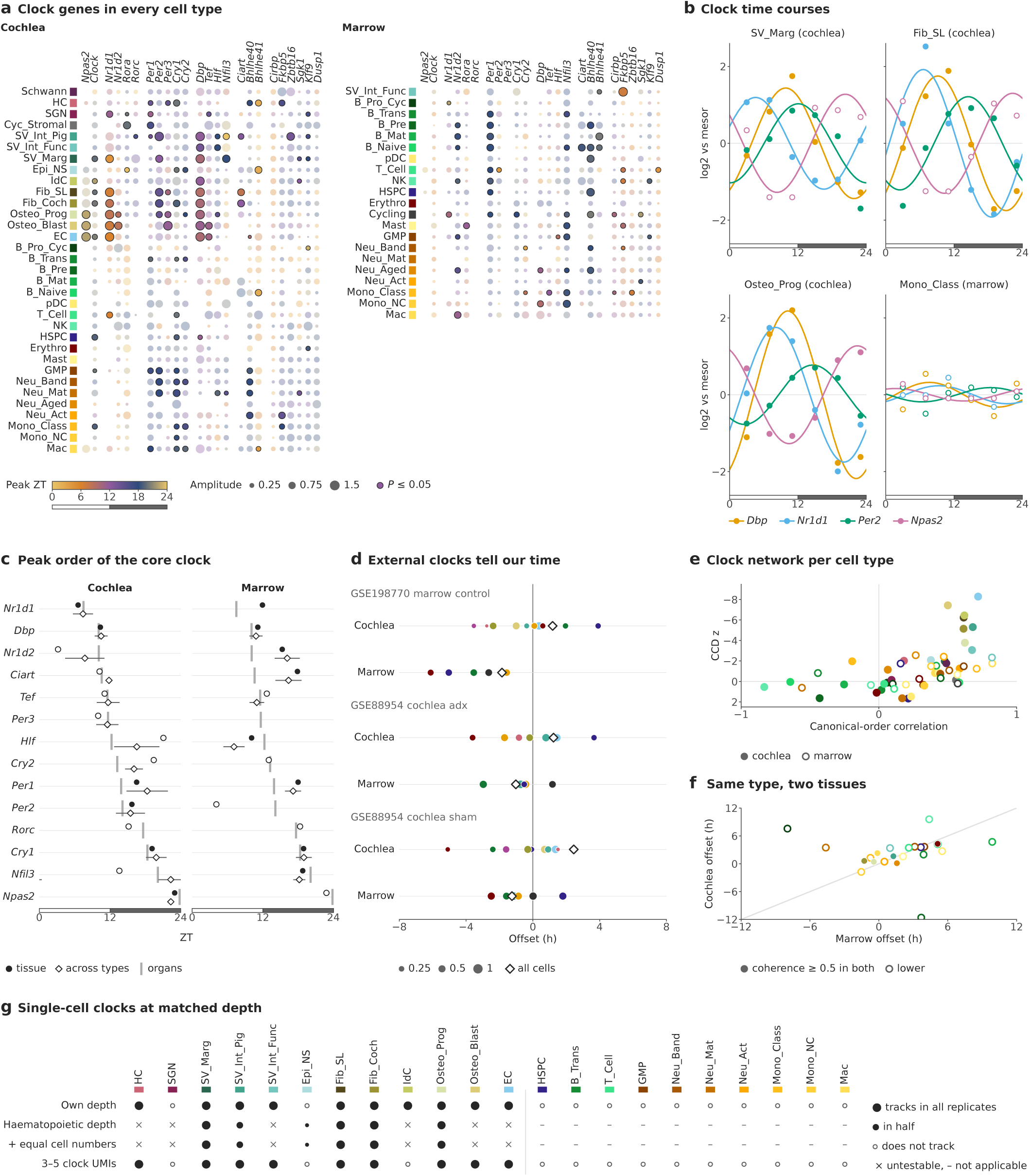
Clock detail. **a**, All 24 clock and systemic-cue genes in every primary cell type (33 cochlear, 21 marrow; *Arntl* not detected), grouped by loop: colour, peak ZT; dot area, amplitude (log_2_); outlined, exact permutation *P ≤* 0.05; blank, not expressed. **b**, Time courses of four clock genes (log_2_ CPM relative to the mesor; six samples) with cosinor fit; filled, *P ≤* 0.05. **c**, Peak of each core-clock gene: tissue pseudobulk (filled, *P <* 0.05), circular mean across cell types with *P <* 0.05 (diamond; line, circular s.d.; genes with *≥* 3 such types) and organ-atlas reference peak (GSE54650; grey). **d**, Time-telling: clock references from external data (GSE198770 marrow, control; GSE88954 cochlea, adrenalectomy or sham) predict our sampling times; offset from our ZT per class (dot area, resultant length) and for all cells (diamond). **e**, Clock network per cell type: clock correlation distance to the mouse pan-tissue reference (*z* against a null with genes permuted across samples; negative, clock-like; axis reversed) and circular correlation of the peak order with the canonical order. **f**, Clock offset of the same cell type in cochlea and marrow (23 shared types; line, identity); filled, phase coherence *≥* 0.5 in both tissues. **g**, Single-cell clock tracking (exact *P ≤* 0.05) per cochlear cell type at its own depth, at the clock-transcript depth of haematopoietic cells (20 thinnings; with and without equal cell numbers) and among cells with 3–5 clock transcripts; dot area, share of replicates; *×*, untestable.

**Extended Data Fig. 6.**
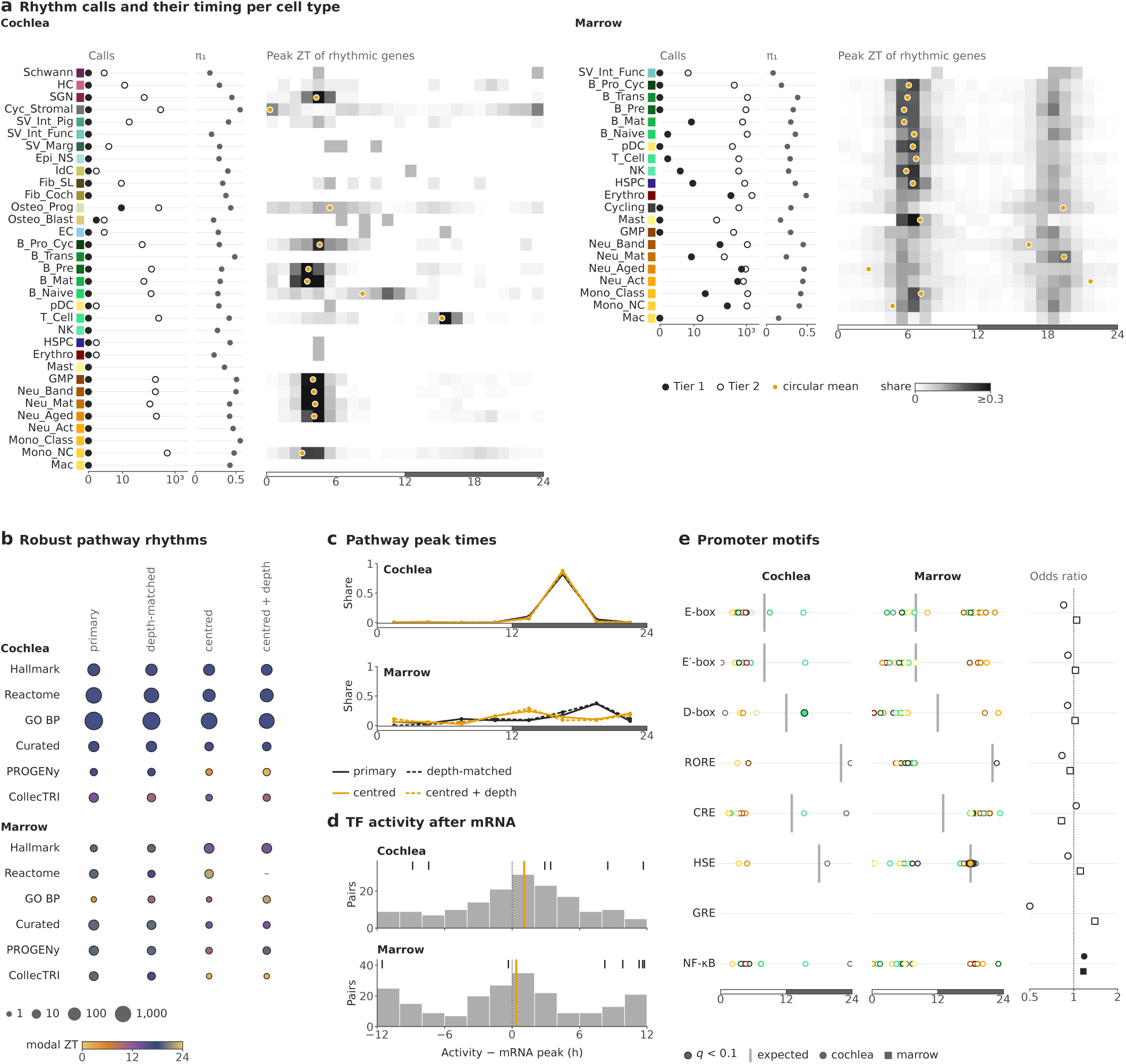
Rhythm programmes. **a**, Per primary cell type: Tier-1 (filled) and Tier-2 (open) rhythm calls (log scale), estimated share of rhythmic genes (*π*_1_, Storey) and peak-time distribution of the Tier-1/2 genes in 1-h bins (grey level, share of the type’s rhythmic genes; faded, *<* 20 genes; dot, circular mean when Rayleigh *P <* 0.05). **b**, Robust pathway and transcription factor (TF) activity rhythms (*q <* 0.1, permutation *P ≤* 0.05 and leave-one-time-point-out phase s.d. *<* 3 h) per resource and analysis version: primary; depth-matched (binomial thinning); centred on the per-sample tissue median (removing shifts shared by all cell types of a sample); both. Dot area, number of robust rhythms (log); colour, modal 3-h peak bin.**c**, Peak-time distribution of the robust pathway rhythms of each version (share per 3-h bin). **d**, TF activity (CollecTRI) minus TF mRNA peak for TF *×* cell-type pairs with both rhythms *P <* 0.05; line, circular mean lag of non-clock TFs; ticks, core-clock TFs. **e**, Promoter motifs (1 kb upstream): circular mean peak of rhythmic motif carriers per cell type (filled, directional permutation *q <* 0.1 towards the expected phase; grey tick, expected phase: E-box and E*^′^*-box ZT8, D-box ZT12, CRE ZT13, HSE ZT18, RORE ZT22) and tissue-level enrichment among rhythmic genes (Cochran–Mantel–Haenszel odds ratio over cell type *×* expression decile; filled, *q <* 0.05).

**Extended Data Fig. 7.**
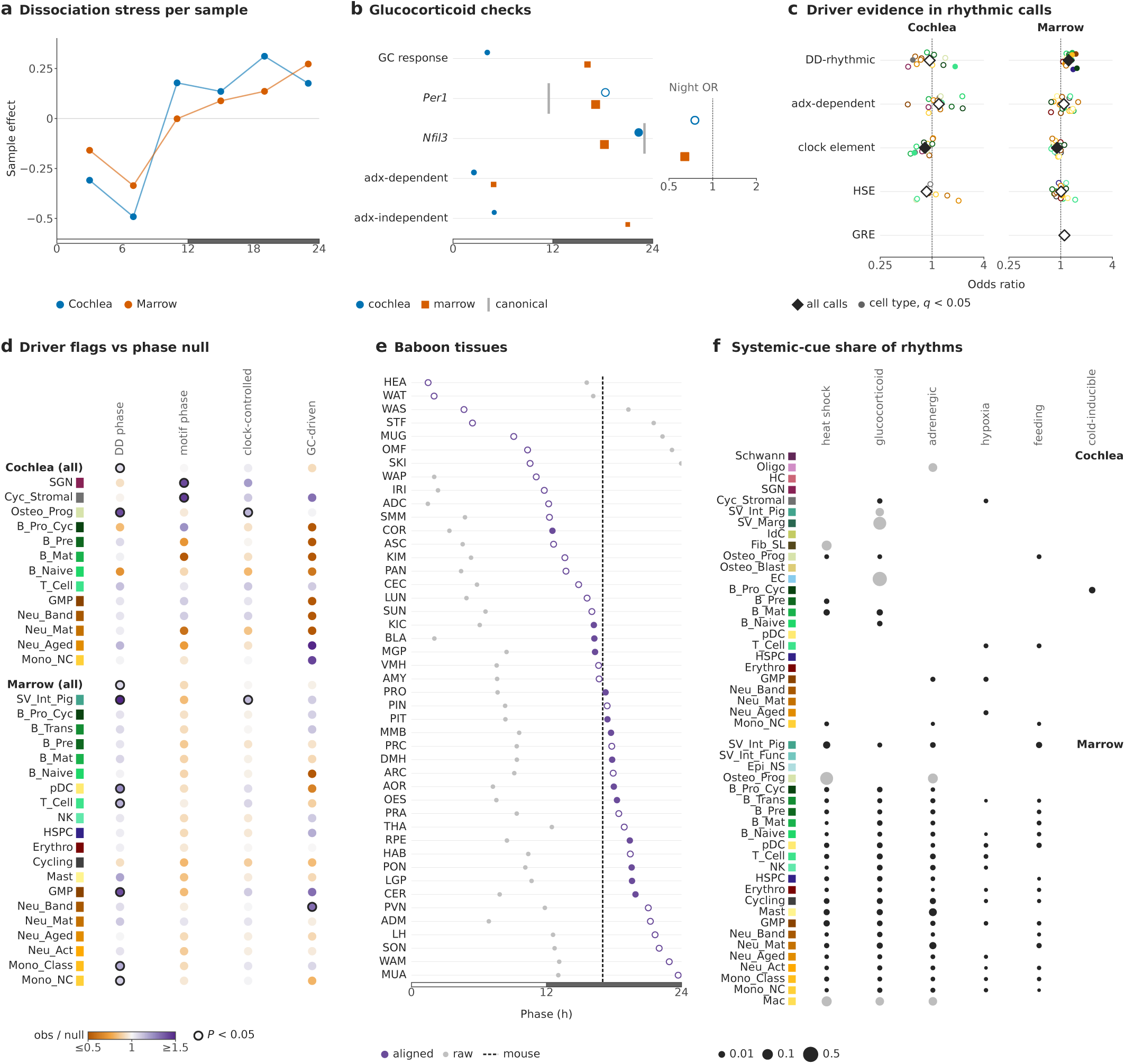
Heat shock and rhythm drivers. **a**, Per-sample dissociation-stress effect (sample term of a model of each cell type’s dissociation-stress profile). **b**, Gluco-corticoid (GC) checks: circular mean peak of curated GC-response genes, of *Per1* and *Nfil3*, and of rhythmic genes that lose (adx-dependent) or keep (adx-independent) their rhythm after adrenalectomy in an independent cochlear series (GSE88954); dot area, resultant length; filled, Rayleigh *P <* 0.05; grey tick, canonical peak. Right, odds that night-peaking rhythmic genes are adx-dependent relative to day-peaking ones (Fisher’s exact test). **c**, Driver evidence among rhythm calls versus expression-matched non-rhythmic genes (Cochran–Mantel–Haenszel odds ratio over expression deciles): rhythmic in constant darkness in *≥* 2 organs (GSE54650), adx-dependent, promoter clock element, heat-shock element (HSE) and GC-response element; dots, cell types (filled, *q <* 0.05); diamond, all calls. **d**, Phase-dependent driver flags of Tier-1/2 calls (peak within 4 h of the constant-darkness phase or of the phase expected from a promoter clock element, either, or GC-driven) relative to a null with peaks shuffled within each cell type (200 draws): colour, observed/null share; outlined, permutation *P <* 0.05. **e**, Heat-shock programme phase in 45 baboon tissues^40^ (tissue codes as in that study): clock-aligned phase (purple; filled, *P <* 0.05) and raw ZT (grey); dashed line, mouse heat-shock phase (aligned). **f**, Share of each cell type’s rhythmic genes that belong to a systemic-cue programme and peak in that programme’s common phase (dot area; faded, *<* 20 rhythmic genes).

**Extended Data Fig. 8.**
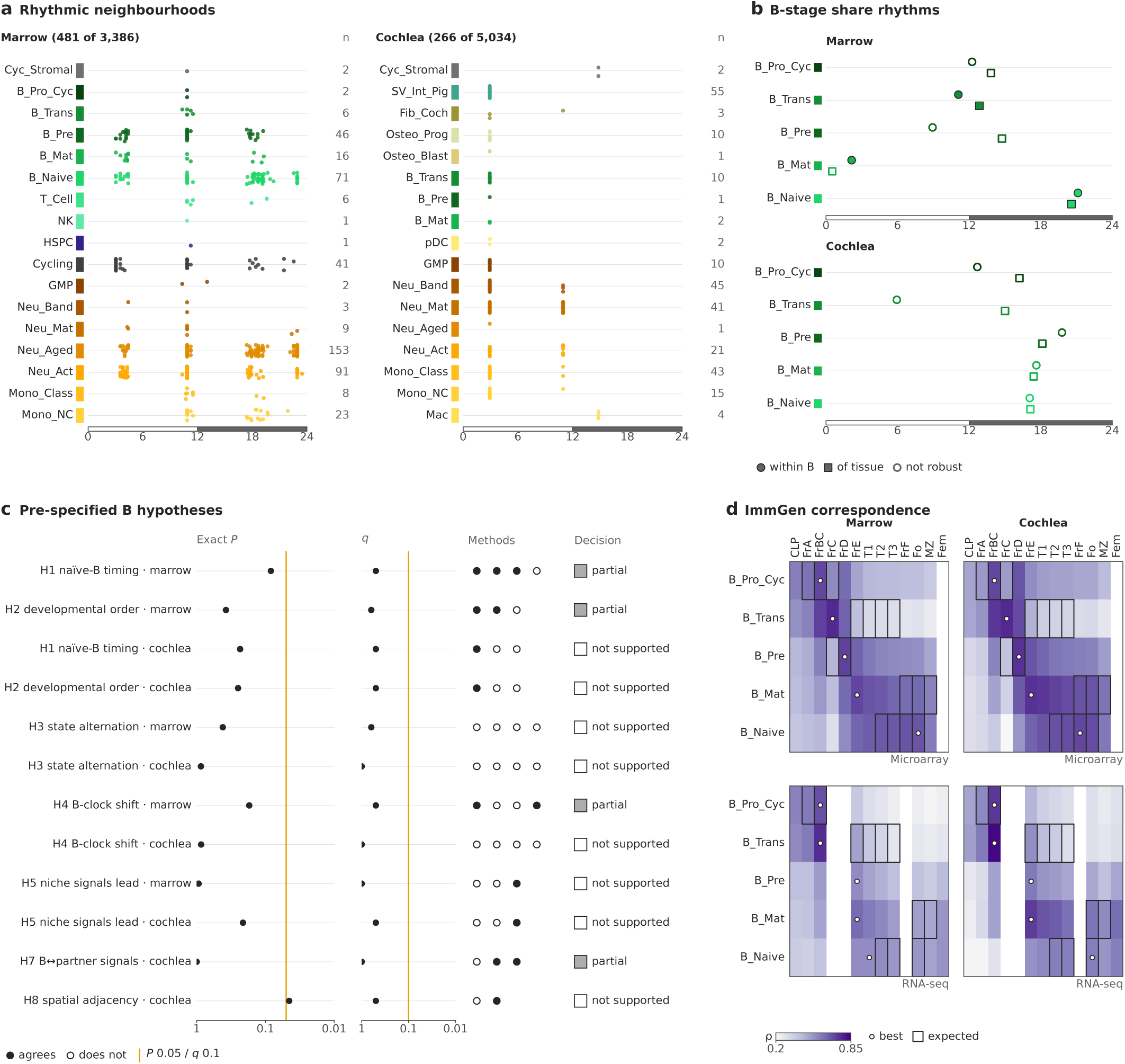
Marrow neighbourhoods and the B lineage. **a**, Milo neighbourhoods with a 24-h rhythm in abundance (spatial FDR *<* 0.1; scANVI embedding): peak ZT of each neighbourhood by its majority cell type; right, number of rhythmic neighbourhoods. **b**, B-stage share rhythms within the B lineage (circle) and as share of the tissue (square); filled, multi-model verdict robust or supported. Stages in ImmGen order, our labels with their ImmGen equivalents: Pro-B (FrB/C), Transitional (large pre-B, FrC*^′^*), Pre-B (small pre-B, FrD), Mature (immature, FrE/T1), Naive (recirculating, FrF/follicular). **c**, The 12 pre-specified B-lineage hypotheses: exact permutation *P* of the primary statistic, Benjamini–Hochberg *q* across the primary set, independent method lines (filled, agrees) and decision (supported, *q <* 0.1 and *≥* 2 agreeing methods; partial, one criterion met). **d**, Spearman correlation between our B-stage pseudobulk profiles and ImmGen B-cell populations (microarray, GSE15907; RNA-seq, GSE109125; marrow and cochlea): dot, best match per stage; box, expected population.

**Extended Data Fig. 9.**
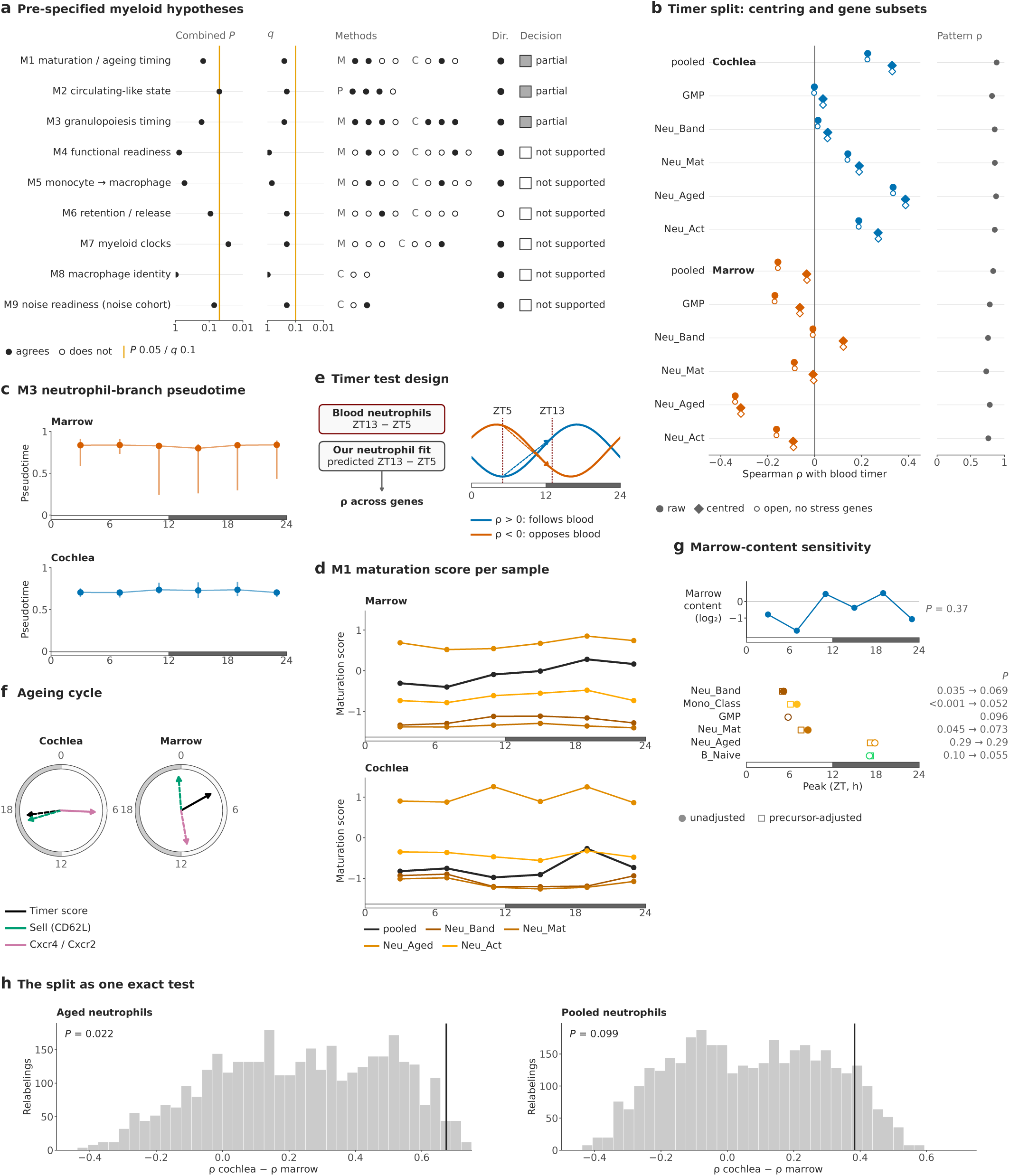
Myeloid hypotheses and timer robustness. **a**, The nine pre-specified myeloid hypotheses: combined exact *P* (Fisher’s method over tissues), Benjamini–Hochberg *q*, independent method lines per tissue (M, marrow; C, cochlea; P, paired; filled, agrees), pre-specified direction (filled, met) and decision (Supplementary Note 13). **b**, Spearman *ρ* of each neutrophil state’s predicted change (ZT13 *−* ZT5) with blood neutrophils (GSE86619), raw and within-sample centred, with and without stress and immediate-early genes; right, correlation with the change implied by the sample-level pattern alone. **c**, Per-sample median neutrophil-branch pseudotime of GMP, band and mature neutrophils (bar, interquartile range). **d**, Per-sample median maturation score ^42^ per neutrophil group (marrow above cochlea). **e**, Design of the timer test (idealized curves): each gene’s blood day–night change (ZT13 *−* ZT5) against the change that a neutrophil state’s cosinor fit predicts (Spearman *ρ* across genes). **f**, Exploratory: peak phase of the timer score, *Sell* (CD62L) and the *Cxcr4* /*Cxcr2* ratio in cochlear and marrow neutrophils (solid, permutation *P <* 0.05). **g**, Marrow content of the cochlear samples (sensitivity analysis). Top, log_2_ ratio of marrow-resident precursors to structural cells per sample (cosinor *P*). Bottom, peak of each cochlear share unadjusted (circle; primary) and adjusted (square); filled, *P <* 0.05; right, *P* unadjusted *→* adjusted. **h**, The split as one statistic, *ρ* (cochlea) *− ρ* (marrow), against all 3,600 joint relabellings for aged and pooled neutrophils (line, observed; one-sided exact *P*; with phase-shifted labels, 720 *×* 720, *P* = 0.0073 and 0.069). *n* = 6 pooled samples per tissue.

**Extended Data Fig. 10.**
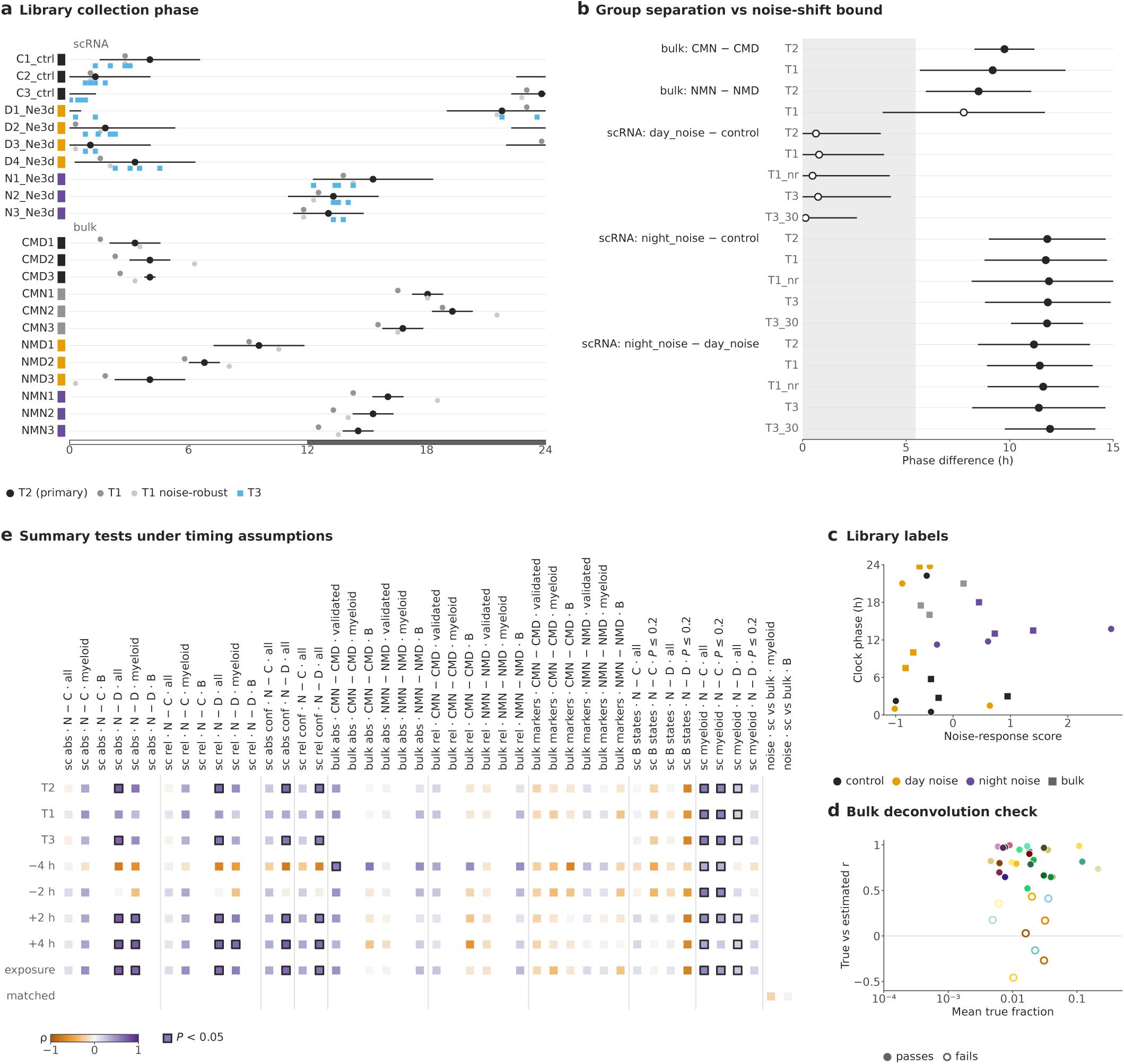
The independent cohort in detail. **a**, Inferred collection phase (our ZT) of every library of the independent cohort ^28^ (single-cell, CRA019820: controls C1–C3, day noise D1–D4, night noise N1–N3; bulk, CRA019827: CMD, CMN, NMD, NMN, *n* = 3 each) under each clock estimator: T2 (primary; template from an independent sham-cochlea series, GSE88954; bar, 95% interval), T1 (template from our cochlear clock fits), noise-robust T1 (clock genes with a time-matched noise effect removed) and composition-robust T3 variants (per structural cell type with *≥* 50 or *≥* 30 cells per library). **b**, Phase separation of each group contrast under each estimator (bar, 95% interval) against the upper 95% bound of a noise-associated clock shift (grey; Methods). **c**, Label check: noise-response score and clock phase of every library, coloured by its label. **d**, Bulk deconvolution validated on pseudobulk mixtures of the cohort’s single cells: correlation of true and estimated fractions per label against its mean fraction (filled, passes). **e**, Exploratory summary tests (Spearman *ρ* between observed and predicted change) under each timing assumption: inferred phase (T2), estimator T1 or T3, global offsets of *−*4, *−*2, +2 and +4 h, the exposure-time assumption and the time-matched noise-response comparison. Rows: data (sc, single cell; abs/rel, absolute or immune-relative shares; conf, confidently mapped cells; markers, bulk marker signatures; states), contrast (N *−* C, night noise *−* control; N *−* D, night *−* day noise; CMN *−* CMD and NMN *−* NMD, bulk) and scope. Colour, *ρ*; outlined, one-sided permutation *P <* 0.05.

