## Supplementary Information for "An anatomical and day–night atlas links cochlear immunity to adjacent bone marrow"

#### Contents

**Supplementary figures** **2**

**Supplementary Notes** **10**

**Supplementary Tables** **27**

### Supplementary figures

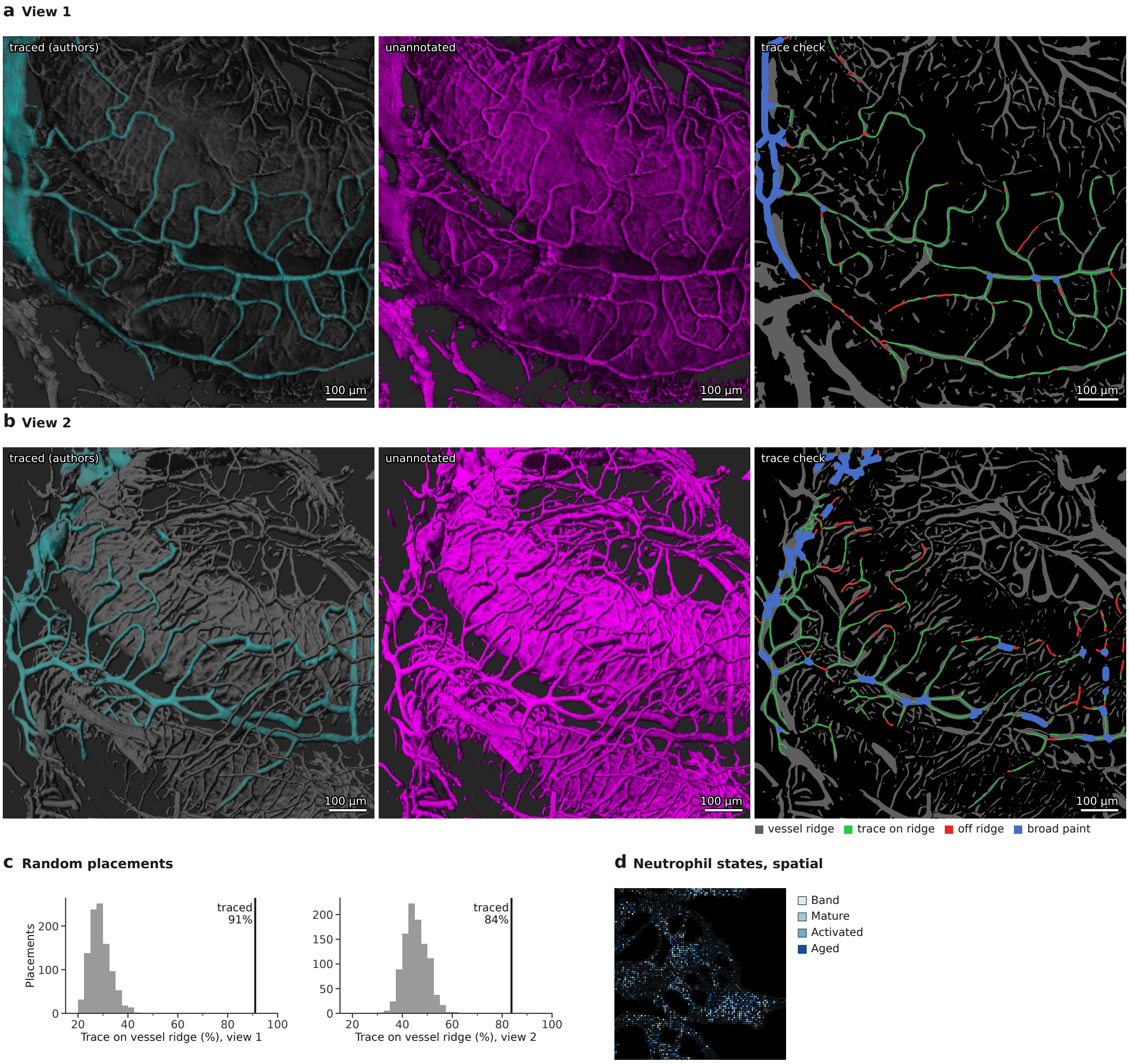


**Supplementary Fig. 1 | Check of the vessel tracing between the temporal-bone marrow and the cochlea.** **a**, View 1 of the three-dimensional render (as Fig. 4a): left, the authors' hand tracing of vessels (teal) over a grey copy of the render; middle, the same view unannotated (vessel signal, magenta); right, the check: vessel ridges detected in the unannotated render without reference to the tracing (grey), centrelines of thin traces on a ridge (green) or off a ridge (red), and broad paint over wide vessels or bone, not tested (blue). The two renders registered without shift. **b**, View 2, as in a. **c**, Share of thin-trace centreline within 2 pixels of a vessel ridge for 999 random rotations and translations of the same traces over the rendered tissue (histograms) and for the tracing itself (lines): 91% in view 1 and 84% in view 2, against means of 29% and 45% for random placements ($P = 0.001$ each); the longest stretch off a ridge was 37 and 63 $\mu$m. **d**, Spatial transcriptomics map of the deconvolved neutrophil states (as Fig. 4c). Scale bars as labelled.

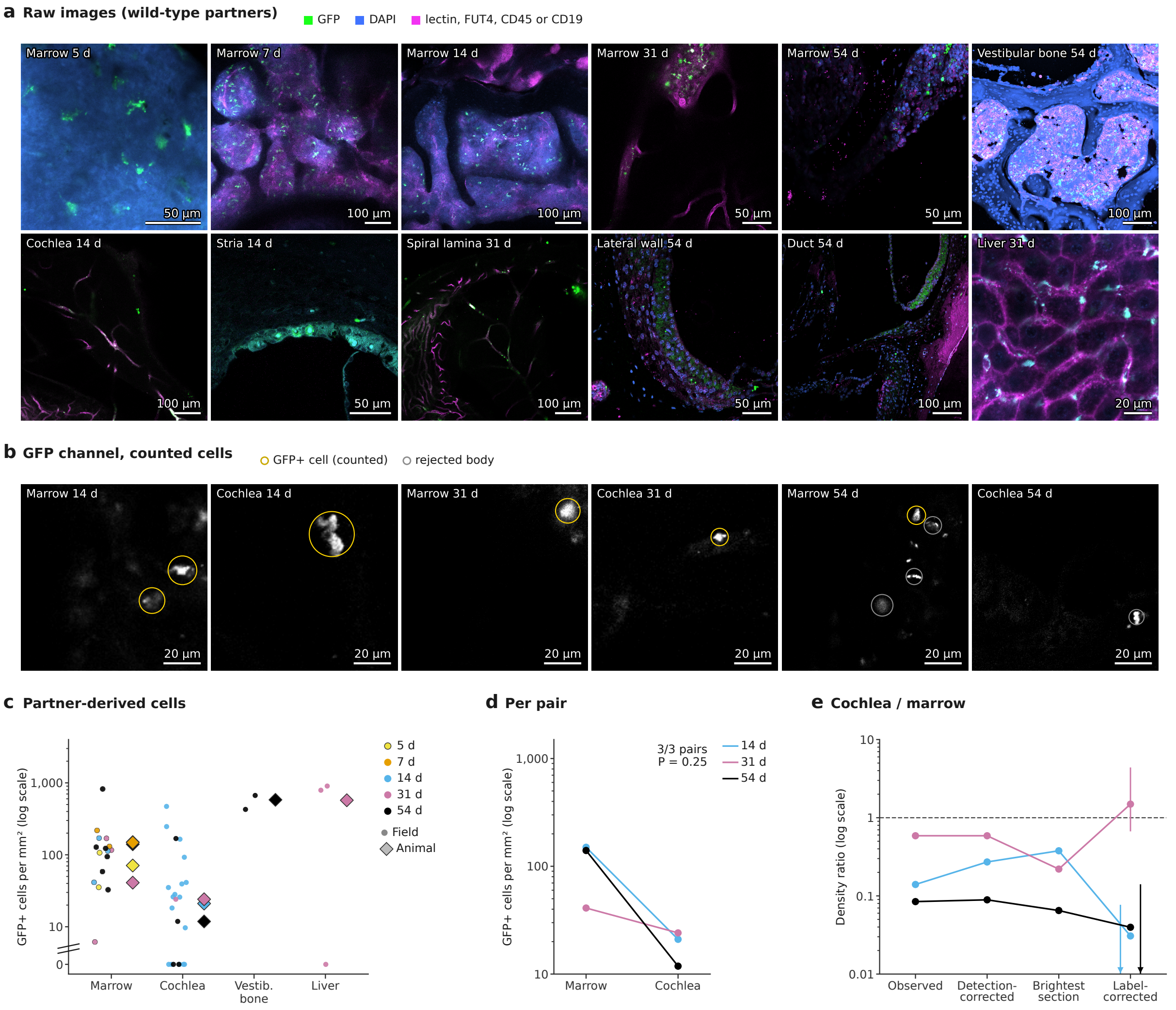


**Supplementary Fig. 2 | Partner-derived cells across the tissues of wild-type parabionts.** Wild-type mice were joined to Lyz2-EGFP partners, one pair per pairing duration; all images are from the wild-type partners, in which GFP marks partner-derived myeloid cells. **a**, Raw images: one optical section per animal and tissue (for z-stacks, the analysed section with the strongest GFP signal) of the temporal-bone marrow (5, 7, 14, 31 and 54 days), vestibular bone (54 days), cochlea (14 days, cochlear section and stria vascularis; 31 days, osseous spiral lamina; 54 days, lateral wall and cochlear duct) and liver (31 days). GFP, green; DAPI, blue; lectin, FUT4, CD45 or CD19, magenta; linear display per channel. GFP settings differed between samples, so intensities are not compared between images. **b**, GFP channel of 100-$\mu$m crops of the marrow and cochlea of the 14-, 31- and 54-day pairs; circles, $GFP^+$ bodies kept as partner-derived cells by the upgraded cell rule (yellow) or rejected by it (grey). **c**, Partner-derived cells per $mm^2$ of tissue per field (points; colour, pairing duration; fields without a counted cell at the axis floor) and pooled per animal (diamonds); 16 marrow, 22 cochlear, 2 vestibular-bone and 3 liver fields. **d**, Pooled partner-derived cells per $mm^2$ of marrow and cochlea of every pair with both compartments imaged. The cochlea was lower in 3 of 3 pairs (exact sign test, pairs as the unit, $P = 0.25$). **e**, Cochlea-to-marrow ratio per pair as observed, corrected for GFP detection, in each acquisition's brightest optical section (below 1 in every pair, 0.06–0.59) and corrected with the blinded visual labels (vertical lines, bootstrap 95% CI; arrowheads, intervals reaching 0): 14 days 0.03 (0–0.08), 31 days 1.49 (0.67–4.39), 54 days 0.04 (0–0.14). Scale bars as labelled.

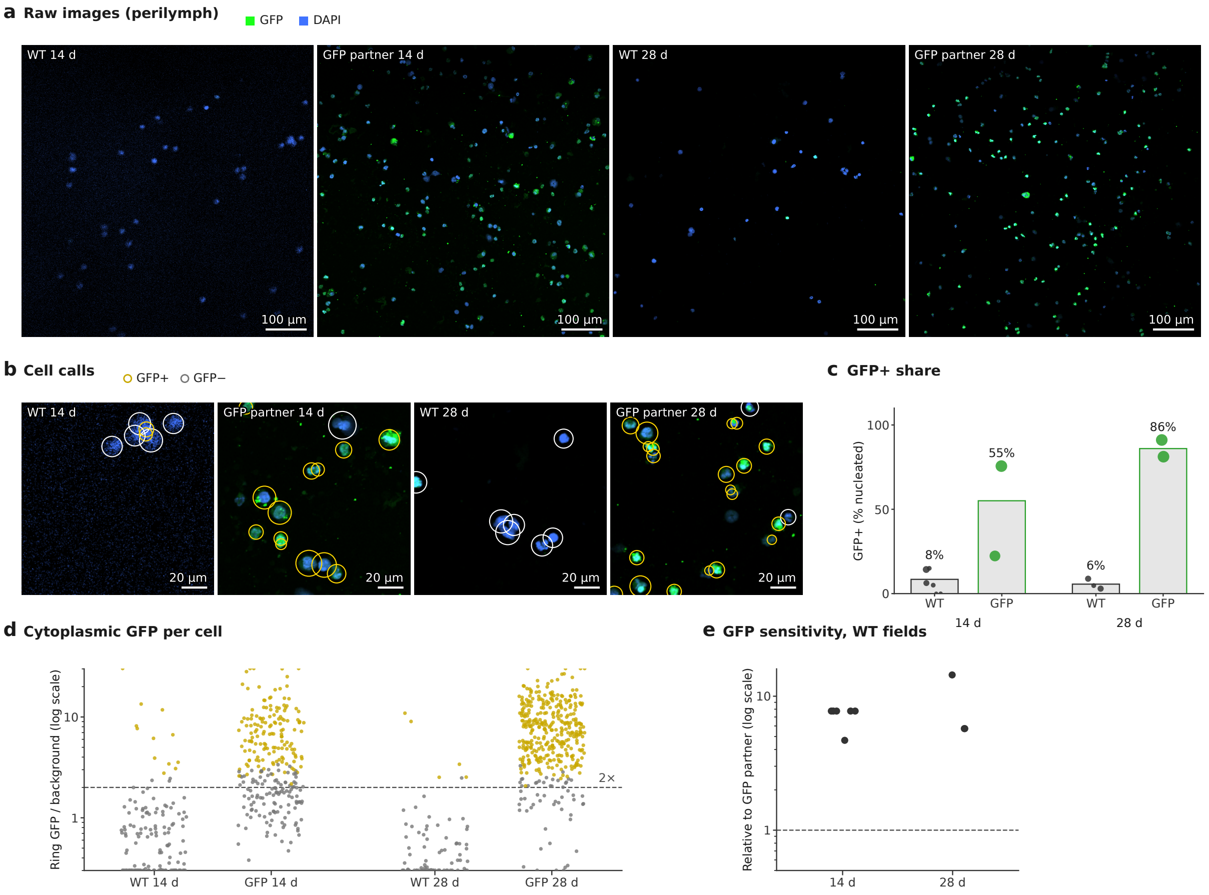


**Supplementary Fig. 3 | Perilymph cells of wild-type and GFP partners.** **a**, Raw images: full fields of perilymph preparations of the wild-type and the GFP (Lyz2-EGFP) partners of the 14- and 28-day pairs (GFP, green; DAPI, blue), GFP shown with one display range for both partners of a pair. **b**, 150-$\mu$m crops (display as in a) with the nucleated cells called GFP$^+$ (yellow circles) or GFP$^-$ (white circles); cytoplasmic GFP counts, signal confined to the nucleus does not. **c**, GFP$^+$ share of nucleated cells per field (points; size, nucleated cells) and pooled (bars): wild-type partners 8.5% (14 days; 6 fields, 142 cells) and 5.6% (28 days; 3 fields, 89 cells); GFP partners 55% (2 fields, 267 cells) and 86% (2 fields, 393 cells). **d**, Ring GFP over local background for every nucleated cell of c; dashed line, twice the background; yellow, cells called GFP$^+$. **e**, GFP detection sensitivity of each wild-type field relative to the same pair's GFP-partner fields: 4.7–14.4-fold, so the low wild-type shares do not reflect weaker detection. Fields imaging the same position were counted once. Scale bars as labelled.

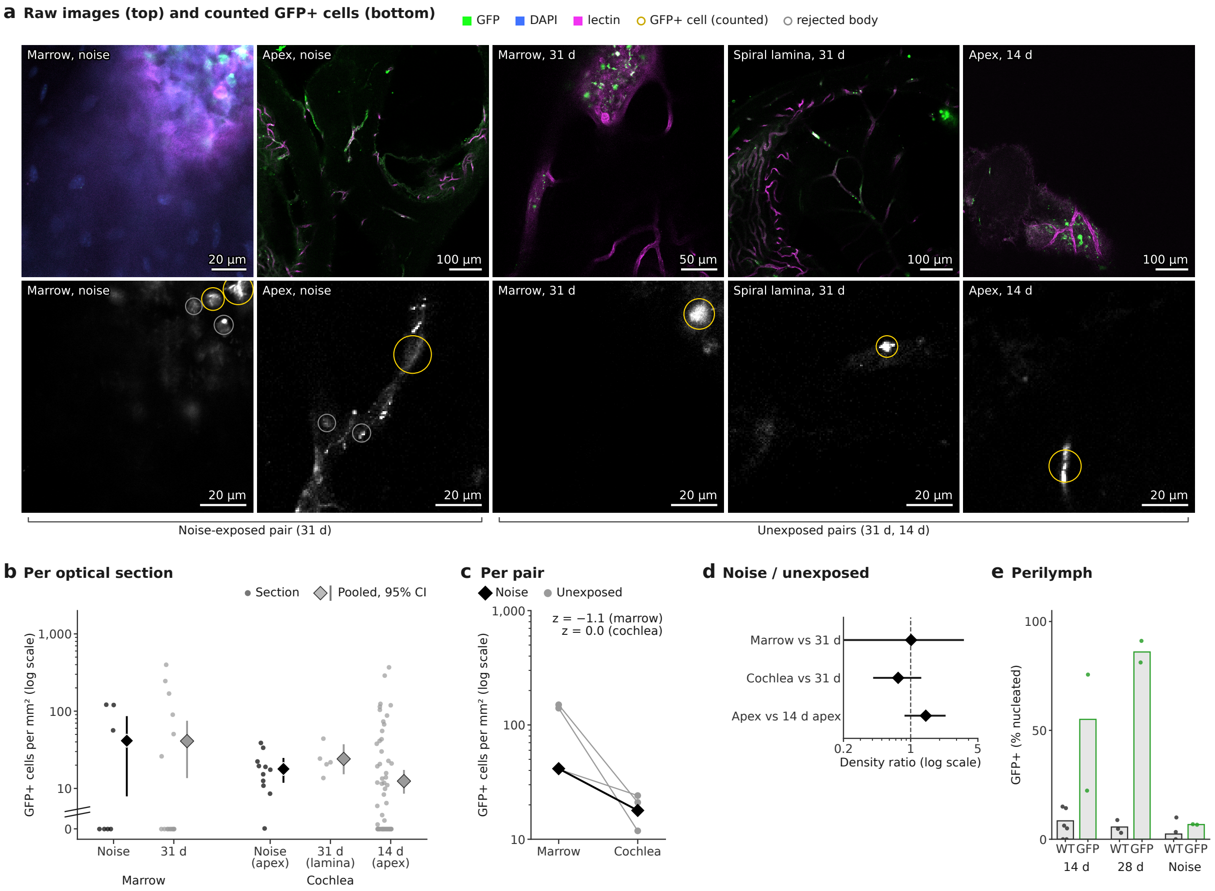


**Supplementary Fig. 4 | Partner-derived cells after noise exposure.** One 31-day pair was exposed to noise before tissue collection (Methods). This unreplicated comparison with unexposed pairs cannot test a noise effect. Counts by the upgraded cell rule, as in Fig. 4g. **a**, Raw images (top; full optical sections; GFP, green; DAPI, blue; lectin, magenta) and GFP channel of 100-$\mu$m crops of the same sections (bottom; circles, GFP$^+$ bodies kept as cells in yellow, rejected in grey): marrow and cochlear apex of the noise-exposed wild-type partner; marrow and osseous spiral lamina of the unexposed 31-day partner (same pairing duration); cochlear apex of the unexposed 14-day partner (same cochlear region). **b**, Partner-derived cells per mm$^2$ per analysed optical section (points; sections without a counted cell at the axis floor) with the pooled density and its 95% bootstrap interval (diamond and bar): marrow, noise 41 (8–84) versus unexposed 41 (14–73); cochlea, noise apex 18 (12–24) versus 31-day spiral lamina 24 (16–36) and 14-day apex 12 (9–17). **c**, Pooled densities per pair: the noise-exposed pair (black) among the unexposed pairs (grey). Its cochlear density lay at the mean of the unexposed pairs (log scale, $z = -0.04$) and its marrow density within their range ($z = -1.1$), both inside the 95% prediction interval for one unexposed pair; its cochlea-to-marrow ratio was 0.43 (unexposed pairs 0.08–0.59). **d**, Noise over unexposed density ratios with 95% bootstrap intervals: marrow 1.01 (0.14–3.50); cochlea against the same pairing duration 0.74 (0.42–1.26); cochlear apex against the same region 1.44 (0.88–2.26). The intervals describe sections within one animal per condition, not variation between animals. **e**, Perilymph GFP$^+$ share of every partner (bars, pooled; points, fields): GFP detection failed in the noise session (its GFP partner scored 7%, against 55–86% in the other GFP partners), so the noise pair's perilymph is not interpreted. This single pair provided no evidence of increased recruitment of circulating GFP$^+$ cells to the cochlea or marrow after noise. Scale bars as labelled.

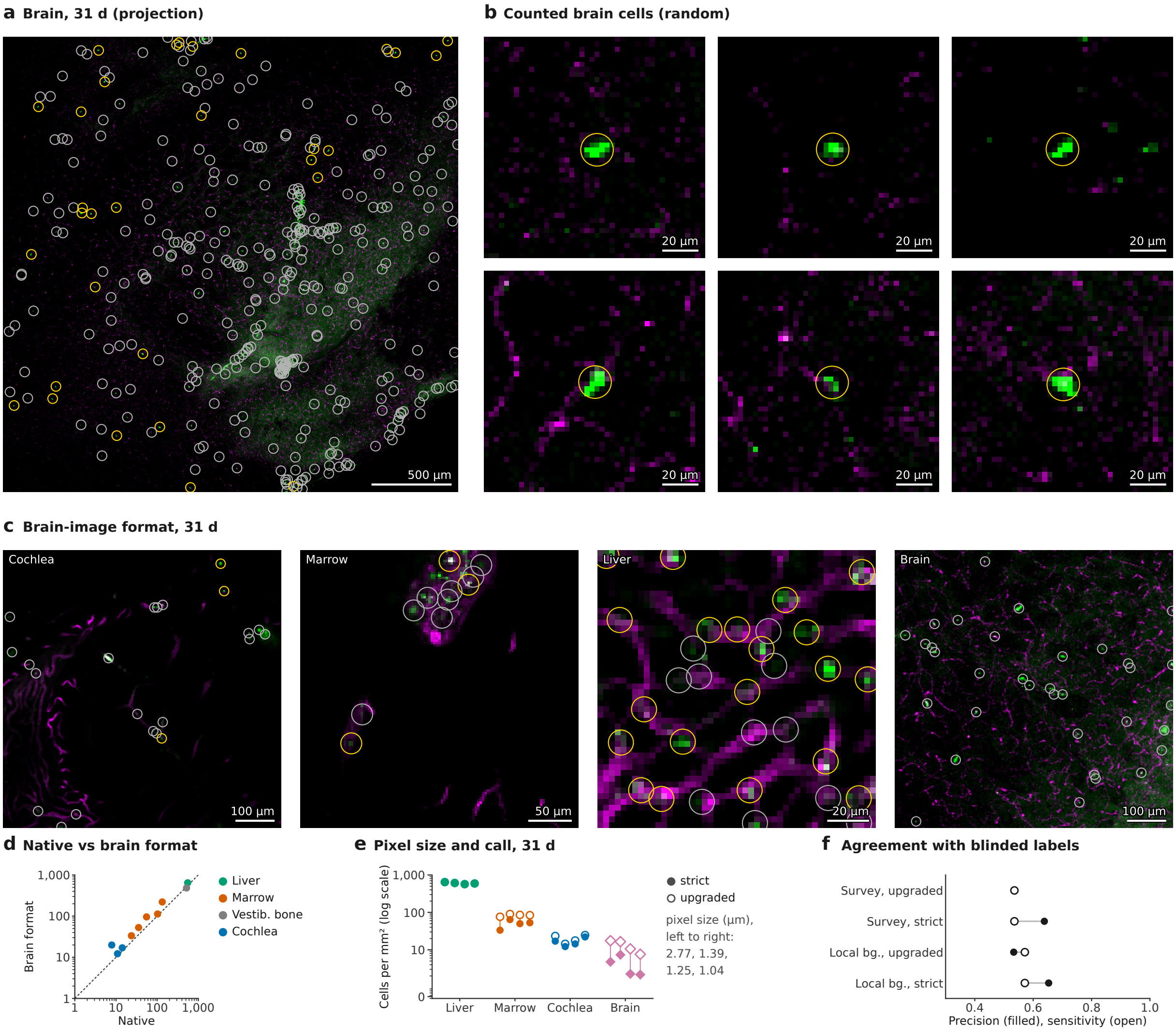


**Supplementary Fig. 5 | The brain, the brain-image format and the stricter cell rule.** The only brain image of the parabiosis collection is a rendered maximum projection of the 31-day wild-type partner without DAPI or pixel size; it was counted at the collection's usual overview pixel size (2.773 µm), and every other tissue was converted to its format, so that all tissues were counted alike: one optical section at a time, box-averaged to the same pixel size, GFP and lectin only. $GFP^+$ bodies were detected on the local background (fixed before its run), and a stricter cell rule, chosen from a fixed grid by agreement with the blinded visual labels, was applied to every tissue (Methods). **a**, Brain (GFP, green; lectin, magenta); circles, $GFP^+$ bodies counted as partner-derived cells (yellow) or not (grey). **b**, Six counted brain cells drawn at random; 120- µm crops. **c**, Cochlea (osseous spiral lamina), marrow and liver of the 31-day partner in the brain-image format, and the central 710- µm window of the brain (the field size of the cochlear section). **d**, Partner-derived cells per $mm^2$ by the stricter rule, native against the brain-image format, per animal and tissue; line, identity. **e**, The 31-day partner in the brain-image format by assumed pixel size (left to right 2.773, 1.387, 1.248 and 1.040 µm): filled, stricter rule; open, upgraded criteria. **f**, Agreement with 152 blinded labels in the brain-image format: precision (filled) and sensitivity (open) of the global-threshold (survey) and the local-background detector, each with the upgraded and the stricter call. The brain rendering saturates (its bodies reach the 8-bit ceiling); local-background detection raised its count from 2.1 to 4.8 cells per $mm^2$ (stricter rule) but cannot recover intensities above the ceiling, so it may remain an underestimate. Scale bars as labelled.

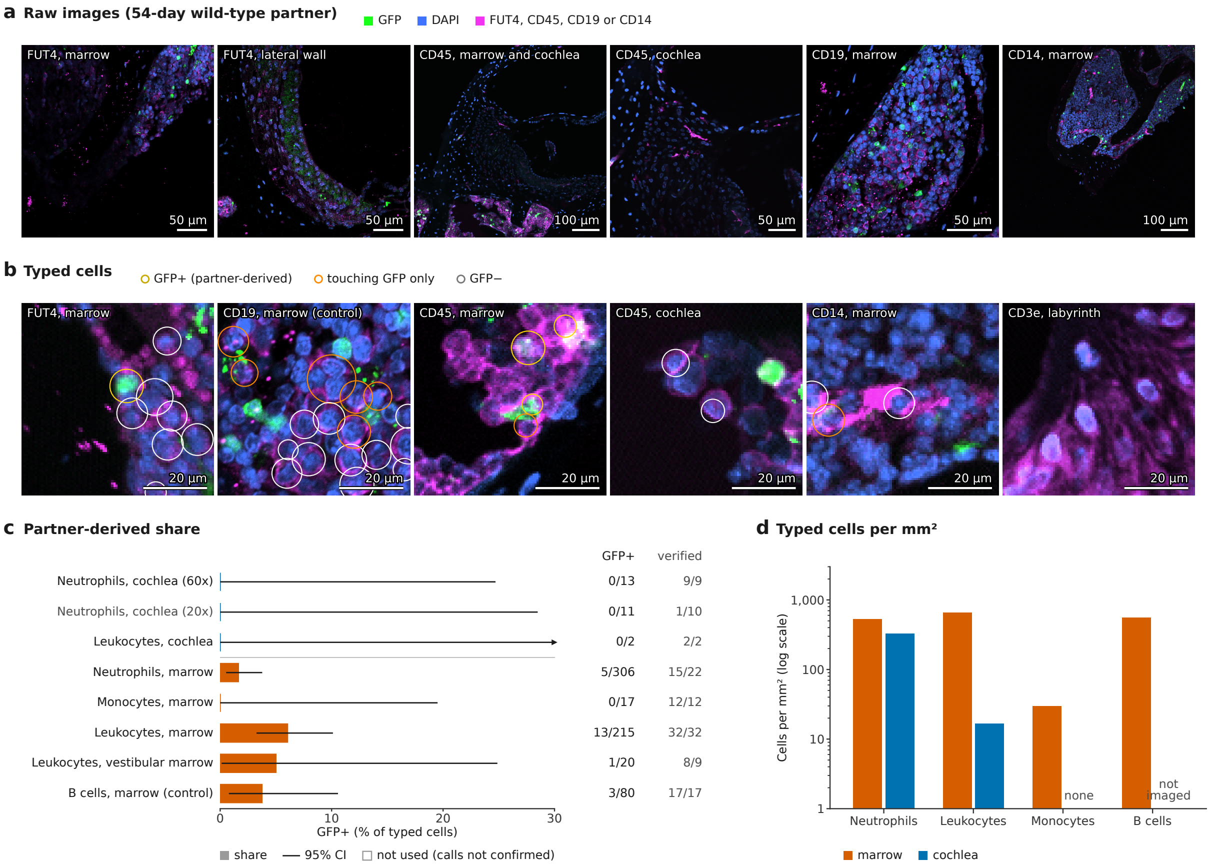


**Supplementary Fig. 6 | Immune cell types of the 54-day wild-type partner.** **a**, Raw images (one optical section each; marker, magenta; GFP, green; DAPI, blue): neutrophils (FUT4) in marrow and lateral wall, leukocytes (CD45) in marrow and cochlea, B cells (CD19) and monocytes (CD14) in marrow. **b**, Crops with the immune cells typed on segmented nuclei (a cell is the nucleus with a 1.5-$\mu$m ring; it is marker-positive when the ring is positive around the nucleus and at least twice its surroundings), circled by their GFP call: partner-derived (yellow), touching GFP only (orange) or GFP-negative (white). CD3e stained the labyrinth epithelium and is not interpreted. **c**, Partner-derived share per cell type and compartment (bars; lines, Clopper–Pearson 95% CI), with GFP$^{+}$ over typed cells and the typed cells confirmed as single marker-positive cells in the blinded review (verified). Cochlear neutrophils come from 60× sections of the lateral wall; the 20× cochlear FUT4 calls (1 of 10 confirmed) are shown in grey and not used. Reporter-negative B cells give the false-positive background (3 of 80). **d**, Typed cells per mm$^2$ of tissue per cell type in marrow and cochlea (cochlear neutrophils from the 60× lateral wall; no cochlear cell was typed CD14-positive, and CD19 was not imaged in the cochlea). Scale bars as labelled.

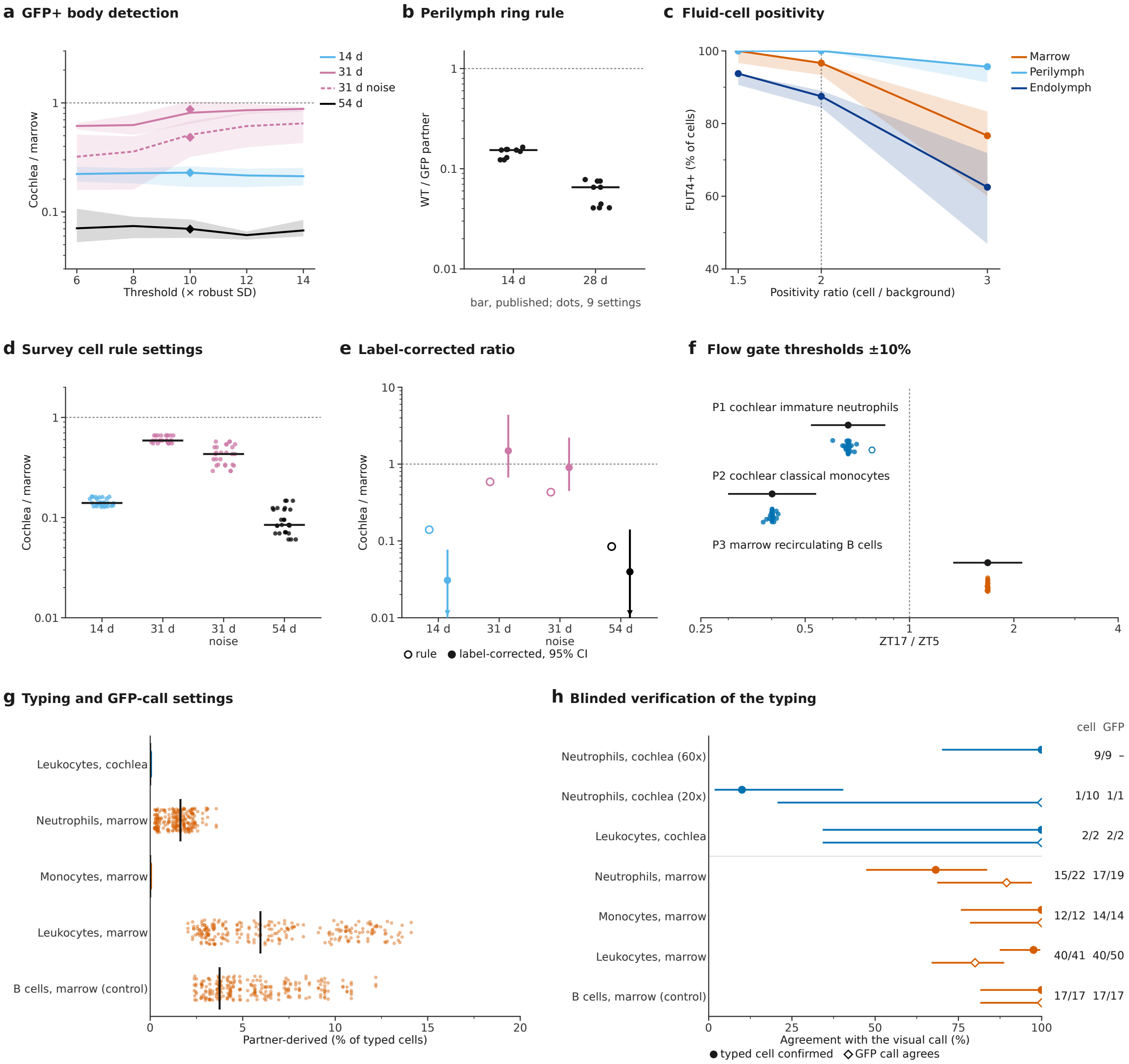


**Supplementary Fig. 7 | Robustness and blinded verification of the image and flow quantification.** **a**, $GFP^+$ body detection in the parabiosis survey: cochlea-to-marrow density per pair against the threshold (lines, median over area and solidity settings; bands, range; diamonds, published setting): below 1 in every unexposed pair at 44 of 45 settings. **b**, Perilymph ring rule: wild-type over GFP-partner $GFP^+$ share (points, 9 settings; bar, published): below 1 at all. **c**, Cochlear-fluid cells: $FUT4^+$ share against the positivity ratio (lines, median; bands, range over cell size and background annulus): perilymph at least 91% at every setting, endolymph 47–94%. **d**, Upgraded cell rule: cochlea-to-marrow ratio per pair at 27 settings of contrast, area and nucleus share (points; bar, published): below 1 in every pair at all 27 (0.06–0.66). **e**, Ratio corrected with the blinded labels (filled; bootstrap 95% CI; arrowheads, intervals reaching 0) beside the rule's ratio (open): noise pair 0.90 (0.45–2.21); others as in Supplementary Fig. 2e. **f**, Flow gates: ZT17/ZT5 ratio of each primary readout with each gate threshold moved by ±10% in turn or all fluorescence thresholds together (points; open, not significant after Holm correction) and with the published gates (black, 95% CI): replicated at all 31 settings, except cochlear immature neutrophils when all fluorescence thresholds fell together (Holm $P = 0.072$). **g**, Typing: partner-derived share per cell type at 243 settings (points; bar, published): no cochlear leukocyte partner-derived at any setting, marrow neutrophils below 5%, marrow leukocytes above the B-cell background at 159. **h**, Blinded verification of the typing (samples pooled): typed cells confirmed as single marker-positive cells (filled) and agreeing GFP calls of confirmed cells (open), Wilson 95% CIs; numbers, agreeing over decided items.

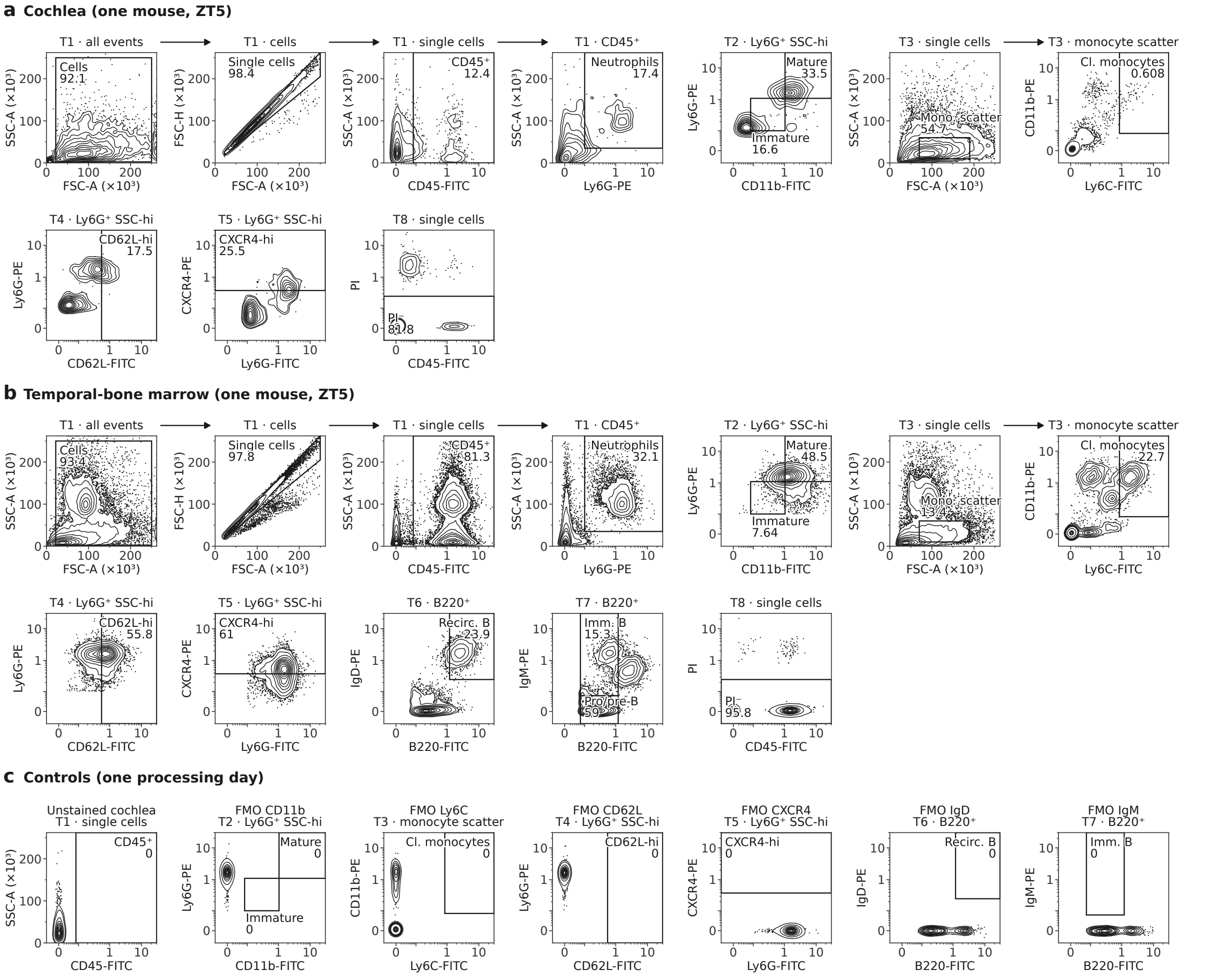


**Supplementary Fig. 8 | Flow-cytometry gating strategy and controls.** Contour plots with outliers (as in Fig. 5c) of one mouse of the validation cohort (ZT5, cohort 1; of the mice complete in both tissues, the one whose cochlear neutrophil share was closest to the median), with every gate and its frequency (% of the plotted parent population). Fluorescence in bead-relative units on an asinh scale; scatter linear ($\times 10^3$); gate thresholds in Supplementary Note 12. Arrows link successive gates of one tube; the other plots name their parent population. **a**, Cochlea: cells, single cells, CD45$^+$ cells and neutrophils (Ly6G$^+$ SSC$^{\text{hi}}$; tube T1); immature and mature neutrophils (CD11b versus Ly6G; T2); monocyte scatter and classical monocytes (Ly6C versus CD11b; T3); CD62L-hi (T4) and CXCR4-hi (T5) neutrophils; viability (propidium iodide; T8). The cochlear B-cell tubes (T6, T7) were run only when cells sufficed (3 and 6 of 20 cochleae) and feed no primary readout. **b**, Temporal-bone marrow of the same mouse, with recirculating B cells (B220$^{\text{hi}}$ IgD$^+$ of B220$^+$ cells; T6) and immature (B220$^{\text{int}}$ IgM$^+$) and pro/pre-B cells (B220$^{\text{lo/int}}$ IgM$^-$; T7). **c**, Controls of one processing day (marrow cells unless stated): unstained cochlea (CD45 gate) and the fluorescence-minus-one (FMO) control of every gated marker (CD11b, Ly6C, CD62L, CXCR4, IgD, IgM), drawn with the same gates. On all four processing days, no unstained or FMO control placed an event in its target gate (32 control–gate checks; 215–937 parent events each); one B220-bright, IgM-bright event in one IgM FMO (1 of 706 B220$^+$ events) lies outside the immature-B gate.

#### Supplementary Notes

Bench-level and parameter detail for the Methods and the pre-specified plan of the flow-cytometry cohort, numbered in the order in which the Methods cite them.

##### Supplementary Note 1 | Atlas samples, quality control and annotation

*Dissection.* After removal of the mandible and overlying skin, heads were immersed in ice-cold PBS and the brain, meninges and excess blood vessels were removed. Under a stereomicroscope, the endolymphatic sac was released and the cochlea was isolated with the semicircular canals preserved; the oval window was kept closed to limit the loss of labyrinthine fluids, vessels crossing the stapes footplate were cut at a distance from it, and facial-nerve fibres in the facial recess were removed together with the auditory-nerve fibres attached to the modiolus. In ice-cold MACS tissue storage solution, the cochlea was separated from the vestibule and its bony capsule fragmented: the cochlear soft tissue, stripped of residual bone, formed the cochlear sample, and the apical bone fragments containing marrow formed the marrow sample. Single-cell suspensions were prepared with the 10x Genomics tissue dissociation procedure, with 30 min at 37 °C for each digestion step, and filtered through a 40-$\mu$m Flowmi cell strainer; viability, measured by acridine orange/propidium iodide staining on an automated cell counter, was above 80%.

*Count matrices.* Cell Ranger 10.0.0 performed alignment, UMI collapsing and cell calling for the 10x libraries, and MobiVision 2.0 performed barcode and UMI identification, quality control and quantification for the MobiSeq libraries.

*Quality control.* Genes detected in fewer than three cells were removed, and the protein-coding and mitochondrially encoded genes of the GRCm39 autosomes and mitochondrial chromosome were selected by Ensembl BioMart coordinates retrieved with infercnvpy. Cells were retained with 200–7,000 detected genes, at least 500 UMIs and mitochondrial, haemoglobin and ribosomal transcript fractions below 5%, 10% and 40% (OmicVerse), and doublets were removed per library with Scrublet.

*Integration and annotation.* A batch-corrected latent space was learned with scVI (20 dimensions, conditioned on library) from 2,000 highly variable genes selected across libraries by the Seurat v3 method. Cells were clustered on a 15-nearest-neighbour graph of this space by landmark-based clustering (landmarks chosen by weighted $k$-medoids, cell memberships by random walk with restart, labels from the most probable landmark), and the marker genes of each cluster were derived by correlating expression with landmark memberships. Markers were confirmed by pseudobulk differential expression of each cell type against all other cells with PyDESeq2 (aggregated per library, accounting for batch; Benjamini–Hochberg FDR $< 0.05$ and $\log_2$ fold change $\geq 0.25$).

*Label audit (Extended Data Fig. 1).* The audit covered per-sample quality metrics, cells per subtype and sample, a marker-based label margin per cell (score for the own label minus the best alternative; score, mean $z$-scored expression of the label's marker genes) and the mixing of samples among same-type nearest neighbours.

#### Supplementary Note 2 | Benchmark of rhythm-aware embeddings

*Cell identity.* Conservation of cell identity was scored on label-stratified subsamples (30,000 cochlear, 20,000 marrow and 40,000 joint cells) by isolated-label score, $k$-means normalized mutual information and adjusted Rand index, label silhouette and cell-type LISI (scib-metrics), five-fold $k$-nearest-neighbour label-transfer balanced accuracy and the Spearman correlation of diffusion pseudotime with the stage order of the neutrophil and B-cell lineages.

*Time of day.* Time-of-day structure was scored within each tissue and cell type by two measures, each expressed relative to a 30-component principal component analysis (PCA) of the same genes, capped at 1 and then averaged: the variance of the embedding explained by the cosine and sine of ZT, and the excess share of same-sample (that is, same-time) cells among the same-type cells within each cell's 15 nearest neighbours over the share expected from sample sizes.

*Platform mixing.* The joint embedding was also scored for mixing of the two platforms in shared immune types (batch-removal-adapted silhouette, integration LISI, graph connectivity and principal-component regression).

*Features and models.* Feature sets (Seurat v3 highly variable genes with log-normalization, or analytic Pearson residuals; 3,000 or 5,000 genes; with or without mitochondrial, ribosomal, haemoglobin and stress genes) were screened first by PCA. Candidate models had 30 latent dimensions; scVI and scANVI had two hidden layers of 128 units and a negative-binomial likelihood, and sysVI used platform as system and sample as covariate. The two best models were refitted with 20, 30 and 50 latent dimensions and with two further seeds at the best size; scANVI was trained with the labels of the evaluation subsample masked.

*Selection.* The composite score was 0.7× identity + 0.3× time per tissue and 0.5× identity + 0.2× platform mixing + 0.3× time jointly (feature sets were ranked by the per-tissue weights after min–max scaling), and embeddings keeping less than 80% of the baseline time-of-day structure were rejected. None of the selected embeddings was conditioned on sample; they defined neighbourhood graphs with $k = 30$, and the unsupervised scVI embeddings used for diffusion pseudotime and trajectories had 30 dimensions.

#### Supplementary Note 3 | Rhythm model, single-sample criteria, calibration and simulation

*Model.* After an expression filter and TMM normalization, the limma–voom cosinor had the design $\sim \cos(\omega t) + \sin(\omega t)$, and rhythmicity was tested by the moderated $F$-test with 2 degrees of freedom after robust empirical Bayes.

*Single-sample dependence.* A rhythm collapsed when one time point was left out if the leave-one-time-point-out maximum $P > 0.25$ or the minimum amplitude ratio $< 0.5$.

*Calibration.* Dense public time series were subsampled to our design, with the held-out samples as truth: 12 organs in constant darkness sampled every 2 h (GSE54650; 480 draws) and sham-operated cochlea sampled every 4 h (GSE88954; 40 draws), with and without a sample-level nuisance shaped like our cochlear sample pattern added to 40% of genes.

*Simulation on our data.* For four cell types per tissue, samples were relabelled with a random non-identity class (removing true rhythms but keeping all sample-level structure), and rhythms of known fold change were injected into 5% of genes by binomial thinning.

#### Supplementary Note 4 | Clock alignment and comparison at matched depth

*Joint alignment.* In the alternating weighted circular means, each fit was weighted by its signal-to-noise ratio, and 95% intervals came from resampling genes.

*Single-cell clock phase.* Cells with at least 3 clock-gene UMIs were scored.

*Matched depth.* All counts of each structural cell were thinned by one factor (binomially for clock genes) so that each structural type matched the clock-transcript distribution of the cochlear haematopoietic cells (quantile matching; 20 replicates; with and without subsampling to their median number of cells); without thinning, the comparison was repeated among cells with 3–5 clock transcripts. By a rule fixed before the analysis, a structural type tracked time at haematopoietic depth if exact $P \leq 0.05$ in at least half of all replicates (replicates with fewer than 300 testable cells counted as not tracking), and the structural–haematopoietic contrast was judged power-limited unless at least half of the structural types tracked.

#### Supplementary Note 5 | Transcription-factor activity, promoter motifs and rhythm drivers

*Activity versions.* Besides the primary test, activities were tested depth-matched by binomial thinning and centred on the per-sample median across a tissue's cell types; the lag between activity and the factor's own mRNA phase was taken in $(-12, 12]$ h.

*Promoter motifs.* E-box, E′-box, D-box, RORE, GRE, CRE, heat-shock element and NF-$\kappa$B sites were counted on both strands of 1-kb upstream sequences (UCSC mm10 RefSeq) and tested for enrichment among rhythmic genes (Cochran–Mantel–Haenszel test stratified by expression decile) and for the phase of rhythmic carriers (directional permutation test towards the expected phase: E-box ZT8, D-box ZT12, CRE and GRE ZT13, heat-shock element ZT18, RORE ZT22).

*Driver classes.* The evidence flags were, in their order of precedence for the primary class, core clock, temperature, handling-linked, composition-linked, glucocorticoid-driven, clock-controlled and sample-pattern-linked.

*Constant darkness.* The test asked whether calls peaked within 4 h of the clock-aligned phase of the same gene in constant darkness (GSE54650; rhythmic in $\geq 2$ organs) more often than with peaks shuffled within each cell type (200 draws).

*Handling.* A per-sample and per-cell-type dissociation-stress score was split by least squares into cell-type and sample effects; a call was handling-linked if its profile followed its cell type's stress profile ($r > 0.8$).

*Glucocorticoids and heat shock.* A call was glucocorticoid-dependent with sham $P < 0.05$ and adrenalectomy $P > 0.2$ in the independent cochlear series with and without adrenalectomy (GSE88954). The night–day heat-shock check used the independent cochlear cohort without noise (GSE107086; sham surgery or adrenalectomy, about ZT17.5 versus ZT5.5).

#### Supplementary Note 6 | Composition models and neighbourhood abundance

*Frames and models.* Cell-type shares were tested by least squares on percentages of tissue cells (primary, as plotted) and on logits; by centred and additive log-ratios (structural and marrow-content references) and within-lineage fractions, each with the exact relabelling null; by propeller; by sccomp; by quasi-binomial and beta-binomial generalized linear models; and, for cochlear immune types, by the marrow-content decomposition below.

*Marrow-content decomposition (sensitivity analysis of cochlear immune shares).* For cochlear sample $s$ (sampling time $t_s$) and haematopoietic cell type $k$ (types with at least 5 cells in every cochlear sample; marrow-restricted precursors excluded), $y_{sk} = \log_2[(n_{sk} + 0.5)/(n_{\text{struct},s} + 0.5)]$, where $n_{\text{struct},s}$ counts the structural cells of the sample, was modelled as $y_{sk} \sim \mathcal{N}(\alpha_k + \beta_k z_s + a_k \cos \omega t_s + b_k \sin \omega t_s, \sigma_k)$, where $z_s$ is the centred $\log_2$ ratio of marrow-restricted precursors (haematopoietic stem and multipotent progenitor cells, granulocyte–monocyte progenitors, cycling pro-B cells, pre-B cells and erythroblasts) to structural cells. Priors: $\alpha_k \sim \mathcal{N}(0, 5)$; $\beta_k \sim \mathcal{N}(\mu_\beta, \tau_\beta)$ with $\mu_\beta \sim \mathcal{N}(1, 0.5)$ and $\tau_\beta \sim \text{HalfNormal}(0.5)$; $a_k, b_k \sim \mathcal{N}(0, \tau_r)$ with $\tau_r \sim \text{HalfNormal}(1)$; $\sigma_k \sim \text{HalfNormal}(\sigma_0)$ with $\sigma_0 \sim \text{HalfNormal}(1)$. Sampling used the No-U-Turn sampler in NumPyro (target acceptance 0.95; four chains of 2,000 warm-up and 2,000 retained draws). A second fit without $\beta_k$ served as the unadjusted comparison.

*sccomp.* Composition rhythms were also fitted with sccomp (Bayesian beta-binomial regression on cell counts with the composition formula $\sim \cos \omega t + \sin \omega t$), using the package's default priors and its outlier-removal step.

*Neighbourhood abundance.* The sparse re-implementation of Milo used the nearest-neighbour graph ($k = 30$) of the scANVI latent space, index cells refined from a random 5% (cochlea) or 10% (marrow) of cells, an edgeR quasi-likelihood negative-binomial cosinor with a 2-df $F$-test and a spatial false-discovery rate with $k$-distance weights. For the production waves, diffusion pseudotime was computed on the scVI latent space, rooted in granulocyte–monocyte progenitors for the neutrophil branch and in pro-B cells for the B lineage.

#### Supplementary Note 7 | Neutrophil-timer tests

*Concordance.* Spearman's $\rho$ between predicted and observed changes from ZT5 to ZT13 was computed for all shared genes, for genes with $P < 0.05$ in both data sets and for all genes except dissociation-stress and immediate-early genes, with gene-label permutation $P$ values (2,000 shuffles). Specificity rankings were computed for all, neutrophil-enriched and non-enriched genes.

*Cross-tissue split.* The cochlear minus the marrow correlation was compared with all 3,600 combinations of the two tissues' 60 relabelling classes (one-sided in the predicted direction; aged and pooled neutrophils specified before the analysis) and with all 720 × 720 permutations of the time labels, which include phase-shifted and reversed versions of the true labels.

*Scores.* The blood-timer score of a cell was the mean expression of the 50 genes most increased minus that of the 50 most decreased between ZT5 and ZT13 in GSE86619 ($|\log_2$ fold change$| \geq 0.5$, $P < 0.05$). The maturation score used the 50 genes most positively minus the 50 most negatively correlated with diffusion pseudotime in reference neutrophils (GSE137540).

#### Supplementary Note 8 | External data and the noise cohort

All external data were public (Supplementary Table 20). *External atlases.* Raw matrices were processed with emptyDrops (lower bound 100 UMIs, FDR $< 0.001$), followed by per-sample quality control ($\geq 200$ genes and $\geq 500$ UMIs; log UMIs and log genes within median $\pm 3$ median absolute deviations (MAD); mitochondrial share $\leq \min(25\%, \text{median} + 3\,\text{MAD})$) and Scrublet. A transferred label was confident with scANVI probability $\geq 0.7$ and $k$-nearest-neighbour vote $\geq 0.5$.

*Spatial deconvolution.* Rows were weighted by $1/\sqrt{\text{gene mean}}$ in the non-negative least-squares fit.

*Attribution of bulk rhythms.* Composition-only and regulation-only profiles held cell numbers or expression at their means. The null reassigned the external curves to random rhythmic genes (200 draws), and cell-type contributions were compared with expression-matched non-rhythmic genes by bootstrap (1,000 draws).

*Noise cohort.* Mice were kept on a 12-h light–dark cycle (lights on at 06:00) and exposed for 2 h to noise (2–20 kHz, 107 dB SPL) at ZT2–4 (day) or ZT14–16 (night). Single-cell libraries were made before noise (controls) and 3 days after noise (11–12 mice pooled per library), and bulk RNA-seq before and 1 day after noise ($n = 3$ per group). STARsolo used the 10x Genomics mm10-3.0.0 reference, and its filtered matrices (EmptyDrops-based cell calls) were analysed. Library phases were estimated by template matching against the independent sham-cochlea series (T2, primary), our cochlear fits (T1) and our structural cell types (T3, composition-robust), with noise-robust variants and 500 gene-bootstrap replicates.

#### Supplementary Note 9 | Imaging, spatial transcriptomics and parabiosis procedures

*Three-dimensional imaging.* Deeply anaesthetized mice received 50 $\mu$l of fluorescent lectin by slow tail-vein injection and were left under anaesthesia for 30 min to allow circulation and endothelial binding; they were then perfused transcardially with ice-cold PBS to remove unbound lectin, followed by 4% paraformaldehyde (PFA). Temporal bones were dissected and freed of muscle and brain under a stereomicroscope, the round and oval windows were opened with fine forceps and a small apical fenestra was made to let reagents into the scalae. Cochleae were post-fixed in 4% PFA overnight at 4 °C, washed in PBS and decalcified in 10% EDTA (pH 7.4), changed daily, until the otic capsule became compliant. They were permeabilized and blocked for 1–2 days in PBS with 1% Triton X-100, 5% normal serum and 1% bovine serum albumin, incubated with primary antibodies for 2–3 days and with fluorophore-conjugated secondary antibodies for 1–2 days, and washed in PBS. For cellular resolution, the modified ScaleS protocol was sorbitol-based, with refractive-index matching in antipyrine and 2,2′-thiodiethanol; cochleae were mounted in custom chambers and imaged as tiled z-stacks with 1-$\mu$m steps, and stitched volumes were used for reconstruction. Volumes were inspected, reoriented, cropped and converted in Fiji with the Bio-Formats importer and rendered in Imaris 11.0 (volume rendering, orthogonal slices and surfaces); displayed volumes are registered multichannel stacks without non-linear intensity remapping.

*Stereo-seq.* The mouse was decapitated after cervical dislocation, and its cochleae were dissected on ice after removal of the brain, meninges and blood. The round and oval windows were opened, and cochleae were fixed in RNase-free 10% neutral buffered formalin at 4 °C for 24 h, decalcified in EDTA at 4 °C for 48 h (changed daily), bisected along the modiolus, dehydrated through graded ethanol, infiltrated with paraffin for 1.5 h and embedded for sectioning. SAW 1.0 performed spatial barcode reconstruction, alignment and molecule-identifier correction. Tangram was run in cell-mapping mode, trained on the marker genes shared by the atlas and the spatial data, and the Cell2location estimate was the 5th percentile of the posterior abundance.

*Cochlear fluids and confocal images.* After gross cleaning, perilymph was aspirated first through the cochlear windows with a glass microcapillary; after brief washing and drying, the bone over the basal and middle turns was opened and endolymph was aspirated with a new microcapillary, and the apical bone fragments containing marrow were collected separately. Perilymph and endolymph were processed separately, and marrow cells were released from the apical bone fragments by gentle trituration in ice-cold PBS. Cells were deposited onto poly-L-lysine-coated glass slides by cytocentrifugation (300*g*, 5 min), fixed in 4% paraformaldehyde in PBS for 15 min at room temperature, washed three times in PBS and blocked in 3% bovine serum albumin in PBS for 1 h at room temperature. Slides were incubated overnight at 4 °C with the primary antibodies, washed three times for 5 min in PBS, incubated where required with fluorophore-conjugated secondary antibodies for 1 h at room temperature in the dark, washed three more times and coverslipped in antifade mounting medium containing DAPI. Unstained samples (autofluorescence), secondary-only controls (non-specific secondary binding), single-stained controls (bleed-through between channels) and marker-appropriate positive and negative biological controls were processed alongside and imaged with the same settings as the samples. Images were acquired on an Olympus FV3000 confocal microscope with a 60× objective. Pixel size and laser settings were read from each field's acquisition record. The local background of a cell was the median 3–8 µm from the nucleus. The fields acquired with the 561-nm line at 0% were the perilymph fields and endolymph fields 1–4. Burned-in scale annotations were replaced by vector scale bars of the same length.

*Parabiosis.* Under isoflurane anaesthesia (4% for induction and 2% for maintenance, in oxygen), fur was removed from the adjacent flanks. Longitudinal skin incisions were made on the right flank of the wild-type mouse and the left flank of the Lyz2-EGFP mouse; the dorsal skin and subcutaneous fascia were bluntly dissected to create a continuous shared subcutaneous space, the adjacent forelimbs were joined by a suture at the elbow, and the dorsal and ventral skin edges were joined by continuous sutures. Analgesia was standard-release buprenorphine (0.1 mg $kg^{-1}$ subcutaneously every 8 h for 72 h, starting before surgery) with carprofen (5 mg $kg^{-1}$ subcutaneously once daily for 3 days), and enrofloxacin (2.5 mg $kg^{-1}$ subcutaneously) was given perioperatively and once daily for 3 postoperative days. Mice recovered on a heating pad. The 60× confocal fields of the 54-day animal (Olympus FV3000) had 0.114 and 0.228 µm per pixel. $GFP^{+}$ cell bodies were detected on GFP smoothed at $\sigma = 0.5\,\mu m$ (above the tissue median plus 10 robust s.d., holes filled, $\geq 15\,\mu m^2$, solidity $\geq 0.7$); lateral-wall $FUT4^{+}$ cells were above the Otsu threshold of marrow FUT4, with $\geq 15\,\mu m^2$.

*Parabiosis survey (Supplementary Figs 2–4).* The complete image collection of the seven pairs was surveyed with rules fixed before analysis and extended once, before the figures were drawn, to include every usable acquisition. Native ZEISS CZI acquisitions (8-bit) were used; snapshots, projections and exports were not, except for the liver, which was available only as rendered exports. Included were acquisitions of the wild-type partners with a GFP channel, pixels of at most 0.70 µm and a tissue and duration label, or, for unlabelled fields of a labelled animal, a tissue assigned from morphology (haematopoietic marrow inside bone; cochlear lateral wall, duct or turn), shown with every analysed tissue in Supplementary Fig. 2a. Excluded were overviews, acquisitions whose 488-nm channel reproduced another label (Pearson $r \geq 0.8$ with another channel inside tissue, which removed the unlabelled 5-day phalloidin fields; every included field had $r \leq 0.75$) or carried phalloidin, fields without a localisable region, two 54-day fields of the ear bone that held no marrow, and repeated acquisitions of one position: within a session and specimen, two acquisitions whose fields overlapped by more than half with focus within 10 µm were counted once (the coarser, then the first). Tissue was the sum of the channels scaled to their 99.5th percentiles, smoothed ($\sigma = 6$ µm), above a quarter of its Otsu threshold, closed and hole-filled; z-stacks were sampled at sections at least 10 µm apart. GFP$^+$ bodies followed the rule above; in lectin acquisitions, bodies as bright for lectin as for GFP were flagged as broad-spectrum objects. Upgraded cell rule: a body counted as a partner-derived cell when its area was at least 25 µm$^2$ (the largest minimum area of the detection grid below), its mean GFP was at least twice the median GFP of the tissue 2–6 µm beyond it (the convention of the perilymph ring rule), it was not broad-spectrum and, in fields with DAPI, at least 10% of it lay inside a DAPI nucleus (the nucleus convention of the typing); these conventions were fixed before any count with the rule was inspected. The partner of unlabelled fields was inferred from the animal's marrow (GFP$^+$ area below 10% of tissue, wild type). Detection check: for each cochlear field, the share of the same animal's marrow GFP$^+$ bodies whose signal above the tissue median, scaled to the field's settings (linearly with 488-nm transmission and by a detector-gain factor measured on one marrow position imaged at two voltages with all else equal), would clear the field's threshold; the cochlear count divided by that share is the detection-corrected density (the gain factor was the signal ratio of 0.46 for 664 versus 700 V). Label correction: per pair and compartment, the estimated number of partner-derived cells was the number of kept bodies times the share of labelled kept bodies confirmed as cells, plus the number of rejected bodies times the share of labelled rejected bodies that were cells, from two blinded samples (one drawn within pair and compartment before the rule was set, one within pair, compartment and call afterwards; uncertain labels excluded); the 95% interval of the cochlea-to-marrow ratio came from 10,000 bootstrap draws of the labelled bodies within each stratum. Displayed GFP crops are linear from the section's tissue median to the larger of the crop's 99.9th percentile and the detection threshold; perilymph images show GFP with one range for both partners of a pair (the GFP partner's 0.5th–99.9th percentile) and DAPI per field. Depth check: the density in each acquisition's analysed section with the highest GFP 99.9th percentile within tissue. Perilymph cells: nuclei by Otsu thresholding of smoothed DAPI ($\sigma = 0.5$ µm) with watershed splitting (peaks at least 2.5 µm apart), at least 10 µm$^2$, edge-touching nuclei dropped; a cell was GFP$^+$ when the mean GFP of a 2- µm ring around its nucleus was at least twice the local background (median 3–8 µm from the nucleus outside every cell's 3- µm zone; background floored at 2 grey levels) and at least 5 grey levels above it. Noise comparison: the noise-exposed pair's pooled values were placed among the unexposed pairs on the log scale (z-score; 95% prediction interval, $t$ distribution with $n-1$ degrees of freedom); within animals, sections were resampled within each acquisition (10,000 bootstrap draws) for intervals of pooled densities and of the ratios to the unexposed 31-day pair (same duration) and to the unexposed cochlear apex (same region). The perilymph of the noise session was not interpreted, because its GFP partner's own perilymph cells were scored 7% GFP$^+$ (detector offset $-10$ in that session).

*Brain, brain-image format and stricter rule (Fig. 4g and Supplementary Fig. 5).* The brain exists only as a rendered orthogonal maximum projection of the 31-day wild-type partner (lectin, rendered magenta; GFP, rendered green; 8-bit RGB, 1,024 × 1,024 pixels; no DAPI, no pixel size; its source acquisition is not in the collection). Each rendering was read as the maximum over its colour channels and counted with the detection and upgraded rule above without the nucleus criterion, at 2.773 µm per pixel (21 overview acquisitions of the collection with 1,024-pixel frames; the field then spans 2.84 mm, consistent with the striatal fibre bundles it shows) and at the collection's other overview pixel sizes (1.387, 1.248 and 1.040 µm). For a like-for-like comparison, every wild-type tissue field was converted to the brain's format: each analysed section (or, as a check, the maximum projection of the whole stack) box-averaged to the target pixel size and rounded to 8-bit levels, DAPI used for the tissue mask only. Stricter rule: from a grid fixed before any count with it (GFP signal above the tissue median at least 10, 15, 20 or 30 robust s.d.; contrast to the 2–6- µm ring at least 2, 3, 4 or 6; area at least 25, 40 or 60 µm$^2$; nucleus share at least 10, 30 or 50%), the rule with the highest F1 against the 152 decided blinded labels of the two reviews (56 cells), ties broken by precision; out-of-sample performance by repeating the choice with each animal's labels left out. Native: 20 robust s.d., other criteria unchanged (precision 0.77 versus 0.57 for the upgraded rule; sensitivity 0.70 versus 0.79; left-out animals 0.72 and 0.68). Brain-image format (labelled bodies matched to converted bodies within 8 µm): every brain body reaches the 8-bit ceiling of the rendering (at most 10.7 robust s.d. above the tissue median), so a signal threshold above 10 robust s.d. would exclude every brain body whatever it is; after the full grid had done so, the signal criterion was held at detection and only contrast and area were tightened: contrast at least 6 (precision 0.64 versus 0.54, sensitivity 0.54 for both). The brain's detection threshold lies near that ceiling (241 of 255 grey levels), whereas the converted tissues are searched far more sensitively. A detector on the local background was therefore fixed before its first run (design and use rule recorded beforehand; the pre-run script archived with its MD5): GFP smoothed at $\sigma = 0.5$ µm, white top-hat with a 15- µm disk, bodies above the top-hat's tissue median plus 10 robust s.d. (at least 15 µm$^2$, solidity at least 0.7), called by the upgraded criteria or the stricter contrast (at least 6), and scored against the same labels; it was to replace the survey detector in the brain format if its F1 was higher. It was (strict call 0.61 versus 0.58; precision 0.65 versus 0.64, sensitivity 0.57 versus 0.54), and it raised the brain's count from 2.1 to 4.8 cells per mm$^2$ (upgraded criteria, 8.0 to 17.5) while leaving the cochlear counts nearly unchanged (31-day partner 18 to 17). Blood chimerism was taken from published parabiosis (Methods).

*Immune cell typing of the 54-day partner (Fig. 4i and Supplementary Fig. 6).* Fields of the ZEISS and Olympus series stained for FUT4, CD45, CD19, CD14 or CD3e (marker acquired with identical settings within a series) were typed on nuclei. Nuclei: DAPI smoothed at $\sigma = 0.5$ µm,

Otsu-thresholded within tissue and split by a distance-transform watershed (seeds at least 2.5 µm apart); nuclei of 10–150 µm$^2$ not touching the image border. A cell was its nucleus expanded by 1.5 µm within tissue, and its ring the cell minus the nucleus. A cell was marker-positive when at least half of its ring lay above the series threshold (Otsu of the marker smoothed at $\sigma = 0.3$ µm in the series' marrow tissue), the ring mean was at least twice the local floor (25th percentile of the marker 2–6 µm beyond the cell) and at least 5 of 8 angular ring sectors had 30% of their pixels above the threshold, so that an edge line passing one side of a cell did not count. A cell was GFP$^+$ when at least half of it lay inside GFP$^+$ bodies and its mean GFP was at least twice the median GFP 2–6 µm beyond it. Stains failing the separability or plausibility checks were reported, not interpreted (CD3e stained the labyrinth epithelium). The 13 FUT4$^+$ cells of the 60× lateral wall were typed as marker objects (above the Otsu threshold of marrow FUT4, at least 15 µm$^2$) and confirmed individually in the blinded review. Shares carry Clopper–Pearson 95% intervals.

*Blinded verification.* Samples were drawn with fixed seeds. Survey: a development sample of 137 bodies drawn within pair and compartment before the upgraded rule was set, and a validation sample of up to 6 kept and 4 rejected bodies per pair and compartment, excluding the development bodies. Typing: a development sample for the first nucleus-anchored rule and a validation sample for the final rule of up to 15 typed cells, 6 cells removed by the final rule and 4 untyped cells per marker and compartment, plus every partner-derived call, excluding the development cells. Crops showing GFP, marker and DAPI with the object outlined were coded and shuffled. The rater recorded whether the object was one marker-positive cell (or, for the survey, one partner-derived cell) and whether GFP clearly filled most of the outlined cell at the series display, with 'unsure' allowed and excluded from agreement; the visual assessments were performed by research personnel, blinded to call, compartment and file, and the labels were stored before the answer key was opened.

*Robustness grids (Supplementary Fig. 7).* Detection of GFP$^+$ bodies: threshold 6, 8, 10, 12 or 14 robust s.d., minimum area 10, 15 or 25 µm$^2$, solidity 0.6, 0.7 or 0.8. Upgraded cell rule: local contrast 1.5, 2 or 3, minimum area 15, 25 or 40 µm$^2$, nucleus share 0, 0.1 or 0.25. Typing: ring coverage 0.4, 0.5 or 0.6, marker contrast 1.5, 2 or 3, positive sectors 4, 5 or 6 of 8, GFP overlap 0.3, 0.5 or 0.7 and GFP contrast 1.5, 2 or 3. Cochlear-fluid cells: cell mask 1.0, 1.5 or 2.0 µm beyond the nucleus, background annulus 2–6, 3–8 or 5–10 µm, positivity ratio 1.5, 2 or 3. Perilymph ring: width 1.5, 2 or 3 µm, ratio 1.5, 2 or 3. Flow: every threshold of the gates of the three primary readouts moved by −10% and +10% in turn and all fluorescence thresholds moved together (31 settings), with the events re-gated from the cytometer exports and the pre-specified model refitted.

*Check of the vessel tracing (Supplementary Fig. 1).* The traced and the unannotated renders of each view were registered by phase cross-correlation (shift 0 pixels). Vessel ridges were detected on the vessel signal of the unannotated render (magenta minus green) with a Sato ridge filter (scales of 2–18 pixels) and an Otsu threshold within the rendered tissue. The tracing was skeletonized; thin traces (painted half-width at most 8 pixels, single vessels) were tested, and broad paint over wide vessels or bone was reported separately. The statistic was the share of thin-trace centreline within 2 pixels of a ridge, against 999 random rotations and translations of the same traces over the rendered tissue ($P = (1 + \text{draws at least as large as observed})/1{,}000$); scale from the burned-in 100- µm bars.

#### Supplementary Note 10 | Pre-specified analysis plan of the flow-cytometry validation cohort

The plan below (revision 2) was fixed on 15 October 2026, before any flow data were seen by the analysts. Its predictions, readouts, exclusions and statistics are reproduced without change; bench procedures suggested in the plan are omitted (the procedures used are in Methods and Supplementary Note 11). Expected values were computed only from the atlas (per-sample cell-type shares and the stored cosinor fits), without new analyses or outside data. Two caveats were stated in advance: absolute levels may differ between single-cell capture and flow cytometry, so the tested prediction is the direction and size of the ZT5-to-ZT17 change; and atlas effects may be overestimated (one pooled sample per time point), so the power analysis also considers half-size effects.

#### 1. Pre-specified readouts

| | Tissue | Readout (tubes) | Prediction | Atlas basis |
|---|---|---|---|---|
| P1 | Cochlea | Immature neutrophils ($Ly6G^{int}$ $CD11b^{int}$), % of $CD45^+$ (T1 × T2) | ZT5 > ZT17 | band neutrophils, robust; peak ZT5.2 |
| P2 | Cochlea | Classical monocytes ($CD11b^+$ $Ly6C^{hi}$), % of $CD45^+$ (T3 ÷ T1) | ZT5 > ZT17 | supported; peak ZT7.0 |
| P3 | Marrow | Recirculating B cells ($B220^{hi}$ $IgD^+$), % of $CD45^+$ (T6 ÷ T1) | ZT17 > ZT5 | robust; peak ZT20.5 |
| S1 | Both | CD62L on $Ly6G^+$ cells (% $CD62L^{hi}$, median) (T4) | cochlea ZT17 > ZT5; marrow flat | neutrophil timer split (robust); *Sell* rhythm |
| S2 | Both | CXCR4 on $Ly6G^+$ cells (% $CXCR4^{hi}$, median) (T5) | cochlea ZT5 > ZT17; marrow flat or opposite | timer split; *Cxcr4* / *Cxcr2* rhythm |

Other secondary readouts (descriptive or suggestive in the atlas): cochlear total neutrophils, $B220^+$ B lineage, recirculating and immature B cells and pro/pre-B cells; marrow immature neutrophils and classical monocytes; within-lineage fractions; absolute cells per cochlea; and the marrow content of cochlear samples (no rhythm expected; used as the sensitivity analysis for P1–P2).

#### 2. Expected results

Values are percentages of $CD45^+$ cells from the stored cosinor fits at ZT5 and ZT17, with the atlas samples on either side of each time point (ZT3 and ZT7 around ZT5; ZT15 and ZT19 around ZT17).

*Cochlea.*

| Gate (tube) | ZT5 | ZT17 | ZT17/ZT5 | Samples ZT3, ZT7 → ZT15, ZT19 |
|---|---|---|---|---|
| **Immature neutrophils (T1 × T2), P1** | **23%** | **7%** | **0.32** | 17, 27 → 9, 7 |
| Immature, % of Ly6G$^+$ (T2) | 64% | 38% | 0.59 | 67, 60 → 50, 33 |
| All neutrophils, Ly6G$^+$ (T1) | 35% | 19% | 0.54 | 25, 45 → 19, 22 |
| Mature neutrophils, Ly6G$^{hi}$ (T2) | 12.5% | 11.8% | 0.94 | 8, 18 → 9, 14 |
| **Classical monocytes (T3), P2** | **12.1%** | **5.5%** | **0.45** | 10.4, 13.0 → 7.0, 4.8 |
| B lineage, B220$^+$ (T6, T7) | 26% | 48% | 1.88 | 37, 16 → 41, 51 |
| Recirculating B (T6) | 1.1% | 4.1% | 3.65 | 1.5, 1.1 → 3.3, 4.5 |
| Immature B, B220$^{int}$ IgM$^+$ (T7) | 5.6% | 15.4% | 2.77 | 12.0, 1.2 → 12.0, 18.1 |
| Pro/pre-B, B220$^{lo}$ IgM$^-$ (T7) | 18.9% | 28.4% | 1.51 | 23.8, 13.7 → 26.2, 28.4 |

*Temporal-bone marrow.*

| Gate (tube) | ZT5 | ZT17 | ZT17/ZT5 | Samples ZT3, ZT7 → ZT15, ZT19 |
|---|---|---|---|---|
| **Recirculating B (T6), P3** | **9.6%** | **14.7%** | **1.53** | 11.5, 8.5 → 12.6, 16.3 |
| Recirculating, % of B220$^+$ (T6) | 32% | 41% | 1.26 | 33, 32 → 37, 43 |
| B lineage, B220$^+$ (T6, T7) | 30% | 36% | 1.21 | 35, 26 → 34, 38 |
| Immature neutrophils (T1 × T2) | 8.7% | 2.9% | 0.33 | 7.2, 10.4 → 3.8, 3.9 |
| All neutrophils, Ly6G$^+$ (T1) | 37% | 27% | 0.75 | 33, 39 → 28, 29 |
| Classical monocytes (T3) | 6.3% | 7.4% | 1.16 | 5.8, 7.0 → 7.6, 7.0 |

*Neutrophil surface markers (transcript-derived; lower confidence).* CD62L on Ly6G$^+$ cells was expected to rise from ZT5 to ZT17 in the cochlea (*Sell* peak ZT16.9; about twofold in transcripts) and not to change in marrow (*Sell* peak ZT23.7). CXCR4 was expected to be lower at ZT17 in the cochlea (*Cxcr4*/*Cxcr2* peak ZT6.2; *Cxcr4* 1.2–1.5-fold higher by day in neutrophil states) and flat or slightly higher in marrow. At the protein level, modest effects (about 1.2–2-fold in medians) with an opposite or absent change in marrow (a tissue × time interaction) were expected; absolute percentages positive could not be predicted.

#### 3. Tubes and controls

Eight two-colour tubes (FITC/PE): T1 CD45/Ly6G (CD45$^+$ denominator; neutrophils), T2 CD11b/Ly6G (immature versus mature neutrophils), T3 Ly6C/CD11b (classical monocytes in the monocyte scatter gate), T4 CD62L/Ly6G, T5 Ly6G/CXCR4, T6 B220/IgD (recirculating B cells), T7 B220/IgM (immature and pro/pre-B cells) and T8 CD45/propidium iodide (viability). Controls on each processing day: unstained cochlea and marrow; single-stain compensation controls; and fluorescence-minus-one controls for each gated boundary (CD11b, Ly6C, CD62L, CXCR4, IgD, IgM). A digestion control (marrow cells taken through the cochlear digestion, stained with T4 and T5 next to an undigested aliquot) measures how much digestion lowers CD62L and CXCR4; time points are therefore compared within a tissue, never marker levels between digested and undigested tissues. Cochlear cell-yield priority: T1, T2, T3 > T4, T5 > T6 > T7 > T8.

#### 4. Animals, groups and timing

Single sex, as in the atlas; light–dark 12:12 with lights on at 08:00 for at least 2 weeks; sampling at ZT5 (13:00) and ZT17 (01:00) within ±45 min, mice staggered by about 15 min; 10 mice per time point in two cohorts of 5 + 5, each containing both time points, with the order reversed in the second cohort; randomization by cage, coded sample identifiers and an analyst blind to ZT; dim red light (< 5 lux) at night until terminal anaesthesia; processing within 60 min of euthanasia with the same delay for every mouse.

#### 5. Analysis

*Gating.* One template for all samples with boundaries fixed on the controls before unblinding: cells (FSC-A versus SSC-A), single cells (FSC-H versus FSC-A), then the readout gates of each tube; boundaries for the T2 neutrophil split are drawn on marrow, where both clusters are visible, and applied to the cochlea.

*Calculations per mouse and tissue.* $CD45^+$ % of single cells (T1); neutrophils % of $CD45^+$ (T1); immature fraction of $Ly6G^+$ cells (T2); P1 = neutrophils % of $CD45^+$ × immature fraction; P2 = [$CD11b^+$ $Ly6C^{hi}$ monocyte-scatter cells / single cells] (T3) ÷ $CD45^+$ % of single cells (T1); P3 = [$B220^{hi}$ $IgD^+$ / single cells] (T6) ÷ $CD45^+$ % of single cells (T1); S1 and S2, the percentages of $CD62L^{hi}$ and $CXCR4^{hi}$ $Ly6G^+$ cells and the medians (T4, T5); marrow-content index of the cochlea = [$B220^{lo}$ $IgM^-$ / single cells] (T7) ÷ [$CD45^-$ / single cells] (T1). Consistency checks: $CD11b^+$ % of single cells within 20% (relative) between T2 and T3, and $B220^+$ between T6 and T7; failing samples are flagged and results reported with and without them.

*Exclusions (fixed in advance).* A cochlear tube with fewer than 3,000 single-cell events, or a readout's parent gate with fewer than 100 events, is excluded for that readout only; samples with viability below 50% are excluded; partially perfused samples are analysed with and without.

*Statistics.* P1–P3: linear model on log(% of $CD45^+$) with time and cohort; ZT17/ZT5 ratio with 95% CI; two-sided tests with Holm correction across the three primaries. S1–S2: linear mixed model on logit(% positive) and on log(median) with tissue × time, cohort and a random intercept per mouse; the interaction and the within-tissue contrasts are reported. Sensitivity: P1–P2 divided by the marrow-content index; unadjusted shares remain primary. Verdict per prediction: replicated, Holm $P < 0.05$ in the predicted direction; direction-consistent, predicted direction but not significant; contradicted, significant in the opposite direction.

*Power.* For the Holm threshold of the three primaries ($\alpha = 0.0167$, two-sided, log scale) and an assumed between-mouse coefficient of variation of 0.5 (cochlea) and 0.3 (marrow), 10 mice per time point give 99%, 86% and 72% power for P1, P2 and P3 at the atlas effect sizes (3.1-, 2.2- and 1.5-fold), and 54%, 26% and 18% at half these effects.

##### 6. Outcome

The data were analysed as pre-specified (Methods; Figs 5c and 6h and Supplementary Fig. 8). All three primary predictions were replicated: P1, ZT17/ZT5 ratio 0.67 (95% CI 0.52–0.85; expected 0.32); P2, 0.40 (0.30–0.53; expected 0.45); P3, 1.68 (1.34–2.10; expected 1.53). For the secondary readouts, the CXCR4 prediction was met (cochlea lower at ZT17, marrow unchanged; tissue × time $P < 0.001$) and the CD62L prediction was not (cochlea odds ratio 1.20, $P = 0.35$). One cochlea (ZT5) was excluded from P1, P2 and S2 by the event floor; no sample failed the viability rule.

#### Supplementary Note 11 | Flow-cytometry protocol

*Tissue.* Mice were perfused transcardially with 20–25 ml of ice-cold PBS without calcium and magnesium, and tissues were collected on ice under a dissection microscope. The cochlear preparation excluded the vestibule. Both cochleae of a mouse were opened, minced and digested in 0.5 ml of HBSS with calcium and magnesium containing collagenase type IV (1 mg ml$^{-1}$) and DNase I (0.1 mg ml$^{-1}$) for 25 min at 37 °C with gentle agitation and pipetting at fixed intervals. Digestion was stopped with 2 ml of cold FACS buffer (PBS with 2% heat-inactivated fetal bovine serum and 2 mM EDTA); cells were filtered through a 70-$\mu$m strainer rinsed with 2 ml of buffer, pelleted (400$g$, 5 min, 4 °C) and resuspended in 300 $\mu$l. Marrow was released mechanically into 1 ml of cold FACS buffer, filtered (70 $\mu$m), pelleted, treated with 1 ml of ACK lysis buffer for 2 min, diluted with 4 ml of buffer, pelleted and resuspended in 500 $\mu$l. Total and dye-excluding cells were counted in two 10-$\mu$l aliquots of each suspension mixed 1:1 with trypan blue, and the counted volume was deducted before allocation. Panel aliquots were allocated by fixed priorities (T3, then T1, the unfixed T8 aliquot and the controls); T6 and T7 were omitted when the recovered cells could not supply them, and the reason was recorded.

*Staining.* Cells were kept at 4 °C and protected from light. Fc receptors were blocked with BD Pharmingen Mouse BD Fc Block (unconjugated anti-CD16/32, clone 2.4G2; catalogue no. 553142; RRID AB_394656; about 0.5 $\mu$g per tube) for 10 min on ice; 50 $\mu$l of a twofold-concentrated antibody mixture was added to 50 $\mu$l of blocked cells (100 $\mu$l final; each antibody at 1:100), and cells were stained for 25 min at 4 °C and washed twice with cold FACS buffer. All staining antibodies were from BioLegend (Table 1 below). The eight two-colour panels (FITC/PE) were T1 CD45/Ly6G, T2 CD11b/Ly6G, T3 Ly6C/CD11b, T4 CD62L/Ly6G, T5 Ly6G/CXCR4, T6 B220/IgD, T7 B220/IgM and T8 CD45/propidium iodide.

**Table 1 | Flow-cytometry antibodies** (BioLegend; all at 1:100)

| Tubes | Antibody | Clone | Catalogue no. | RRID |
|---|---|---|---|---|
| T1, T8 | CD45–FITC | 30-F11 | 103108 | AB_312973 |
| T1, T2, T4 | Ly6G–PE | 1A8 | 127608 | AB_1186099 |
| T2 | CD11b–FITC | M1/70 | 101206 | AB_312789 |
| T3 | Ly6C–FITC | HK1.4 | 128006 | AB_1186135 |
| T3 | CD11b–PE | M1/70 | 101208 | AB_312791 |
| T4 | CD62L–FITC | MEL-14 | 104406 | AB_313093 |
| T5 | Ly6G–FITC | 1A8 | 127606 | AB_1236494 |
| T5 | CXCR4 (CD184)–PE | L276F12 | 146506 | AB_2562783 |
| T6, T7 | B220–FITC | RA3-6B2 | 103206 | AB_312991 |
| T6 | IgD–PE | 11-26c.2a | 405706 | AB_315028 |
| T7 | IgM–PE | RMM-1 | 406508 | AB_315058 |

Tubes T1–T7 were fixed in 1% paraformaldehyde for 15 min at 4 °C, washed and stored cold and dark until acquisition. T8 was kept unfixed, and propidium iodide (1 $\mu$g ml$^{-1}$ final) was added shortly before acquisition; T8 reports the viability of its own aliquot only.

*Controls.* On each of the four processing days, controls comprised unstained cochlear and marrow suspensions, single-stained compensation controls, fluorescence-minus-one controls for T2–T7, marrow reference tubes for T2, T3 and T6, and digestion controls for T4 and T5 (marrow cells taken through the cochlear digestion beside undigested marrow).

*Acquisition.* Data were acquired on a BD FACSCelesta with BD FACSDiva 9.0, at low sample pressure and with one acquisition template, detector settings and trigger threshold. FITC was excited at 488 nm and detected through a 530/30-nm filter, PE was excited at 561 nm and detected through a 586/15-nm filter, and propidium iodide was excited at 488 nm and detected through a 695/40-nm filter. Instrument performance was checked daily with BD FACSDiva CS&T Research Beads (catalogue no. 655050). Compensation for FITC and PE was set with BD CompBeads Anti-Rat Ig$\kappa$ (catalogue no. 552844) stained with each antibody, and for propidium iodide with a stained-cell control. Fluorescence intensities were expressed relative to BD SPHERO Rainbow Calibration Particles (8 peaks; reference = 1; catalogue no. 559123), which also monitored intensity across reagent lots; these bead-relative units are dimensionless ratios rather than calibrated molecule counts. Fixed tubes of both time points were acquired in a common session within each cohort, with balanced order, and T8 tubes fresh near each collection window. FSC-A, FSC-H, SSC-A, both fluorescence areas and time were recorded; time traces were inspected for flow interruptions; the recoverable volume of each tube was acquired without extending acquisition to reach a target count; and acquired, single-cell, parent and target events were recorded for every tube.

#### Supplementary Note 12 | Flow-cytometry gates, readouts and atlas labels

*Gates.* Fixed gates were applied to every sample in this order: cells (FSC-A versus SSC-A), single cells (FSC-H versus FSC-A), CD45$^+$ cells and Ly6G$^+$ SSC$^{hi}$ neutrophils, then the readout gates. Thresholds (fluorescence in bead-relative units, calibration beads = 1; scatter in channel units): cells, FSC-A 22,000–250,000 and SSC-A 2,500–250,000; single cells, FSC-H/FSC-A 0.82–1.18; CD45$^+$, $\geq$ 0.08; Ly6G$^+$, $\geq$ 0.1 with SSC-A $\geq 35,000$; immature neutrophils, CD11b

0.08–1.05 and Ly6G $< 1.1$; mature, CD11b $\geq 1.05$ and Ly6G $\geq 1.1$; monocyte scatter, FSC-A 70,000–190,000 and SSC-A 10,000–60,000; classical monocytes, CD11b $\geq 0.08$ and Ly6C $\geq 0.85$; CD62L-hi, $\geq 0.555$; CXCR4-hi, $\geq 0.380$; B220$^+$, $\geq 0.07$; recirculating B, B220 $\geq 1.2$ and IgD $\geq 0.25$; immature B, B220 0.07–1.2 and IgM $\geq 0.07$; pro/pre-B, B220 0.07–1.2 and IgM $< 0.07$; viable, propidium iodide $< 0.25$. Plots are contour plots with outliers on an asinh scale.

*Readouts.* P1 was the Ly6G$^+$ share of CD45$^+$ cells (T1) $\times$ the Ly6G$^{int}$ CD11b$^{int}$ fraction of Ly6G$^+$ cells (T2); P2, the CD11b$^+$ Ly6C$^{hi}$ cells in the monocyte scatter gate (T3) divided by the CD45$^+$ share of single cells (T1); and P3, the B220$^{hi}$ IgD$^+$ cells (T6) divided by the CD45$^+$ share (T1). P2 and P3 combine separately stained aliquots, so they are not bounded by 100% and were not truncated; zero denominators were treated as undefined. The secondary readouts came from T4 (S1) and T5 (S2) and were modelled on logit percentage and log median. The consistency check compared tubes T2 and T3, and the marrow-content index of the sensitivity analysis was B220$^{lo}$ IgM$^-$ cells (T7) over CD45$^-$ cells (T1), computed where T7 was available.

*Atlas labels.* The two-colour gates of the flow cohort map to the atlas labels as follows (Supplementary Fig. 8): Ly6G$^{int}$ CD11b$^{int}$ neutrophils correspond to the band and 'mature' neutrophil labels of the atlas ('mature' neutrophils being transcriptionally immature); Ly6G$^{hi}$ neutrophils to the 'aged' and activated labels; B220$^{hi}$ IgD$^+$ cells to naive (recirculating) B cells; B220$^{int}$ IgM$^+$ cells to the 'mature' (immature) B label; and B220$^{lo}$ IgM$^-$ cells to pro-B, transitional (large pre-B) and pre-B cells. The last gate also contains plasmacytoid dendritic cells (B220$^{int}$ IgM$^-$; about 1–2% of CD45$^+$ cells in the atlas). Flow phenotypes are named by their markers throughout.

#### Supplementary Note 13 | Evidence grading of positive findings

Every positive finding of the atlas was scored on pre-specified evidence dimensions. Each dimension received a status (supports, partial, against, neutral or not tested) computed from the result tables, with the numbers behind it recorded.

*Core dimensions.* Model concordance (agreement of independent rhythm or composition models) and resampling (exact time-relabelling permutation, leave-one-time-point-out refits, bootstrap) or calibrated false-discovery rate.

*Independent dimensions.* Reference frame (for compositional claims: shares of tissue, log-ratio and within-lineage frames), annotation confidence, embedding or sub-state, the tissue-wide sample-level pattern, molecular or mechanistic corroboration, cross-tissue logic, external data, literature, power or effect size and, for the immune timetable, the independent flow-cytometry cohort (replicated, supports; significant in the opposite direction, against; otherwise neutral). Cochlear immune rhythms were analysed within the cochlea; frames that express cochlear shares relative to marrow content were reported as sensitivity analyses and not counted.

*Verdicts.* Rules were fixed before the outputs were inspected and evaluated in this order:

- robust: all core dimensions support, at least two independent dimensions support and at most one is against;
- supported: no core dimension against, at least one core and one independent dimension support, and independent supports outnumber independent againsts;

- not supported: no core dimension supports and independent againsts are at least as many as independent supports;
- suggestive: everything else.

In the text, robust and supported findings are stated declaratively, suggestive findings with "suggests", and exploratory analyses (the composition and clock comparisons of the independent noise cohort) with their conditions.

*Pre-specified hypotheses.* The 12 B-lineage and 9 myeloid hypotheses (Extended Data Figs. 8c and 9a) were decided by their own rules: supported only with Benjamini–Hochberg $q < 0.1$ across the primary set and at least two agreeing independent methods (for myeloid hypotheses, also the pre-specified direction); partial when one criterion was met.

#### Supplementary Note 14 | Software versions

Atlas analyses used Python 3.12.3 (NumPy 2.4.6, pandas 2.3.3, SciPy 1.18.1, statsmodels 0.15.0, Scanpy 1.12.4 with its Scrublet implementation, scvi-tools 1.5.1 (including sysVI), scib-metrics 0.6.1, harmonypy 2.1.0, Scanorama 1.7.4, BBKNN 1.6.0, decoupler 2.2.0, NumPyro 0.22.0) and R 4.5.3 (limma 3.66.0, edgeR 4.8.2, MetaCycle 1.2.1, rain 1.44.0, sccomp 2.2.0, speckle 1.10.0, DropletUtils 1.30.0); trajectory checks of the pre-specified myeloid hypotheses used CellRank 2.3.3 and Palantir 1.4.5. Count matrices, quality control, integration and annotation used Cell Ranger 10.0.0, MobiVision 2.0 and Python 3.13 (AnnData 0.12.10, Scanpy 1.12, OmicVerse 1.7.9, Scrublet 0.2.3, scvi-tools 1.4.2, infercnvpy 0.6.1, PyDESeq2 0.5.4); spatial transcriptomics used SAW 1.0, Tangram 1.0.4 and Cell2location 0.1.5; and three-dimensional images used Fiji (release 2026-03-07) and Imaris 11.0. Flow data were acquired with BD FACSDiva 9.0 and gated in FlowJo 11.2 and custom Python code. Image analysis of confocal fields, flow cytometry and figures used Python 3.10.14 (scikit-image 0.25.2, tifffile 2025.5.10, SciPy 1.15.3, statsmodels 0.14.6, Matplotlib 3.10.0). The full list is in the code repository.

#### Supplementary Tables

Supplementary Tables are provided as separate files with a data dictionary of every column.

**Supplementary Table 1** Cell-type labels of the atlas: full names, abbreviations used in the figures, class and cells per tissue and time point.

**Supplementary Table 2** Cell-type summary: cells per sample, rhythm calls per tier, share of rhythmic genes, clock metrics, composition rhythm and power.

**Supplementary Table 3** Gene rhythms per tissue and cell type (cosinor fits, exact permutation statistics, independent tests and evidence tier).

**Supplementary Table 4** Core-clock gene fits and clock offsets per cell type.

**Supplementary Table 5** Composition rhythms of every cell type in every frame and model, with the multi-model verdict.

**Supplementary Table 6** Rhythms of hearing-related genes per cell type, with peak and trough windows.

**Supplementary Tables 7–8** Flow-cytometry cohort: per-mouse readouts and all pre-specified tests.

**Supplementary Table 9** Evidence grading of every positive finding: statement, status counts of the evidence dimensions, verdict and caveats; its finding identifiers are those of Supplementary Table 8.

**Supplementary Tables 10–13** B lineage: stage composition, ImmGen mapping, timeline and the 12 pre-specified hypotheses.

**Supplementary Tables 14–19** Myeloid lineage: composition, multi-method results, reference and neutrophil-map signatures, timeline and the nine pre-specified hypotheses.

**Supplementary Table 20** Public data sets analysed in this study: accession, species, tissue, assay, design, use and reference.
